# ANXA11 biomolecular condensates facilitate protein-lipid phase coupling on lysosomal membranes

**DOI:** 10.1101/2023.03.22.533832

**Authors:** Jonathon Nixon-Abell, Francesco S. Ruggeri, Seema Qamar, Therese W. Herling, Magdalena A. Czekalska, Yi Shen, Guozhen Wang, Christopher King, Michael S. Fernandopulle, Tomas Sneideris, Joseph L. Watson, Visakh V.S. Pillai, William Meadows, James W. Henderson, Joseph E. Chambers, Jane L. Wagstaff, Sioned H. Williams, Helena Coyle, Yuqian Lu, Shuyuan Zhang, Stefan J. Marciniak, Stefan M.V. Freund, Emmanuel Derivery, Michael E. Ward, Michele Vendruscolo, Tuomas P.J. Knowles, Peter St George-Hyslop

## Abstract

Phase transitions of cellular proteins and lipids play a key role in governing the organisation and coordination of intracellular biology. The frequent juxtaposition of proteinaceous biomolecular condensates to cellular membranes raises the intriguing prospect that phase transitions in proteins and lipids could be co-regulated. Here we investigate this possibility in the ribonucleoprotein (RNP) granule-ANXA11-lysosome ensemble, where ANXA11 tethers RNP granule condensates to lysosomal membranes to enable their co-trafficking. We show that changes to the protein phase state within this system, driven by the low complexity ANXA11 N-terminus, induce a coupled phase state change in the lipids of the underlying membrane. We identify the ANXA11 interacting proteins ALG2 and CALC as potent regulators of ANXA11-based phase coupling and demonstrate their influence on the nanomechanical properties of the ANXA11-lysosome ensemble and its capacity to engage RNP granules. The phenomenon of protein-lipid phase coupling we observe within this system offers an important template to understand the numerous other examples across the cell whereby biomolecular condensates closely juxtapose cell membranes.

**GRAPHICAL ABSTRACT:** 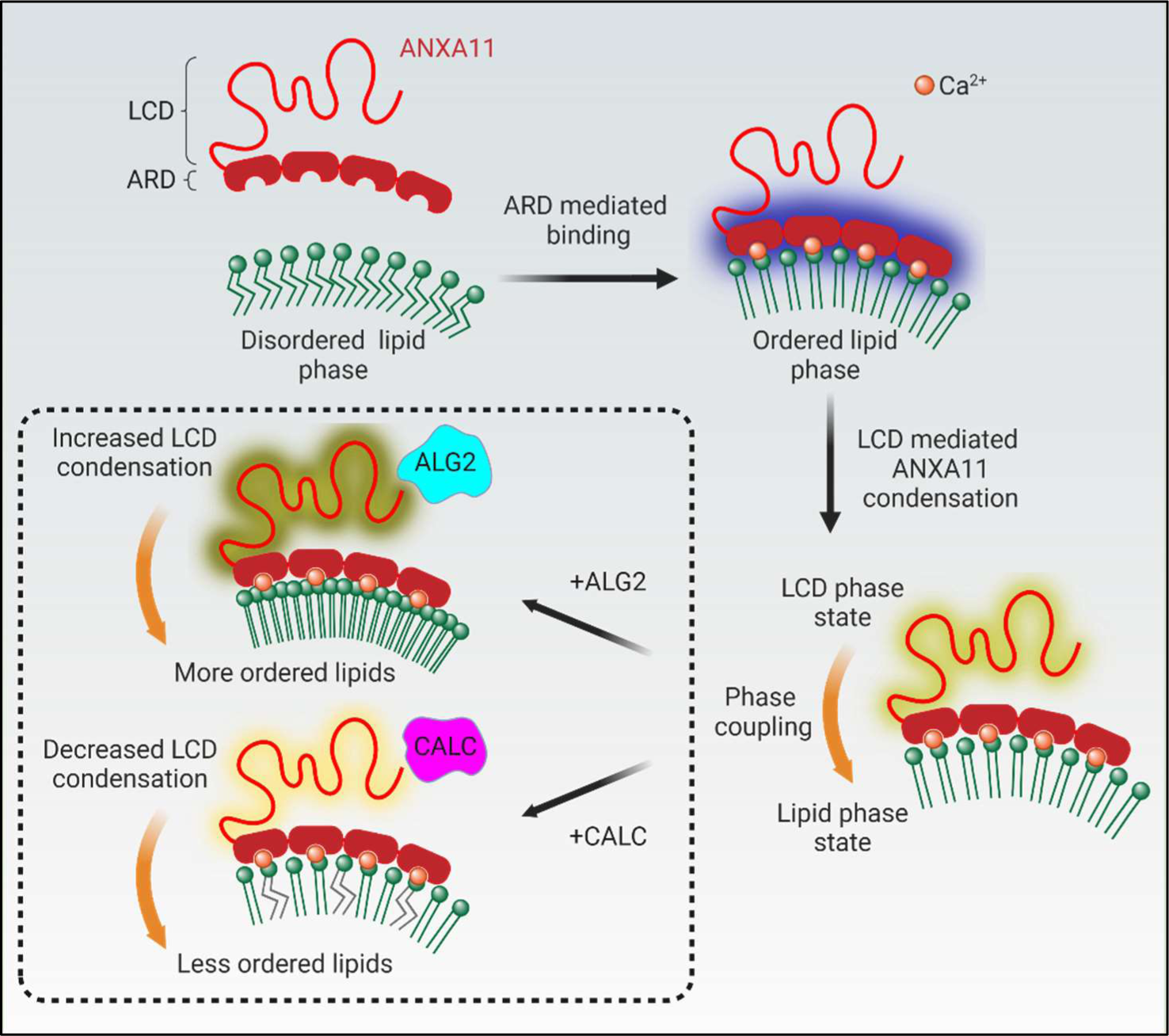

## INTRODUCTION

Intracellular phase transitions of both protein and lipid components have emerged as key physical principles affecting the material properties and organisation of various biological systems. Moreover, across eukaryotic cells there are now numerous examples of protein phase transitions occurring in close spatial proximity to lipid membranes. Despite this, the intriguing possibility that phase transitions in proteins and lipids could be functionally coupled remains to be thoroughly explored.

Phase transitions of cellular lipids have been an intense area of study dating back several decades (Eggeling et al., 2009; Heberle and Feigenson, 2011; Shaikh et al., 2001; Simons and Ikonen, 1997). The field has primarily focussed on the transition between disordered and ordered lipid phase states, which appears to be the most biologically relevant. In the disordered phase, lipids are highly fluid and are free to laterally diffuse within the membrane, whereas in the ordered phase, lipids become less mobile and more tightly packed. These changes in lipid mobility and crowding have been shown to profoundly impact the mechanical stiffness and nanodomain architecture of biological membranes (Levental et al., 2020). Such effects work in conjunction with protein-protein and protein-lipid interactions between structured proteins on or within the membrane to shape diverse biological processes, ranging from transmembrane signal transduction to viral budding (Aman and Ravichandran, 2000; Scheiffele et al., 1999).

Recent work has also led to the recognition of the important role played by protein phase transitions in the formation of biomolecular condensates. Biomolecular condensates (henceforth condensates) are heteromeric assemblies of proteins and/or nucleic acids that form intracellular membraneless compartments. Their assembly occurs as constituent proteins/nucleic acids undergo phase transitions from a dispersed/dilute phase to a condensed phase. Numerous functionally distinct condensates have now been identified across eukaryotic cells, playing important roles in diverse cellular processes ranging from transcription to intracellular signalling (Banani et al., 2017). A subset of these condensates forms on, or near, cellular membranes. Specific examples include: the T-cell receptor (TCR); the postsynaptic density 95 (PSD95) complex; synapsin-associated presynaptic vesicles; TIS granules and Sec bodies on the surface of the ER; P bodies on mitochondria; and recently the ribonucleoprotein (RNP) granule-ANXA11-lysosome complex. These membrane-associated condensates are involved in an assortment of cellular functions spanning transmembrane signalling, intracellular vesicle dynamics, and cargo trafficking (Huang et al., 2011; Lee et al., 2020; Liao et al., 2019; Ma and Mayr, 2018; Milovanovic et al., 2018; Su et al., 2016; Zacharogianni et al., 2014; Zeng et al., 2018; Zhang and Rabouille, 2019).

The juxtapositioning of phase transitioning proteins and membrane lipids within these systems raises the possibility that they could modulate each other’s phase state. For example, a lipid phase transition might influence the local accumulation, clustering, and phase state of protein components in adjoining biomolecular condensates. Conversely, protein components directly apposed to membranes could alter the local composition, ordering and phase state of subjacent lipids. Theoretical studies have suggested such a phenomenon might occur (Andes-Koback and Keating, 2011; Nesterov et al., 2021; Rouches et al., 2021). However, direct evidence for such phase coupling has been restricted to interactions between condensates and the plasma membrane (Chung et al., 2021; Lee et al., 2019; Wang et al., 2022).

Here, we investigate phase coupling within a previously described functional ensemble involving a cytoplasmic biomolecular condensate (RNP Granule) and a membrane-bound intracellular organelle (the lysosome). We have shown previously that the components of this ensemble are tethered together by ANXA11 to enable the long distance co-trafficking of RNP granules with lysosomes (Liao et al., 2019). ANXA11 engages the RNP granule via a low complexity N-terminal domain. At its C-terminus, ANXA11 contains a structured annexin repeat domain which binds to negatively charged phospholipid headgroups in the lysosomal membrane in a calcium-dependent manner.

In this study, we demonstrate that ANXA11 binding to membranes induces coupled changes in the phase state of the subjacent lipids, as has been reported for other annexin family members (Domon et al., 2012; Lin et al., 2020; Menke et al., 2005). However, we show that the magnitude of this lipid phase transition can be tuned according to the phase state of the ANXA11 low complexity domain. We subsequently demonstrate that two known protein interactors of ANXA11, apoptosis-linked gene 2 (ALG2; *PDCD6* (Satoh et al., 2002)) and calcyclin (CALC; *S100A6* (Watanabe et al., 1993)), modulate the phase state of the ANXA11 low complexity domain. We observe that ALG2- and CALC-mediated changes in the ANXA11 phase state evoke a coupled phase state change in the underlying lysosomal lipid membrane. A functional consequence of this phase coupling is the modulation of the mechanical properties of the ANXA11-lysosome ensemble, which correlates with its capacity to engage RNP granules. Together, these findings support the notion that the interaction of biomolecular condensates with intracellular membrane bound organelles can result in coupled changes in their biophysical properties and functions. As such, these observations also provide a framework for understanding how lipid and protein phase coupling may occur in other membrane-associated condensates across the cell.

## RESULTS

### The ANXA11 LCD drives a dispersed to condensed ANXA11 phase transition, while the ARD facilitates lipid binding

To explore the possibility of phase coupling in the RNP granule-ANXA11-lysosome ensemble, we built an *in vitro* reconstitution model comprising recombinantly purified ANXA11 and giant unilamellar vesicles (GUVs) composed of lysosomal membrane lipids (see STAR Methods).

The full length ANXA11 protein (aa 1-502; henceforth **FL**) comprises a mostly disordered N-terminal low complexity domain (aa 1-185; henceforth **LCD**), and a structured C-terminal annexin repeat domain (aa 186-502; henceforth **ARD**) (Figure 1A, Figure S1A-B). First, to explore the contributions of the LCD and ARD in condensate formation in solution, we incubated fluorescently labelled recombinant FL, LCD or ARD (Figure S4A) at physiological pH and temperature conditions. Supporting our previous observations (Liao et al., 2019), the LCD is both necessary and sufficient for inducing a concentration-dependent ANXA11 phase transition from a dispersed to a condensed state (Figure 1B, Movie 1). Importantly, this transition is reversible (Figure S1C-D) and does not elicit secondary structural changes consistent with protein aggregation (Figure S1E) (Qamar et al., 2018; Ruggeri et al., 2015a).

**Figure 1.**
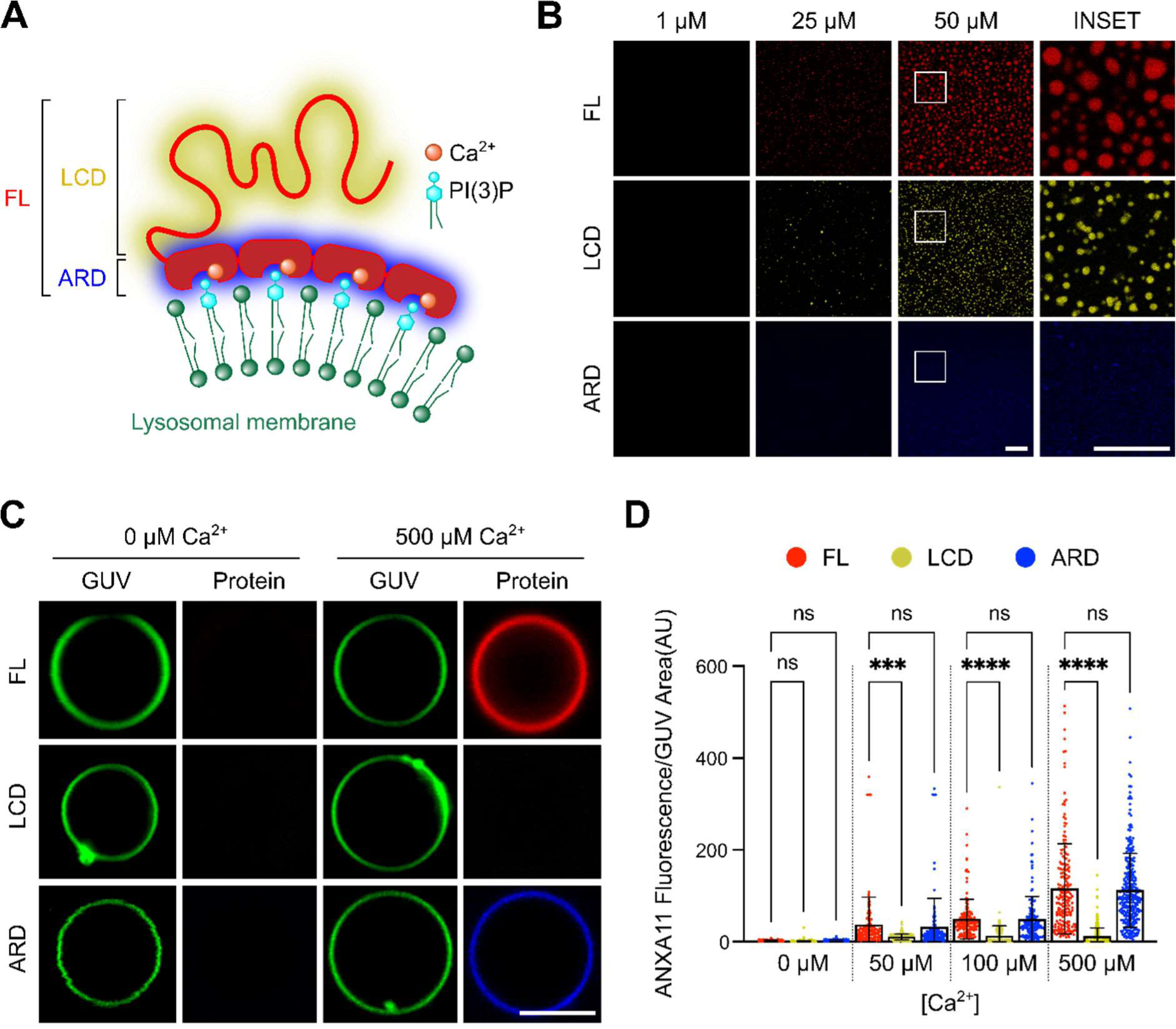
ANXA11 is a structurally bipartite protein capable of biomolecular condensation and lipid binding. **(A)** A schematic of full length (FL) ANXA11 binding to lysosomal membranes, illustrating the Ca^2+^-dependent annexin repeat domain (ARD) association with PI(3)P, and the cytosolic-facing low complexity domain (LCD). **(B)** Fluorescence micrographs of recombinant AF647-labelled FL (aa 1-502), LCD (aa 1-185) and ARD (aa 186-502) ANXA11 at varying protein concentrations. Scale bar – 5 µm. **(C)** Representative fluorescence images of ATTO488 GUVs incubated with 0.5 µM AF555-labelled ANXA11 FL, LCD and ARD in the presence or absence of 500 µM Ca^2+^. Scale bar – 5 µm. **(D)** Quantification of the fluorescence intensity of AF555-labelled FL, LCD and ARD recruited to GUVs as shown in (C) at varying Ca^2+^ concentrations. Mean ± SD. Kruskal-Wallis test with Dunn’s multiple comparison, ***p < 0.001, ****p < 0.0001, ns - not significant (p > 0.05), n=3 (110-379 GUVs).

We next investigated the roles of the LCD and ARD hemiproteins in driving ANXA11-lipid binding. As we have shown previously, ANXA11 binds to phosphatidylinositol 3-phosphate (PI(3)P)-containing GUV membranes in a Ca^2+^-dependent manner (Figure S1F-G) (Liao et al., 2019). Here we demonstrate, across a range of Ca^2+^ concentrations, that this binding is entirely dependent on the ARD. The LCD hemiprotein alone exhibits negligible recruitment to GUV membranes (Figure 1C-D).

### ANXA11-lipid binding causes a liquid-to-gel phase transition of lipid membranes

The binding of several other annexin family members has been shown to alter the organisation and phase state of lipid membranes (Lin et al., 2020; Menke et al., 2005). To interrogate this possibility with ANXA11, we first examined the mobility of fluorescently-labelled PI(3)P in our GUVs using fluorescence recovery after photobleaching (FRAP) (Figure 2A). Binding of ANXA11 FL to GUVs caused a marked reduction in PI(3)P mobility (Figure 2B). By incubating GUVs with the ARD hemiprotein, we determined that this change in lipid diffusivity is entirely attributable to ARD-lipid interactions. The LCD of ANXA11 does not bind membranes (Figure 1C-D) and so was not assessed.

**Figure 2.**
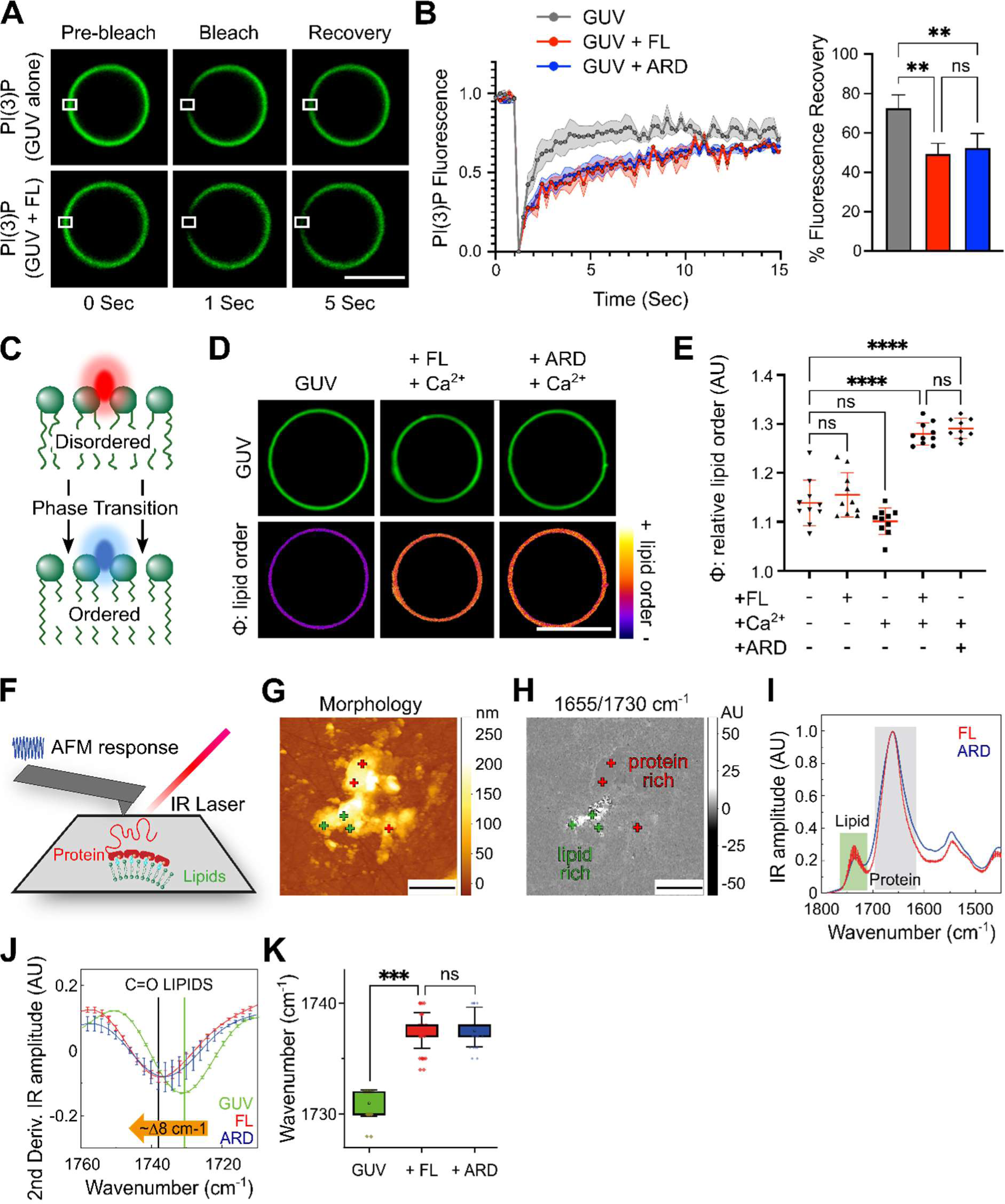
ANXA11-GUV binding causes a liquid-to-gel phase transition of lipid membranes. **(A)** Still images from a lipid FRAP series of BODIPY-PI(3)P in GUVs in the presence of 100 µM Ca^2+^ with and without 0.5 µM ANXA11 FL. Scale Bar - 5 µm. **(B)** PI(3)P fluorescence recovery rates of GUVs in the presence of 100 µM Ca^2+^ co-incubated with either 0.5 µM ANXA11 FL or 0.5 µM ARD. The % fluorescence recovery by 5 sec is plotted alongside. Mean ± SD. One-way ANOVA with Tukey’s multiple comparison, **p < 0.01, ns - not significant (p > 0.05), n=4. **(C)** A schematic illustrating the change in PK dye fluorescence emission as lipids transition from a disordered (red emission) to ordered (blue emission) state. **(D)** Fluorescence images of ATTO647 GUVs labelled with 5 µM PK dye to extract the relative lipid order (φ) of GUVs alone or with 100 µM Ca^2+^ and either 0.5 µM ANXA11 FL or 0.5 µM ARD. Scale Bar - 5 µm. **(E)** Quantification of the relative lipid order (φ) of GUVs as shown in (D), including protein alone and Ca^2+^ alone controls. Mean ± SD. One-way ANOVA with Tukey’s multiple comparison, ****p < 0.0001, ns - not significant, n=3 (30-82 GUVs). **(F)** A schematic of our AFM-IR setup which acquires nanoscale resolved chemical spectra of protein and lipid components. **(G)** An exemplary AFM-IR morphology map of GUV fragments bound to ANXA11. Green crosses indicate regions with a ‘lipid only’ signature. Red crosses indicate regions with ‘protein and lipid’ signatures. Scale bars - 2 µm. **(H)** A ratio of the infrared maps from protein:lipid (1655/1730 cm^-1^) spectroscopic signatures illustrates the heterogeneity of their spatial distribution at nanoscale. Scale bars - 2 µm. **(I)** A comparison of the average (Mean ± SD) AFM-IR spectra of GUV fragments bound to 0.5 µM ANXA11 FL and 0.5 µM ARD at 100 µM Ca^2+^. 450 spectra were collected from across 69 independent GUV fragments (12 lipid only, 31 lipid+FL, 26 lipid+ARD) across 5 experimental repeats. The SD in the ARD condition falls beneath the thickness of the line. **(J)** Second derivatives of the spectra (Mean ± SD) in the IR absorption of the C=O stretching region of lipids for GUV fragments alone or bound to either ANXA11 FL or ARD. **(K)** Quantification of the wavenumber of GUV fragments alone compared with GUV fragments bound to ANXA11 FL or ARD. The ∼1730 to ∼1738 cm^-1^ shift indicates a liquid-to-gel lipid phase transition. Mean ± SD and 25-75^th^ percentile (box). One-way ANOVA with Tukey’s multiple comparison, ***p < 0.001, ns - not significant (p > 0.05), n=5 (12-26 GUV fragments).

To determine the basis of ANXA11’s effect on lipid mobility, we utilised a solvatochromic dye (PK dye) that reports on membrane lipid phase state. The PK dye is a lipophilic push pull pyrene probe that intercalates into membranes and shifts its fluorescence emission based the local lipid order/disorder (Valanciunaite et al., 2020). The ratio between a blue/green (449-550 nm) and red (550-650 nm) emission window thus reports the relative lipid order, or phase state, of the membrane (Figure 2C). Initially, to validate the PK dye in our system, we warmed GUVs from 18 to 37 °C where we observed an expected temperature-dependent decrease in lipid order (Figure S2A) (Chen et al., 2018; Nicolini et al., 2006). We thus reasoned that we could use the PK dye to assess whether ANXA11’s effects on lipid mobility were underpinned by a phase transition in GUV membranes. Indeed, the Ca^2+^-dependent binding of either ANXA11 FL or the ARD hemiprotein to GUVs both induced a significant, and comparable, increase in lipid ordering (Figure 2D-E).

To determine the magnitude of this lipid phase transition, we employed atomic force microscopy-based infrared (IR) nanospectroscopy (AFM-IR) (Ruggeri et al., 2020, 2021). AFM-IR exploits a ∼20 nm diameter probe to simultaneously extract nanoscale morphological and chemical properties of protein and lipid biomolecular systems (Figure 2F, see STAR Methods). To perform the AFM-IR spectroscopy, we deposited GUVs coated in either ANXA11 FL or ARD hemiprotein onto hydrophobic ZnSe surfaces (Figure 2G-H, Figure S2B). From this we were able to extract distinct protein and lipid IR absorption spectra (Figure 2I, Figure S2C-F). Spectra from GUVs without bound ANXA11 possessed typical C=O peaks for phospholipids and cholesterol esters at 1730-32 cm^-1^ (green line, Figure 2J). This is consistent with the lipids existing in a liquid phase (Nicolini et al., 2006). However, binding of either ANXA11 FL or ARD caused a statistically significant 8 cm^-1^ increase in wavenumber in the position of the lipid C=O peak (Figure 2J-K). The magnitude of this shift indicates a phase transition of the lipids into a gel-like state (Mantsch and McElhaney, 1991; Tamm and Tatulian, 1997). Of note, analysis of the protein IR spectra (Figure S2G) reveals that ANXA11 does not exhibit significant secondary structural changes upon GUV binding (Figure S2H-I).

Taken together, these data indicate that ANXA11 binding to lysosomal-like GUV membranes causes a liquid-to-gel phase transition in the underlying lipids. This effect appears to be entirely mediated by the structured ARD. However, we speculated that modulation of the phase state of ANXA11, through its LCD, might influence the magnitude of this observed lipid phase transition.

### Changes in the phase state of ANXA11 mediate coupled changes in the phase state of lipid membranes

To interrogate the possibility of coupled protein-lipid phase state changes within this system, we sought to induce a phase transition in GUV-bound ANXA11 and observe how this affected the underlying lipid phase state. Typically, the phase state of protein components in *in vitro* systems can be modulated by changing the physical or chemical conditions (e.g. [salt], temperature, crowding). However, altering these parameters will also have a direct effect on the lipid phase state or integrity of GUV membranes. To avoid this issue, we exploited the unique domain architecture of ANXA11.

By adding LCD hemiprotein to FL ANXA11, we reasoned that this should promote the dispersed to condensed FL ANXA11 phase transition shown in Figure 1B. Indeed, exogenously added LCD hemiprotein co-condenses with FL protein (Figure S3A). This causes a shift in the phase boundary of FL ANXA11 in favour of the condensed state (Figure S3B), resulting in larger and more numerous condensates (Figure 3A). This is not simply a protein crowding effect, as a purified Halo protein control fails to incorporate into FL ANXA11 condensates (Figure S3A) and does not impact the phase boundary (Figure S3B) or size/number of ANXA11 condensates (Figure3A). Importantly, FRAP studies revealed that the addition of LCD hemiprotein also pushes the ANXA11 condensates into a more immobile state (Figure S3C). These changes in mobility likely represent a shift of the condensates into a gel-like phase, as they do not exhibit morphological or secondary structural changes consistent with protein aggregation (Figure S3D-E). Once again, addition of the Halo control had no effect on ANXA11 mobility (Figure S3C). Collectively, these data suggest that the addition of exogenous LCD hemiprotein to FL ANXA11 enhances condensation of the full-length protein and promotes a phase transition to a gel-like state.

**Figure 3.**
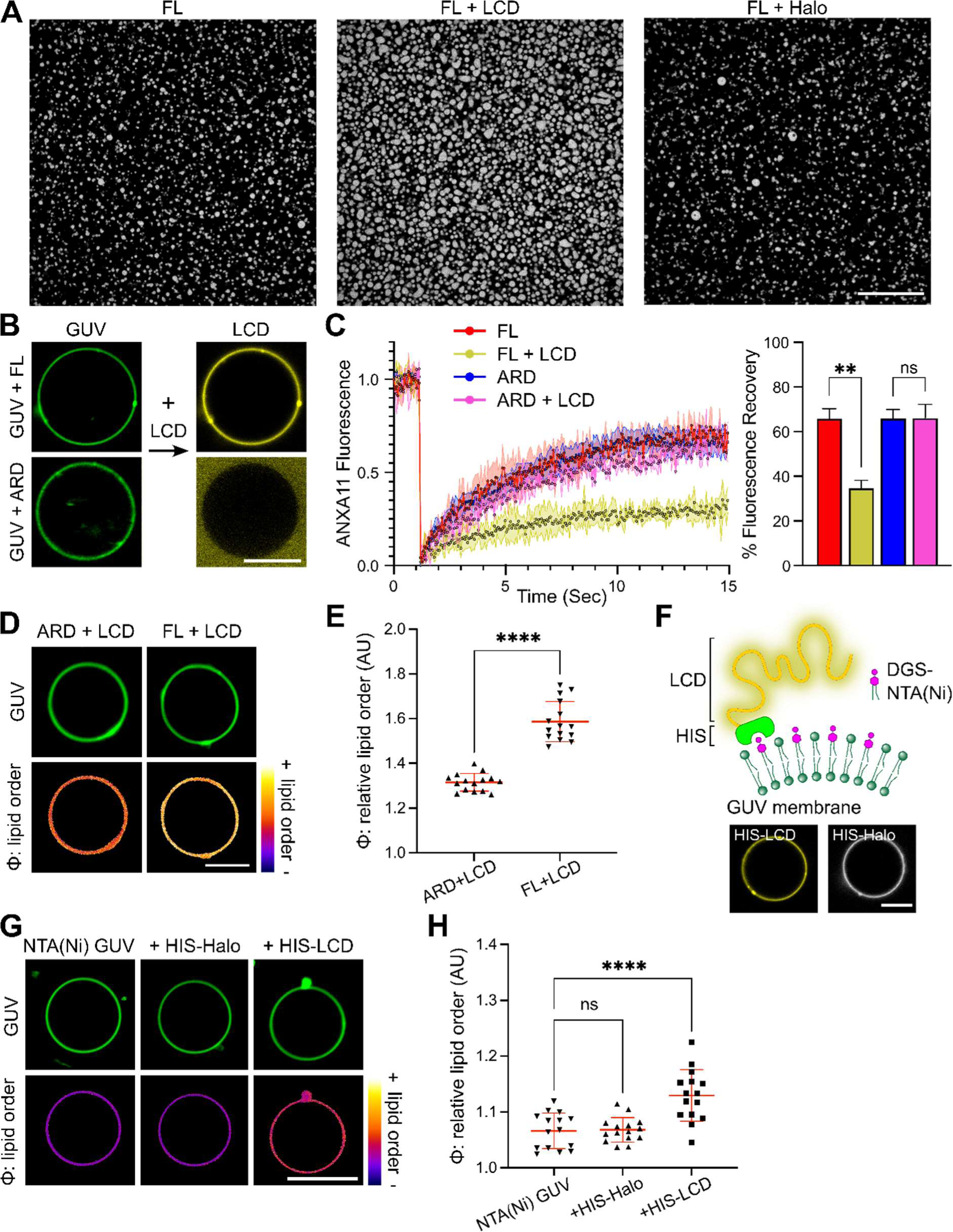
An ANXA11 phase transition induces coupled changes in the phase state of lipids in GUV membranes. **(A)** Fluorescence micrographs of 25 µM recombinant AF647-ANXA11 FL alone or in the presence of unlabelled 25 µM LCD or 25 µM Halo. Scale bar - 20 µm. **(B)** Representative images of ATTO488 GUVs coated in either 0.5 µM ANXA11 FL or 0.5 µM ARD at 100 µM Ca^2+^ (left). On the right, the same GUVs coincubated with 25 µM AF647-labelled LCD. Scale bar - 5 µm. **(C)** FRAP recovery curves of 0.5 µM AF647-labelled ANXA11 FL or ARD on the surface of GUVs at 100 µM Ca^2+^, with and without 25 µM LCD. The corresponding quantification of the % fluorescence recovery after 15 sec is plotted alongside. Mean ± SD. One-way ANOVA with Tukey’s multiple comparison, **p < 0.01, ns - not significant (p > 0.05), n=4. **(D)** Fluorescence images of ATTO647 GUVs labelled with 5 µM PK dye to extract the relative order (φ) of membrane lipids. GUVs were incubated with 100 µM Ca^2+^ and either 0.5 µM ANXA11 or ARD in the presence of 25 µM LCD. Scale bar - 5 µm. **(E)** Quantification of the relative lipid order (φ) of GUVs as shown in (D). Mean ± SD. Unpaired t-test with Welch’s correction, ****p < 0.0001, n=5 (44-126 GUVs). **(F)** A schematic illustrating HIS-LCD conjugation to DGS-NTA(Ni) lipids. Below are images showing 25 µM AF647-labelled HIS-LCD or HIS-Halo binding to NTA(Ni) GUVs. **(G)** Fluorescence images of ATTO647 GUVs labelled with 5 µM PK dye to extract the relative order (φ) of membrane lipids. NTA(Ni) GUVs were incubated with either 25 µM HIS-LCD or HIS-Halo. Scale bar - 5 µm. **(H)** Quantification of the relative lipid order (φ) of GUVs as shown in (G). Mean ± SD. One-way ANOVA with Dunnett’s multiple comparison, ****p < 0.0001, ns - not significant (p > 0.05), n=5 (83-114 GUVs).

To test whether this ANXA11 phase transition could also occur on lipid surfaces, we took GUVs coated in either ANXA11 FL or ARD hemiprotein and added exogenous fluorescently labelled LCD. The labelled LCD was recruited to ANXA11 FL-GUVs and not ARD-GUVs (Figure 3B). This recruitment is thus underpinned by co-condensation of the exogenous LCD with the LCD of the full-length ANXA11 (as seen in Figure S3A). Additionally, the exogenous LCD caused a reduction in the mobility of ANXA11 FL on GUVs (Figure 3C). This suggests that the LCD hemiprotein-induced ANXA11 phase transition observed in solution also occurs in membrane-bound ANXA11. Unsurprisingly, no change in ARD mobility occurred following the addition of exogenous LCD to ARD-GUVs (Figure 3C).

We next used our PK dye to determine whether this gel-like ANXA11 phase resulted in changes in the underlying lipid phase state (Figure 3D). We have demonstrated above (Figure 2D-E) that ANXA11 FL and the ARD hemiprotein exert comparable effects on lipid phase state. However, the induction of a gel-like phase transition in the full-length ANXA11 (through the addition of exogenous LCD) causes a significant increase in lipid order (Figure 3E). This effect is not observed on ARD-GUVs (Figure 3E).

This ANXA11-mediated effect on the lipid phase state could in principle arise by two mechanisms. Firstly, a change in the phase state of the LCD of the full length ANXA11 might indirectly impact the lipids through altering ARD-based interactions with lipid headgroups. Alternatively, a phase transition of the LCD in close proximity to lipid membranes could prove sufficient by itself to directly influence the lipid phase state. To disambiguate these two possibilities, we constructed an experimental system in which LCD hemiprotein could engage with GUV membranes in the absence of the ARD. To accomplish this, we spiked our GUVs with 10% DGS-NTA(Ni), a nickel-chelating lipid species that binds to histidine. We then purified an LCD hemiprotein containing a C-terminal 6x HIS tag. This C-terminal HIS tag permits the LCD to associate with the lipid surface in the absence of the ARD (Figure 3F). A HIS-tagged Halo protein was used as a surface binding control. PK dye readouts from the GUV membranes reveal a modest but significant increase in lipid order following binding of the HIS-LCD protein, but not the HIS-Halo control (Figure 3G-H). This indicates that phase coupling between ANXA11 and lipid membranes can occur independently from a direct contribution of the ARD.

Taken together, these findings suggest that the ARD-mediated liquid-to-gel lipid phase transition described in Figure2 can be tuned by the phase state of the ANXA11 LCD, thus providing evidence of protein-lipid phase coupling within ANXA11-GUV assemblies.

### ANXA11 interactors ALG2 and CALC modulate phase coupling within ANXA11-GUV assemblies

Prior work on other annexins has revealed that their interactions with lipid membranes can be modified by various annexin-binding proteins (Weisz and Uversky, 2020). We wondered whether similar binding partners might exist that could modulate ANXA11’s phase state and thus impact phase coupled changes in underlying lipid membranes. A bioinformatics search revealed two candidate proteins, ALG2 and CALC, which bind to small, structured motifs within the ANXA11 LCD.

Initially we co-incubated FL ANXA11 with either purified ALG2 or CALC (Figure S4A) at molar ratios matching their relative abundance in cells (1[ANXA11]:4[ALG2]:40[CALC]) (Itzhak et al., 2016). Under these conditions, neither ALG2 nor CALC form condensates by themselves (Figure S4B). However, when mixed with ANXA11, both ALG2 and CALC co-partition into FL ANXA11 condensates, unlike a Halo control protein (Figure S4C). Both modulators dramatically alter the dispersed-to-condensed phase boundary of ANXA11, however, they do so in opposite directions (Figure S4D). ALG2 significantly increases ANXA11 condensation, while CALC has the opposite effect (Figure 4A). Our FRAP studies revealed that the addition of ALG2 also pushes ANXA11 condensates into a more immobile gel-like phase (Figure S4E). Of note, this transition does not elicit changes to the secondary structure of ANXA11 that are consistent with protein aggregation (Figure S4F-G). By contrast, CALC increases the mobility of ANXA11 condensates, driving them into a more fluid phase (Figure S4E). Once again, the addition of a Halo control protein had no effect on the ANXA11 phase boundary (Figure S4D), condensate formation (Figure 4A), or ANXA11 mobility (Figure S4E). This suggests that protein crowding is unlikely to be a driving factor in the observed influence of ALG2 or CALC on ANXA11.

**Figure 4.**
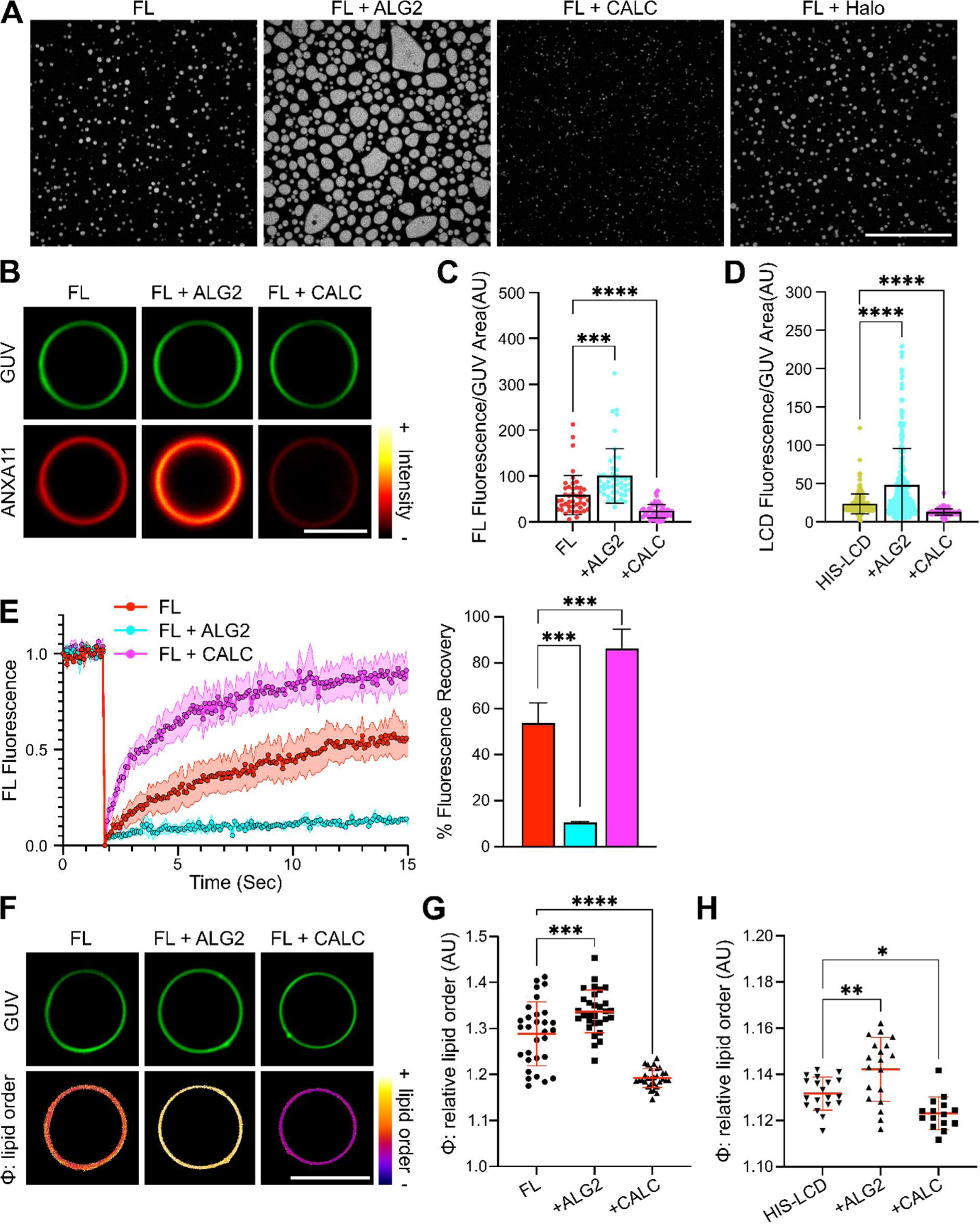
ALG2 and CALC modulate phase coupling within ANXA11-GUV assemblies. **(A)** Fluorescence micrographs of 25 µM recombinant AF647-ANXA11 FL in the presence of unlabelled 0.1 mM ALG2, 1 mM CALC or 1mM Halo. The concentrations of the modulators were determined from molar ratios matching their relative abundance in cells (1[ANXA11]:4[ALG2]:40[CALC]). Scale bar - 20 µm. **(B)** Representative fluorescence images of ATTO488 GUVs incubated with 0.5 µM AF647-ANXA11 FL at 100 µM Ca^2+^ co-incubated with either 2 µM ALG2 or 20 µM CALC. Scale bar - 5 µm. **(C)** Quantification of the fluorescence intensity of AF647-ANXA11 FL recruited to GUVs as in (B). Mean ± SD. Kruskal-Wallis test with Dunn’s multiple comparison, ***p < 0.001, ****p < 0.0001, n=3 (35-55 GUVs). **(D)** Quantification of the fluorescence intensity of 25 µM AF647-labelled HIS-LCD recruited to NTA(Ni)-GUVs co-incubated with either 0.1 mM ALG2 or 1 mM CALC. Mean ± SD. Kruskal-Wallis test with Dunn’s multiple comparison, ****p < 0.0001, n=3 (117-281 GUVs). **(E)** A FRAP recovery profile of 0.5 µM AF647-ANXA11 FL on the surface of GUVs in the presence or absence of 2 µM ALG2 or 20 µM CALC at 100 µM Ca^2+^. The corresponding quantification of the % fluorescence recovery after 15 sec is plotted alongside. Mean ± SD. One-way ANOVA with Dunnett’s multiple comparison ***p < 0.001, n=3-5. **(F)** Fluorescence images of ATTO647 GUVs labelled with 5 µM PK dye to extract the relative order (φ) of membrane lipids. GUVs were incubated with 100 µM Ca^2+^ and 0.5 µM ANXA11 FL co-incubated with either 2 µM ALG2 or 20 µM CALC. Scale bar - 5 µm. **(G)** Quantification of the relative lipid order (φ) of GUVs shown in (F). Mean ± SD. One-way ANOVA with Dunnett’s multiple comparison, ***p < 0.001, ****p < 0.0001, n=5 (58-134 GUVs). **(H)** Quantification of the relative lipid order (φ) of NTA(Ni)-GUVs labelled with 5 µM PK dye. NTA(Ni) GUVs were incubated with 25 µM HIS-LCD and with either 0.1 mM ALG2 or 1 mM CALC. Mean ± SD. One-way ANOVA with Dunnett’s multiple comparison, *p < 0.05, **p < 0.01, n=4 (28-94 GUVs).

Having established ALG2 and CALC as robust antipodal regulators of ANXA11 phase state, we next sought to test their influence on ANXA11-coated lipid membranes. By themselves, neither ALG2 nor CALC associate with GUV membranes (Figure S5A). However, both modulators are recruited to the surface of GUVs when FL ANXA11 is present (Figure S5A). Here, ALG2 acts to increase FL ANXA11 recruitment to GUVs, while CALC decreases it (Figure 4B-C). These effects are likely mediated through the aforementioned influence of ALG2 and CALC on LCD-based ANXA11 condensation. In support of this, neither ALG2 nor CALC altered recruitment of ARD hemiprotein to GUVs (Figure S5B-C). By contrast, the recruitment of HIS-LCD hemiprotein to DGS-NTA(Ni)-containing GUVs (as described in Figure 3F) was increased by ALG2 and impaired by CALC (Figure 4D). These differential effects of ALG2 and CALC on FL ANXA11 recruitment are accompanied by parallel changes in ANXA11 mobility on GUVs as measured by FRAP. Thus, ALG2 reduces and CALC increases FL ANXA11 mobility on GUV surfaces (Figure 4E). Neither modulator protein influences the mobility of ARD hemiprotein (Figure S5D). These data suggest that ALG2 promotes ANXA11 condensation on the surface of GUVs, driving a phase transition of ANXA11 into a more gel-like state. CALC has the opposite effects of minimising ANXA11 condensation and driving the protein into a more fluid state.

Using PK dye labelled GUVs, we next explored whether ALG2- or CALC-based phase transitions in ANXA11 resulted in coupled changes in the underlying lipid phase state. The addition of ALG2 to FL ANXA11-coated GUVs increased lipid order, while CALC decreased it (Figure 4F-G). Crucially, ALG2 and CALC alone do not impact the lipid phase state, and their effects require the ANXA11 LCD (Figure S5E). Indeed, ALG2 and CALC can modulate the phase state of lipids through HIS-LCD hemiprotein directly conjugated to DGS-NTA(Ni)-containing GUVs (Figure 4H). This demonstrates that LCD-based phase coupling between ANXA11 and lipids can occur in the complete absence of the ARD. Control experiments using HIS-Halo protein in place of HIS-LCD reveal no effect on lipid ordering following HIS-Halo membrane association, and no modulation by ALG2 or CALC (Figure S5F). Collectively, this set of experiments identify ALG2 and CALC as putative physiological regulators of ANXA11-lipid phase coupling.

### Effects of ALG2 and CALC on phase coupling in ANXA11-lysosome complexes in living cells

We next looked to determine whether phase coupling between ANXA11 and lipid membranes occurs in living cells. To do this, we investigated whether expression of ANXA11 with either ALG2 or CALC altered the lipid phase state of lysosomal membranes in U20S cells.

Lysosomes are highly dynamic organelles that are typically small and often close to or beneath the resolution limit of standard fluorescence microscopes. Resolving lysosomal membranes and their lipid properties using optical based approaches is therefore challenging. To circumvent these issues, we briefly exposed U2OS cells to a hypotonic buffer to both arrest lysosomal motion and induce lysosomal swelling. Previous work has demonstrated this approach as a powerful method for studying membrane organisation and lipid phase behaviour inside living cells (Guillén-Samander et al., 2021; King et al., 2020; Speckner et al., 2021). It is important to note that exposure to hypotonic conditions might change the morphology of ANXA11 condensates and likely alters the precise lipid ordering of lysosomal membranes. As such, these results should be interpreted comparatively across conditions, and not as absolute measurements. We first validated this approach in U2OS cells by showing that lysosomes retain their morphology and lumenal content upon swelling (Figure S6A).

Under these conditions, fluorescently labelled ANXA11 also retains its previously reported (Liao et al., 2019) localisation to lysosomal membranes (Figure 5A). We thus applied the same PK dye method as above to assess lipid ordering in lysosomal membranes of cells transfected with either ANXA11 alone, or ANXA11 + ALG2, or ANXA11 + CALC. Expression levels of the relative proteins were elevated as expected following transfection (Figure S6B). Because the PK dye non-specifically labels all membranes, these experiments were conducted in the presence of a lysosomal membrane marker to permit lysosome-specific PK dye analysis (see STAR Methods, Figure 5B). The expression of ALG2 or CALC alone, without ANXA11, exert no detectable effect on lipid order in lysosomal membranes (Figure S6C). However, co-expression of ALG2 together with ANXA11 significantly increases the order of lysosomal membranes (Figure 5C), thereby confirming that the ANXA11-mediated phase coupling effects observed in biochemical reconstitution systems can also occur in cells. Co-expression of CALC with ANXA11 also does not significantly alter lysosomal lipid ordering. This is likely because under basal conditions lysosomal membranes exist in a relatively disordered state (Yang et al., 2000). As such, the influence of CALC is not detectable under these experimental conditions (Figure 5C).

**Figure 5.**
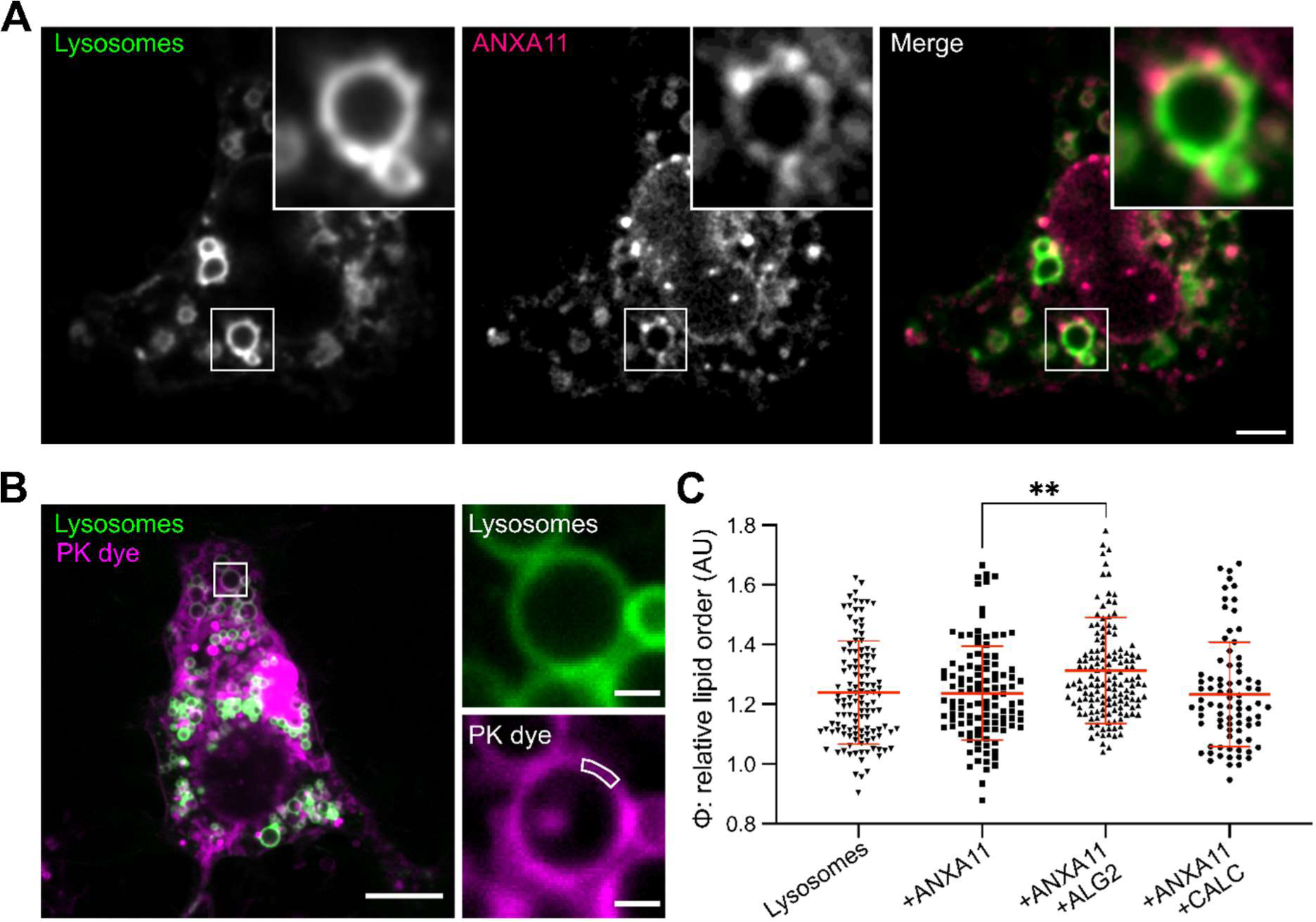
ALG2 acts via ANXA11 to increase the lipid order of lysosomes in cells. **(A)** U2OS cells under hypotonic conditions (95% ddH_2_O, 5% DMEM, pH 7.0) expressing a fluorescently labelled lysosome marker (TMEM192-Halo-JF647) and ANXA11 (ANXA11-mEm). The cytosolic pool of ANXA11 was photobleached to reveal ANXA11-membrane associations. The white ROI indicates the displayed zoomed regions. Scale bar - 5 µm. **(B)** A representative image of a hypotonic U2OS cell with fluorescently labelled lysosomes (TMEM192-Halo-JF647) incubated in 100 nM PK dye to extract the relative order of lysosomal membrane lipids. The white ROI within the zoomed panels indicates an example analysis segment. Scale bars - 10 µm (left panel) and 1 µm(right panel). **(C)** Quantitation of the relative order (φ) of lysosomal membranes as displayed in (B) from U2OS cells alone, or expressing either ANXA11, ANXA11+ALG2, or ANXA11+CALC. Mean ± SD. One-way ANOVA with Tukey’s multiple comparison, **p < 0.01, n=3 (82-146 lysosomes).

### ALG2 and CALC modulate the nanomechanical properties of ANXA11-lipid assemblies and their ability to engage RNP granules

As discussed, ANXA11 plays an important role in long-distance transport of RNP granules in neurons by tethering them to lysosomes. We wondered whether the effects of ALG2 and CALC on ANXA11-lipid phase coupling might: (i) alter the nanomechanical properties of the ANXA11-lysosome ensemble, perhaps to resist disruption by shear stress during rapid axon transport (Shen et al., 2020), or (ii) impact the RNP granule-tethering capacity of the ensemble.

To assess the resistance of ANXA11-GUV assemblies to shear stress we designed a microfluidic-based mechanical stiffness assay (Figure 6A, see STAR methods). The deformation of GUVs (strain) was recorded under different flow rates (stress) as they were pushed back and forth into V-shaped channels which tapered in diameter (Figure 6B). From the relationship between the applied pressure and the elongation of the GUV, its relative elastic modulus could be calculated (Xu et al., 2022). Binding of ANXA11 to GUVs resulted in a substantial increase in the relative elastic modulus (Figure 6C), reflecting the ARD-based liquid-to-gel lipid phase transition described in Figure2. Of particular interest, the addition of ALG2 alongside ANXA11 caused a further increase in relative elastic modulus. Adding CALC had the opposite effect, reducing the relative elastic modulus of the GUVs. ALG2 and CALC on their own can not impact the elastic modulus as they are unable to bind GUVs (Figure S5A). These findings suggest that the ALG2- and CALC-based effects on phase coupling described in Figure 4 profoundly impact the nanomechanical properties of ANXA11-lipid assemblies.

**Figure 6.**
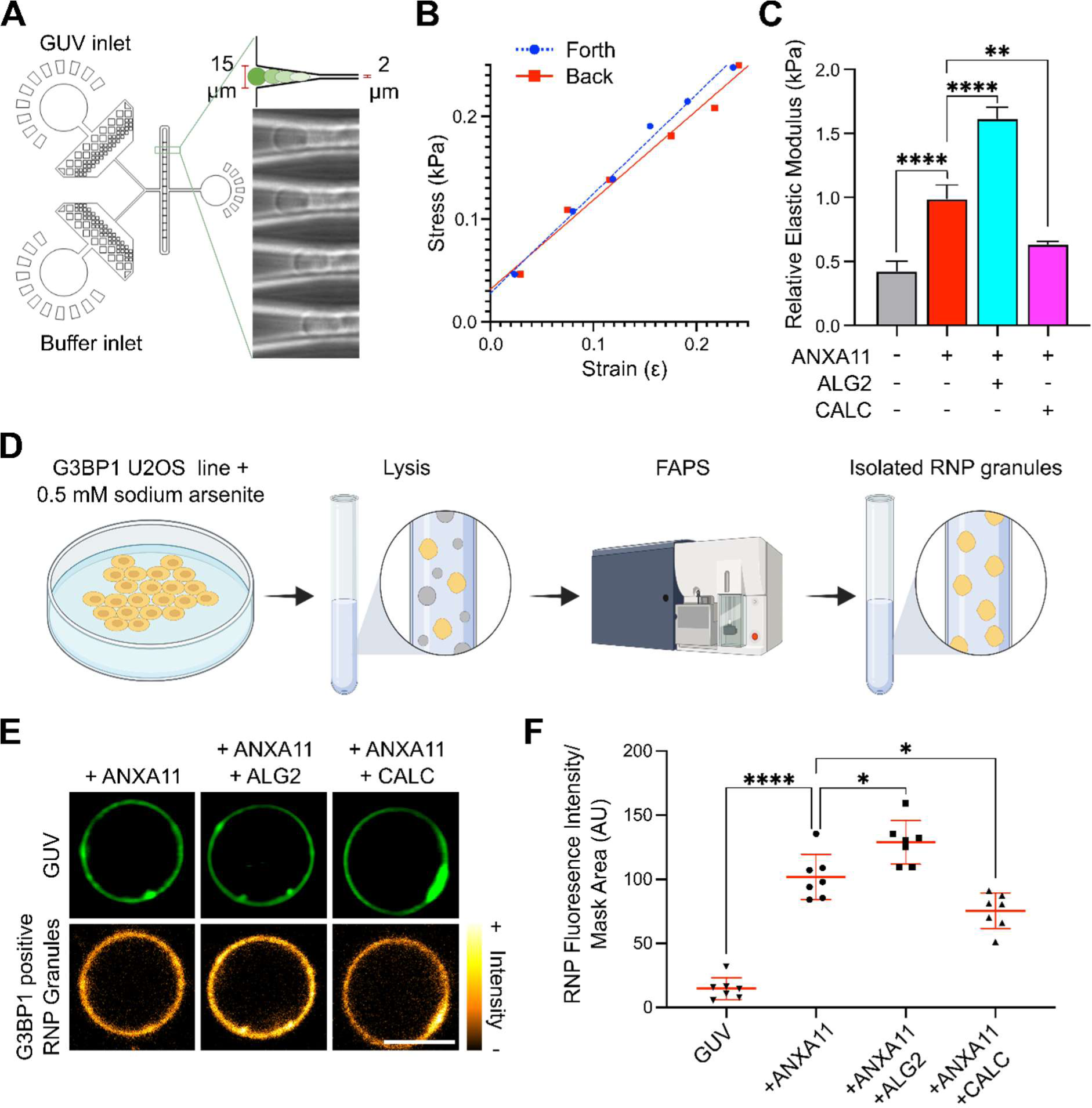
ALG2 and CALC alter the nanomechanical properties of ANXA11-GUV assemblies and their ability to tether RNP granules. **(A)** A Schematic of our microfluidic device used to extract the relative elastic modulus of GUVs. The bright-field images on the right illustrate GUV deformation within the device with a channel size: opening = 15 µm, tapered end = 2 µm. **(B)** A plot of the GUV deformation (strain) under variable pressure applied across the V-shaped channel (stress). Simple linear regression, R^2^ (Forth/Back) – 0.991/0.980. ANCOVA-slopes (p=0.26) and intercepts (p=0.13) are not significantly different. **(C)** Quantification of the relative elastic modulus of GUVs at 100 µM Ca^2+^ with 0.5 µM ANXA11 FL, coincubated with either 2 µM ALG2 or 20 µM CALC. Mean ± SD. One-way ANOVA with Tukey’s multiple comparison, **p < 0.01, ****p < 0.0001, n=3. **(D)** The experimental pipeline for FAPS-based RNP granule isolation from a stable G3BP1-mEmerald U2OS line. **(E)** Representative fluorescence images of ATTO594 GUVs incubated with 100µM Ca^2+^ and 0.5µM ANXA11 co-incubated with either 2µM ALG2 or 20µM CALC. To each condition, purified RNP granules (labelled with mEmerald-G3BP1) were added to a final concentration of 0.2 mg/ml. Scale bar - 5 µm. **(F)** Quantification of the fluorescence intensity of RNP granules (mEm-G3BP1) recruited to ANXA11-GUV assemblies as displayed in (E). Mean ± SD. One-way ANOVA with Tukey’s multiple comparison, *p < 0.05, ****p < 0.0001, n=7 (21-78 GUV).

To explore the impact of ALG2 and CALC on the capacity of ANXA11-GUV assemblies to engage RNP granules, we next established an *in vitro* RNP granule binding assay (see STAR methods). First, we isolated RNP granules from live U2OS cells using fluorescence activated particle sorting (FAPS) (Hubstenberger et al., 2017). This process involved: (i) culturing of a stable U2OS line expressing a fluorescent RNP granule marker (G3BP1-mEmerald, Figure S7A); (ii) inducing RNP granule formation prior to cell lysis; and (iii) FAPS-isolating purified RNP granules (Figure 6D). Mass spectrometry of the FAPS-isolated material confirmed the presence of numerous proteins previously identified as core components in RNP granules within living cells (Table S1).

Co-incubation of the FAPS-purified RNP granules with ANXA11-coated GUVs results in accumulation of the RNP granules onto the GUV surface (Figure 6E). In further support of the fidelity of FAPS-isolated RNP granules, recombinantly purified G3BP1 alone is not recruited to ANXA11-GUVs (Figure S7B). Also, RNP granules do not bind to GUVs alone, or ARD-coated GUVs (Figure S7C-D). Thus, RNP granule recruitment to GUVs requires FL ANXA11 holoprotein. Crucially, the addition of ALG2 increases RNP granule binding to ANXA11-GUVs, while CALC decreases it (Figure 6F). This demonstrates that ALG2 and CALC modify the ability of ANXA11 to tether RNP granules to lysosomal phospholipids.

These data suggest that ALG2 and CALC-mediated effects on phase coupling impact the nanomechanical properties of the ANXA11-lysosome ensemble, which correlates with the capacity of the ensemble to engage RNP granules. Together, these functional consequences have important implications for how RNP granule trafficking might be regulated in neurons (discussed below).

## DISCUSSION

The experiments that we describe here uncover several important observations. Principal amongst them is the finding that in juxta-membrane biomolecular condensates there is rich crosstalk between the biophysical states of protein and lipid components.

Specifically, we show that ANXA11 undergoes a dispersed to condensed phase transition to form biomolecular condensates, and that this effect requires the N-terminal low complexity domain (Liao et al., 2019). In parallel, we show that the binding of the C-terminal annexin repeat domain to lysosomal-like membranes causes a liquid-to-gel phase transition in the underlying lipids. This observation is in agreement with prior work in artificial membranes using ANX5 and ANXA2 (Gokhale et al., 2005; Lin et al., 2020; Menke et al., 2005). Both ANX5 and ANAXA2 contain four annexin repeat motifs that are homologous to the ANXA11 ARD, and both elicit lipid phase transitions or clustering of specific lipid species in synthetic membranes (Gokhale et al., 2005; Lin et al., 2020; Menke et al., 2005). However, ANXA5 and ANXA2 lack intrinsically disordered low complexity domains at their N-termini. As a result, phase coupling between protein and lipid components could not be explored for either ANX5 or ANXA2.

We have taken advantage of the presence of the ANXA11 LCD to explore the possibility of phase coupling. We have shown that the addition of purified LCD hemiprotein to full length ANXA11 promotes protein condensation, ultimately driving ANXA11 condensates into a less mobile, more gel-like state. On membrane surfaces, this phase transition in ANXA11 acts to tune the magnitude of the ARD-based liquid-to-gel phase transition in the subjacent lipids. Additionally, the role of the ANXA11 ARD can be replaced by direct chemical conjugation of the ANXA11 LCD to membrane lipids. In this scenario an increase in lipid membrane order is still observed (albeit to a lesser extent than with the ARD present). This indicates that the phase state of the ANXA11 LCD can be directly coupled with the phase state/ordering of underlying membrane lipids. We use the term phase coupling to denote this phenomenon. Recent work suggests that similar protein-lipid phase coupling occurs at the plasma membrane and plays an important role in T-cell activation, suggesting a broader role such mechanisms across cellular systems (Chung et al., 2021; Wang et al., 2022).

In this study we also identify two known ANXA11 interacting proteins, ALG2 and CALC, as potent regulators of phase coupling in ANXA11-lipid assemblies. Specifically, ALG2 promotes condensation and a gel-like phase transition of ANXA11, which in turn results in a coupled increase in lipid order. By contrast, CALC impairs ANXA11 condensation, fluidising the protein, which causes a coupled decrease in lipid order. Elucidating the structural mechanisms by which ALG2 and CALC have such differing effects requires further structural studies. However, we speculate that the different locations of their binding motifs within the ANXA11 LCD (Satoh et al., 2002; Watanabe et al., 1993) could promote/disrupt the necessary inter- and intra-molecular interactions required for ANXA11 condensation.

Additionally, we have shown that a functional consequence of ALG2/CALC-based alterations in phase coupling is the modulation of the nanomechanical properties of the ANXA11-lysosome ensemble, which correlates with its capacity to engage RNP granules. These observations have several interesting biological implications. We speculate that ALG2-based stiffening of the ensemble might enable the complex to withstand shear stresses during cotransport through the crowded axonal cytoplasm. Indeed, we have recently shown that shear stress itself can increase stiffness of condensates comprised of FUS and RNA (components of RNP granules) (Shen et al., 2020). Conversely, CALC, by softening the ensemble, could facilitate on/off-loading of the RNP granule from the ensemble at specific ‘pick-up’ or ‘drop off’ locations. Future studies in appropriate cell types will be needed to examine these hypotheses. Changes in the mechanical towing properties of the ensemble are not the only biologically significant effect that phase state changes to ANXA11-lysosome assemblies might impart. For example, membranes of the vacuole, the yeast lysosomal counterpart, have been shown to undergo lipid phase transitions to form lipid ordered (Lo) and lipid disordered (Ld) domains which have been shown to regulate docking to other organelles (Seo et al., 2021). Additionally, phase transitions of lysosomal membranes have been implicated across various stages of lipid biogenesis, transport and metabolism (Díaz et al., 2021; Thelen and Zoncu, 2017). A role for ANXA11 in these processes would offer an interesting topic for future exploration and ultimately might offer important insights into the mechanisms of ALS-associated ANXA11 mutations (Topp et al., 2017).

In summary, our study identifies protein-lipid phase coupling as an important phenomenon in governing the mechanical properties and tethering capacity of the RNP granule-ANXA11-lysosome ensemble. These findings offer a previously unrecognised mechanism for determining the biophysical state and organisation of biomolecular condensates and their associated membranous partners. However, several exciting and important questions remain for future work. Principal amongst these is how sequence differences across low complexity domains and how the biophysical properties of different biomolecular condensates contribute both to the magnitude of phase coupling as well as to the specificity and regulation of phase coupling mechanisms across cellular systems.

## ACKNOWLEDGMENTS

We thank the Klymchenko lab for the kind gift of the push pull pyrene solvatochromic PK dye. We also thank Barbara Diaz-Roher and Volker Gerke for their critical feedback during preparation of the manuscript. In addition, we thank Matthew J Gratian and Mark Bowen of the CIMR microscopy core facility and Robin Antrobus of the CIMR proteomics facility for important technical assistance.

## AUTHOR CONTRIBUTIONS

JNA, FSR, GW, TWH, MAC, YS, TPJK, MV and PStGH designed experiments. Molecular biology and protein purification was performed by JNA, SQ, WM, JH and SHW in the PStGH lab. Microscopy based experiments were conducted by JNA, GW, JEC and MSF. Temperature cycling experiments were performed by JLWatson in the lab of ED. NMR was carried out by JLWagstaff at the LMB core facility run by SMVF. Diffusional sizing, mechanical stiffness assays and the generation of phase diagrams were performed by TWH, TS, MAC, YS, YL and YS in the TPJK lab. Spectroscopy and nanoscale experiments were undertaken in the FSR, TPJK and MV labs. CK and JNA performed experiments in cells in the MEW and PStGH labs. The manuscript was written by JNA and PStGH and edited by all other authors. Funding was provided by JNA, FSR, SJM, MEW, ED, MV, TPJK and PStGH.

## DECLARATION OF INTERESTS

Tuomas Knowles and Peter St George-Hyslop are co-founders of TransitionBio. Jonathon Nixon-Abell and Seema Qamar are consultants in TransitionBio. TransitionBio has no involvement in the work described in this paper, but has an interest in biomolecular condensates in cancer and infectious disease.

**Figure S1.**
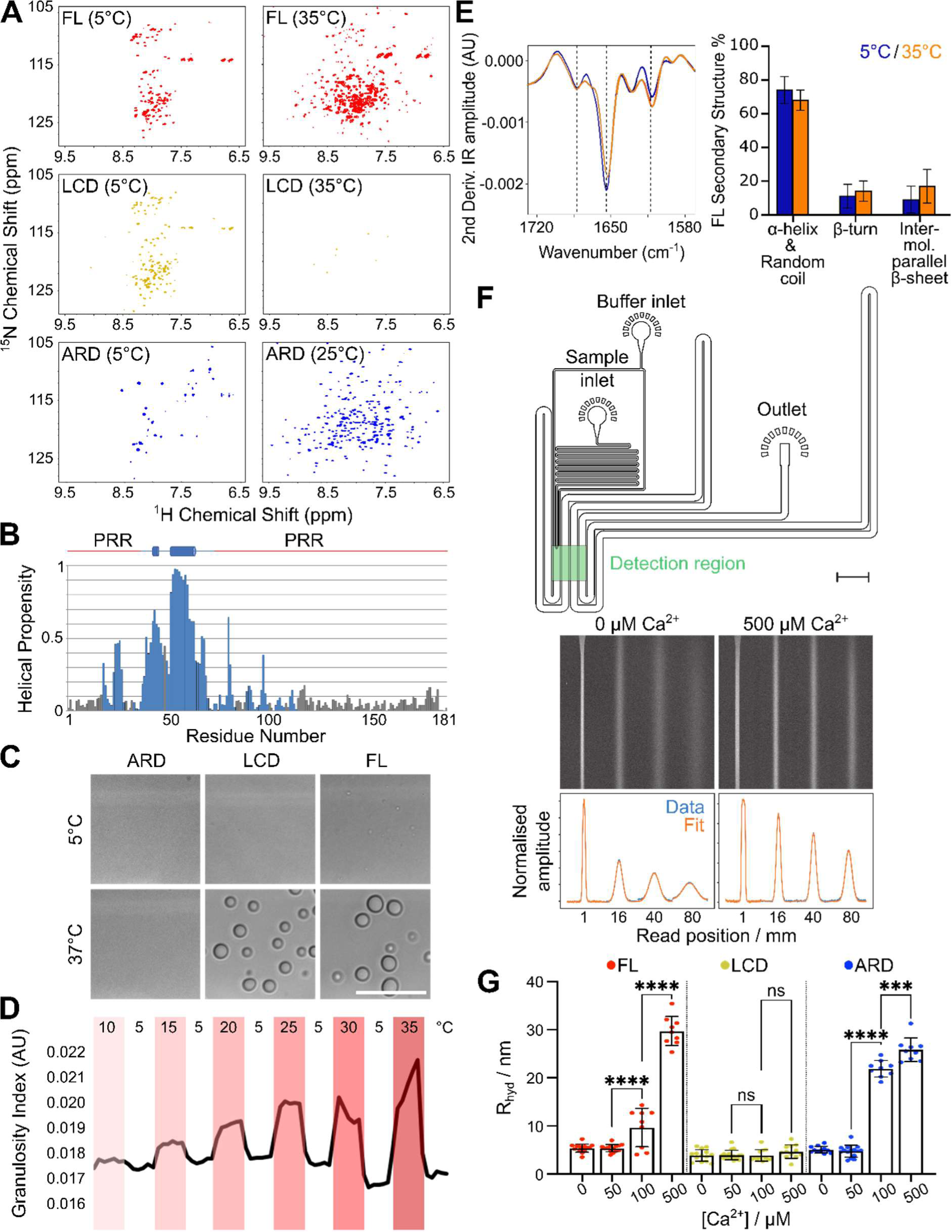
ANXA11 condensation and lipid membrane binding. **(A)** ^1^H-^15^N BEST-TROSY spectra of ANXA11 FL and LCD collected at 5°C and 35°C, and ARD collected at 5°C and 25°C. Each non-proline residue is represented by a peak in the spectrum, its location is indicative of the protein fold. A narrow ^1^H distribution of peaks characterizes disordered residues as observed for the LCD protein at 5°C, a wide ^1^H distribution is seen for folded, globular proteins such as for the ARD at 25°C. These spectra illustrate the individual temperature dependency of the two domains. **(B)** Talos-N analysis based on the partial backbone assignment (blue residues) of LCD reveals two helical regions. Assignment of the two flanking, proline-rich regions (PRR) remains incomplete, however secondary structure predictions by Talos-N based on primary sequence are also reported (grey residues). **(C)** Bright-field images of 50 µM ANXA11 FL, LCD, and ARD imaged at 5 °C and 37 °C. Scale bar – 5 µm. **(D)** Quantification of 50 µM AF647 ANXA11 FL condensate formation during repeated cycles of temperature transitions. The granulosity index assesses the level of AXA11 condensation at any given temperature (see STAR methods). **(E)** Second derivative FTIR spectra of the Amide I protein IR absorption region. Spectra were collected from 80 µM ANXA11 FL at 5 °C (dispersed state) and 35 °C (condensed state). The three dashed lines mark β-turn (left), α-helix and random coil (middle), and intermolecular parallel β-sheet (right) peak positions. On the right, a quantification of the % composition of ANXA11 secondary structure (Mean ± SD), demonstrating negligible changes between the dispersed and condensed state. Three spectra (256 co-averages) were collected across 5 experimental repeats. **(F)** Design of the microfluidic device for the diffusional sizing assay. Representative fluorescence micrographs show AF647-labelled ANXA11 FL at 0 and 500 µM Ca^2+^ incubated with SUVs as they pass through the microfluidic device. These data were analysed to obtain the diffusional profiles and fitted with numerical model simulations to obtain the hydrodynamic radius (see STAR methods). Scale bar – 1 mm. **(G)** Quantification of the hydrodynamic radius as established in (F) for SUVs incubated with 0.5 µM ANXA11 FL, LCD, or ARD at varying Ca^2+^ concentrations. Mean ± SD. One-way ANOVA with Tukey’s multiple comparison, ***p < 0.001, ****p < 0.0001, ns - not significant (p > 0.05), n=9 (1.2-2.3×10^8^ SUVs).

**Figure S2.**
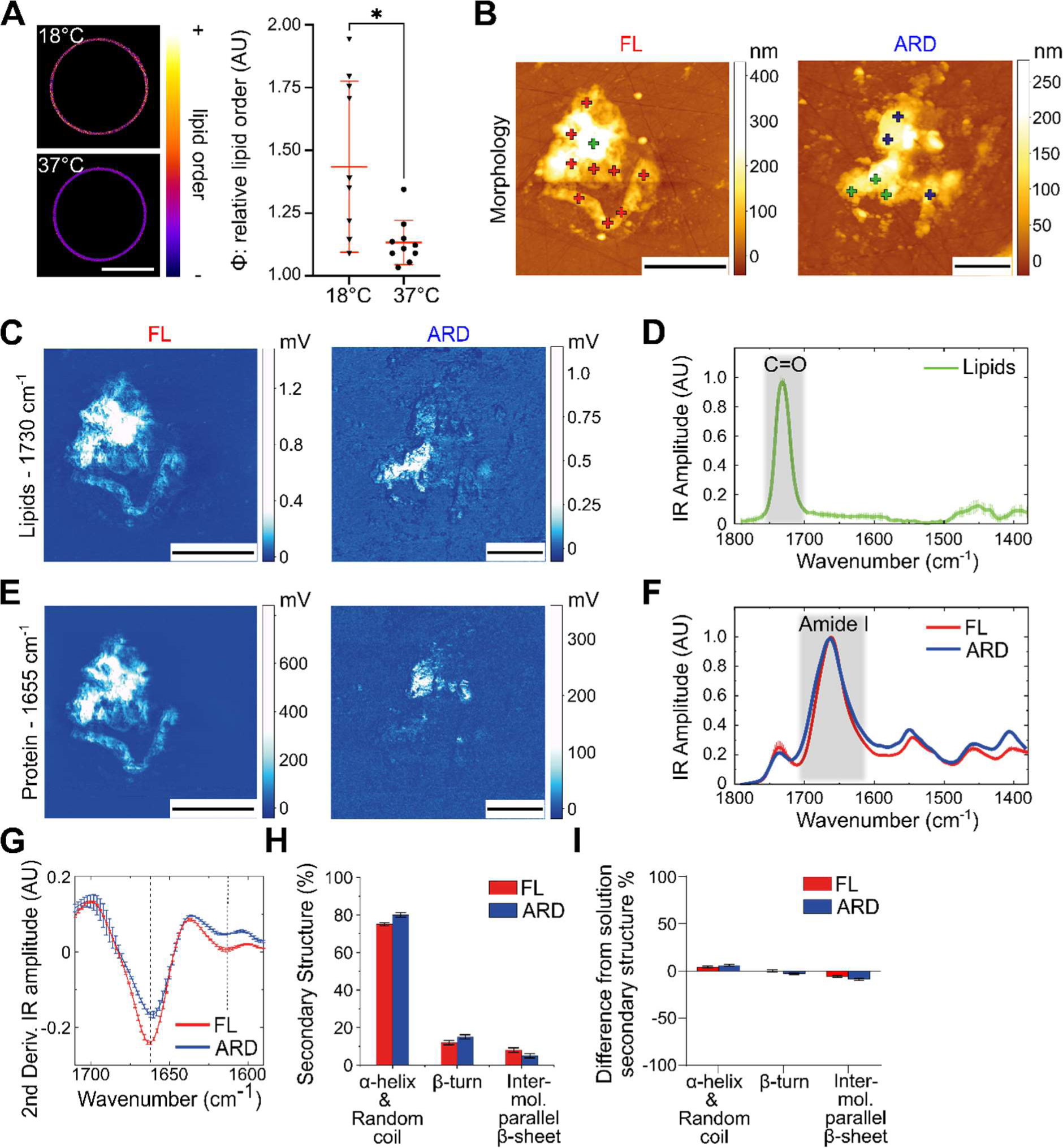
Chemical changes in protein and lipid components within ANXA11-GUV assemblies. **(A)** Fluorescence images of GUVs incubated with 5 µM PK dye to extract the relative order (φ) of membrane lipids at 18 °C and 37 °C. Corresponding quantification is plotted alongside. Mean ± SD. Unpaired t-test with Welch’s correction, *p < 0.05, n=3 (10 GUVs). **(B)** AFM-IR morphology maps of GUV fragments bound to 0.5 µM ANXA11 FL and 0.5 µM ARD at 100 µM Ca^2+^. Scale bars - 5 µm (FL), 2 µm (ARD). N.B. the ARD panel shown here is used as the example in Figure 2G. **(C)** Nanoscale-resolved IR absorption maps collected at 1730 cm-1 (C=O stretching lipid region) of the ANXA11 FL and ARD GUVs described in (B). Scale bars - 5 µm (FL), 2 µm (ARD). **(D)** IR spectra (Mean ± SD, n=12) collected from ‘lipid only’ regions (e.g. green crosses in (B)) show a maximum peak of absorption at 1730 cm^-1^ (C=O, lipids). **(E)** IR absorption maps collected at 1655 cm^-1^ (Amide I protein region) of the ANXA11 FL and ARD GUVs described in (B). Scale bars - 5 µm (FL), 2 µm (ARD). **(F)** IR spectra (Mean ± SD) collected from GUV fragments bound to ANXA11 FL (e.g. red crosses in (B), n=31) or ARD (e.g. blue crosses in (B), n=26) show a maximum peak of absorption at 1655 cm^-1^ (Amide I, protein). The SD in the ARD condition falls beneath the width of the line. **(G)** Second derivative IR spectra of the Amide I protein IR absorption region displayed in (F). The two dashed lines mark α-helix and random coil (left), and intermolecular parallel β-sheet (right) peak positions. **(H)** The secondary structural composition (Mean ± SD) derived from (G) of ANXA11-FL and ARD when bound to GUVs. **(I)** Difference-spectroscopy quantification of the deviation of the secondary structure (Mean ± SD) of ANXA11 FL and ARD bound to GUVs compared to the native secondary structure of the protein in solution (Figure S1E). AFM-IR data reported in this figure was collected from across 69 independent GUV fragments (12 lipid only, 31 lipid+FL, 26 lipid+ARD) across 5 experimental repeats.

**Figure S3.**
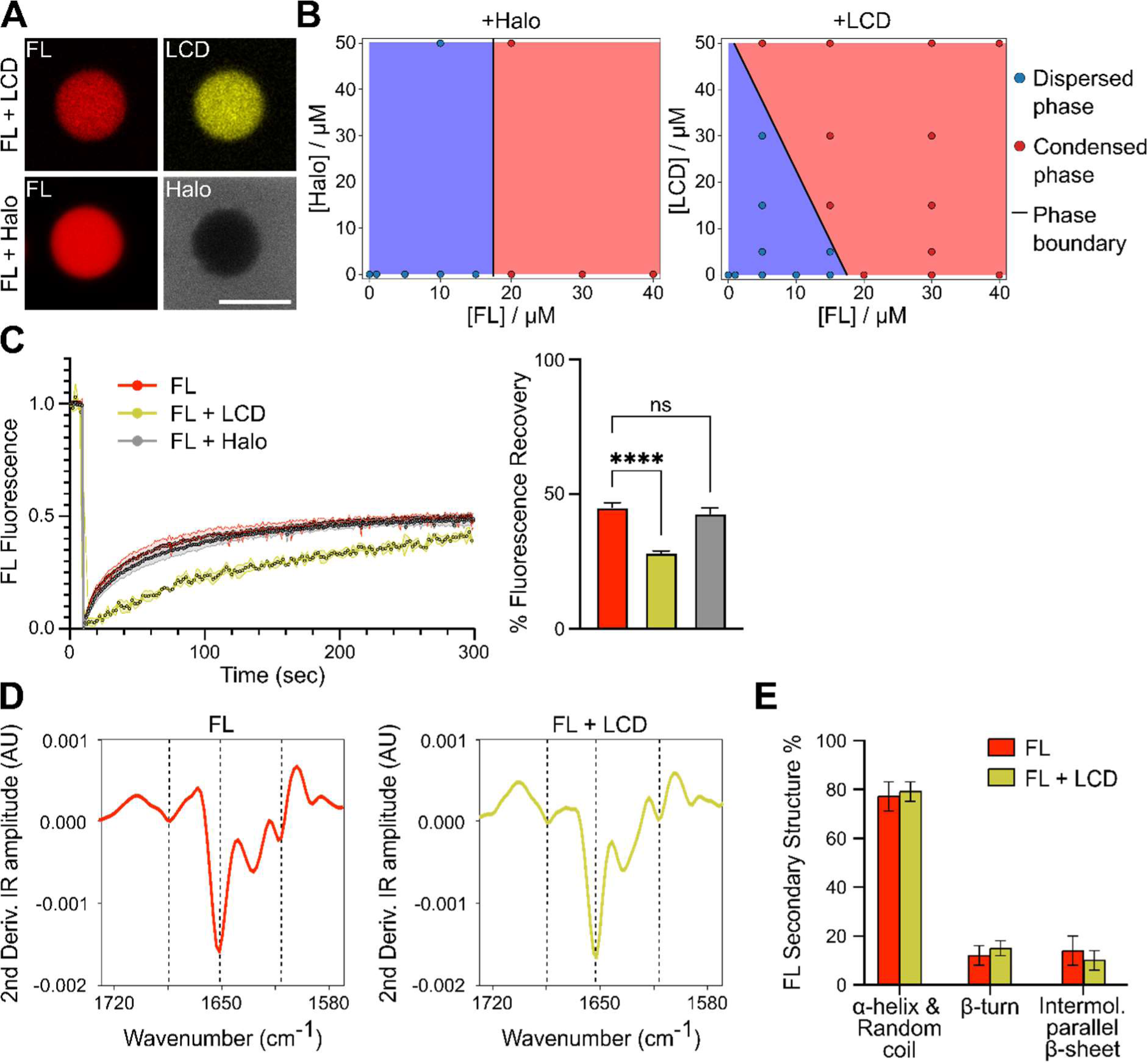
The impact of exogenous LCD-hemiprotein on ANXA11 phase state. **(A)** Fluorescence micrograph of individual AF647-ANXA11 FL condensates co-incubated with either AF488-LCD or JF549-Halo. Scale bar - 2.5 µm. **(B)** Phase diagrams plotting [ANXA11 FL] against either [Halo] or [LCD], with the phase boundary between the dispersed and condensed state marked with a black line. Each dot represents an imaging datapoint in which ANXA11 condensates were present (condensed phase), or not (dispersed phase) (see STAR methods). **(C)** FRAP recovery curves of 25 µM recombinant AF647-ANXA11 FL alone or in the presence of unlabelled 25 µM LCD or 25 µM Halo. The corresponding quantification of the % fluorescence recovery at 150 sec is plotted alongside. Mean ± SD. One-way ANOVA with Dunnett’s multiple comparison, ****p < 0.0001, ns - not significant (p > 0.05), n=4. N.B. The ANXA11 FL dataset displayed here is the same as in Figure S4E as experiments were performed concurrently **(D)** Second derivative FTIR spectra of the Amide I protein IR absorption region. Spectra collected are from 150 µM ANXA11 FL alone (left) or coincubated with 150 µM LCD (right). The three dashed lines mark β-turn (left), α-helix and random coil (middle), and intermolecular parallel β-sheet (right) peak positions. **(E)** Quantification of the % composition of ANXA11 secondary structure (Mean ± SD). Note the low % of intermolecular parallel beta sheet, even after addition of LCD. 19 spectra were collected across 5 experimental repeats. N.B. The FL dataset here is the same as in Figure 4F as the experiments were conducted concurrently.

**Figure S4.**
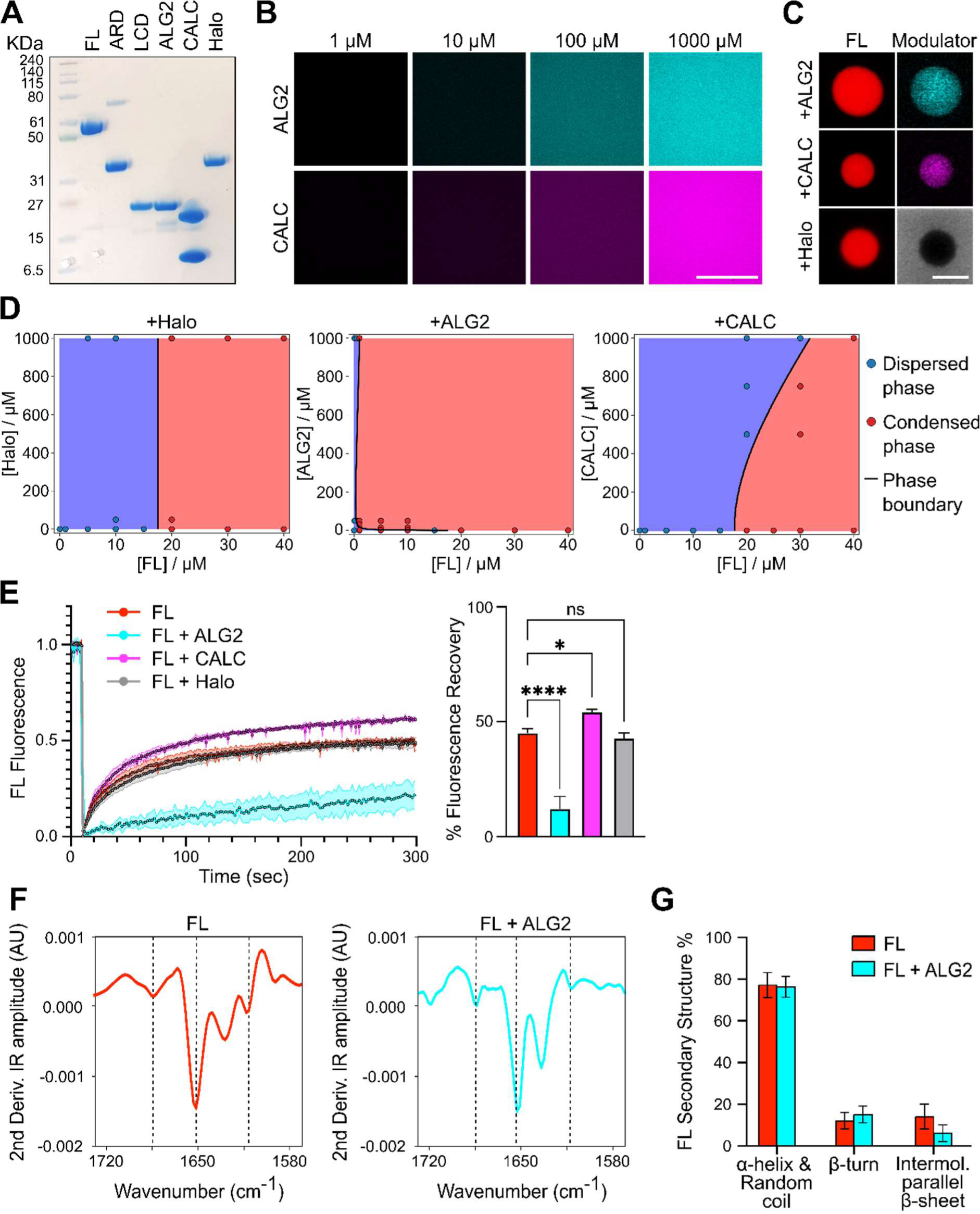
The impact of ALG2 and CALC on ANXA11 phase state. **(A)** Coomassie stained SDS-PAGE gel demonstrating the purity of recombinant proteins used throughout this study. **(B)** Fluorescence images of purified AF488-ALG2 and AF488-CALC at various protein concentrations. Note there are no visible condensates formed **(C)** Fluorescence micrograph of individual AF647-ANXA11 FL condensates co-incubated with either AF488-labelled ALG2, CALC, or Halo. Scale bar - 2.5 µm. **(D)** Phase diagrams plotting [ANXA11 FL] against either [ALG2] or [CALC], with the phase boundary between the dispersed and condensed state marked with a black line. Each dot represents an imaging datapoint in which ANXA11 condensates were present (condensed phase), or not (dispersed phase) (see STAR methods). **(E)** FRAP recovery curves of 25 µM recombinant AF647-ANXA11 FL alone or in the presence of unlabelled 0.1 mM ALG2, 1 mM CALC, or 1mM Halo. The corresponding quantification of the % fluorescence recovery at 150 sec is plotted alongside. Mean ± SD. One-way ANOVA with Dunnett’s multiple comparison, *p < 0.05, ****p < 0.0001, ns - not significant (p > 0.05), n=3. N.B. The ANXA11 FL dataset displayed here is the same as in Figure S3C as experiments were performed concurrently. **(F)** Second derivative FTIR spectra of the Amide I protein IR absorption region. Spectra collected are from 150 µM ANXA11 FL alone (left) or coincubated with 600 µM ALG2 (right) (1[ANXA11]:4[ALG2]). The three dashed lines mark β-turn (left), α-helix and random coil (middle), and intermolecular parallel β-sheet (right) peak positions. **(G)** Quantification of the % composition of ANXA11 secondary structure (Mean ± SD). Note the low % of intermolecular parallel beta sheet, even after addition of ALG2. 19 spectra were collected across 5 experimental repeats. N.B. The FL dataset here is the same as in Figure 3D as the experiments were conducted concurrently.

**Figure S5.**
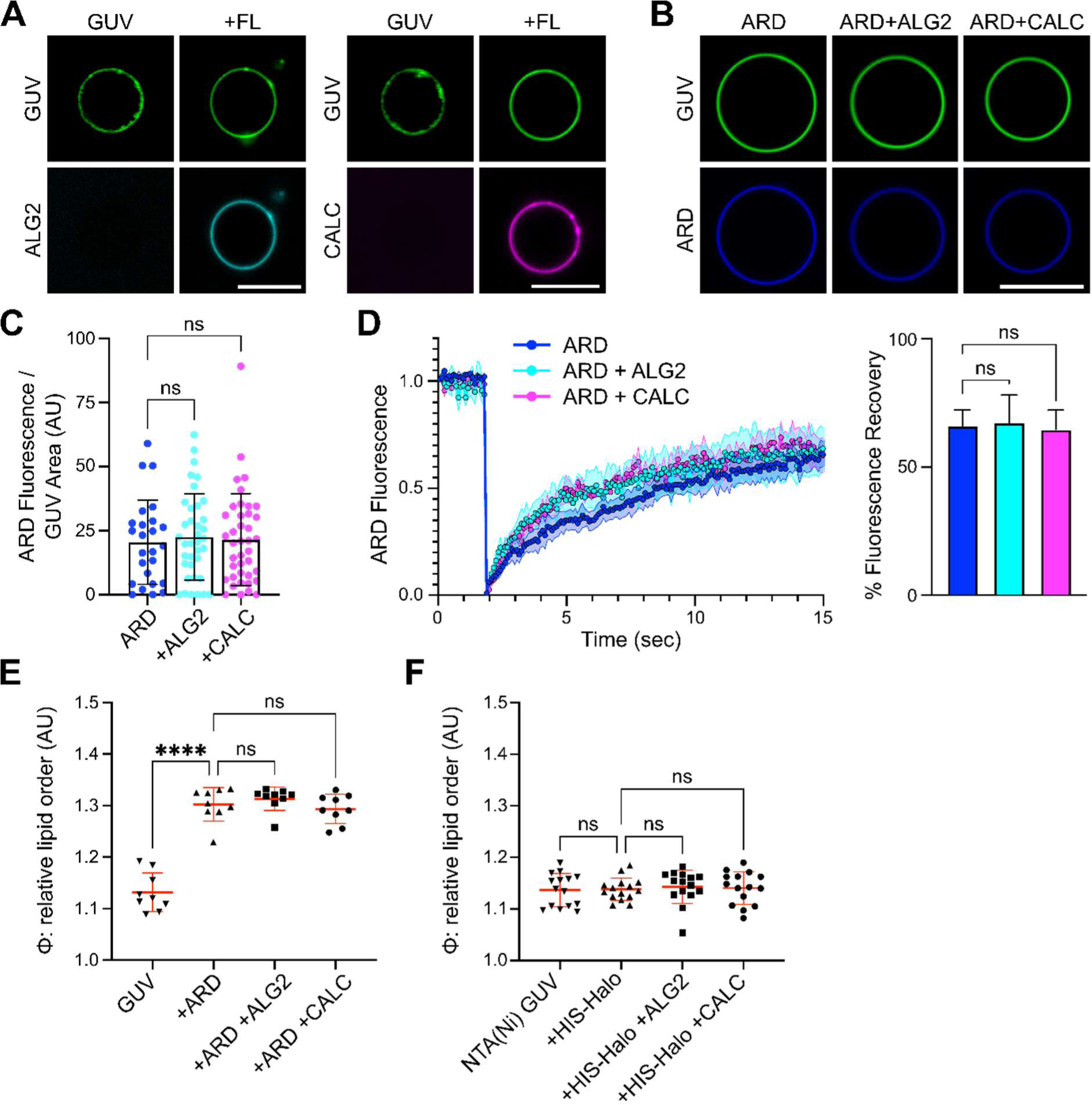
Effects of ALG2 and CALC on lipid membranes. **(A)** Fluorescence images of ATTO647 GUVs incubated with 100 µM Ca^2+^ and AF488-labelled 2 µM ALG2 or 20 µM CALC in the presence or absence of unlabelled 0.5 µM ANXA11 FL. Scale bar - 5 µm. **(B)** Representative fluorescence micrographs of ATTO488 GUVs incubated with 100 µM Ca^2+^ and 0.5 µM AF647-ARD, with and without unlabelled 2 µM ALG2 or 20 µM CALC. Scale bar – 5 µm. **(C)** Quantification of ARD recruitment to GUVs as displayed in (B). Mean ± SD. Kruskal-Wallis test with Dunn’s multiple comparison, ns - not significant (p > 0.05) n=3 (25-40 GUVs). **(D)** FRAP recovery curves of 0.5 µM AF647-ARD on the surface of GUVs in the presence or absence of 2 µM ALG2 or 20 µM CALC at 100 µM Ca^2+^. The corresponding quantification of the % fluorescence recovery at 150 sec is plotted alongside. Mean ± SD. One-way ANOVA with Dunnett’s multiple comparison, ns - not significant (p > 0.05), n=3. N.B. The ARD condition is extracted from the same dataset used in Figure 3C. **(E)** Quantification of the relative lipid order (φ) of ATTO647 GUVs labelled with 5 µM PK dye incubated with 0.5 µM ARD and at 100 µM Ca^2+^ in the presence or absence of 2 µM ALG2 or 20 µM CALC. Mean ± SD. One-way ANOVA with Tukey’s multiple comparison, ****p < 0.0001, ns - not significant (p > 0.05) n=4 (55-113 GUVs). **(F)** Quantification of the relative lipid order (φ) of NTA(Ni)-GUVs labelled with 5 µM PK dye. NTA(Ni) GUVs were incubated with 25 µM HIS-Halo and with either 0.1 mM ALG2 or 1 mM CALC. Mean ± SD. One-way ANOVA with Tukey’s multiple comparison, ns - not significant (p > 0.05), n=5 (17-64 GUVs).

**Figure S6.**
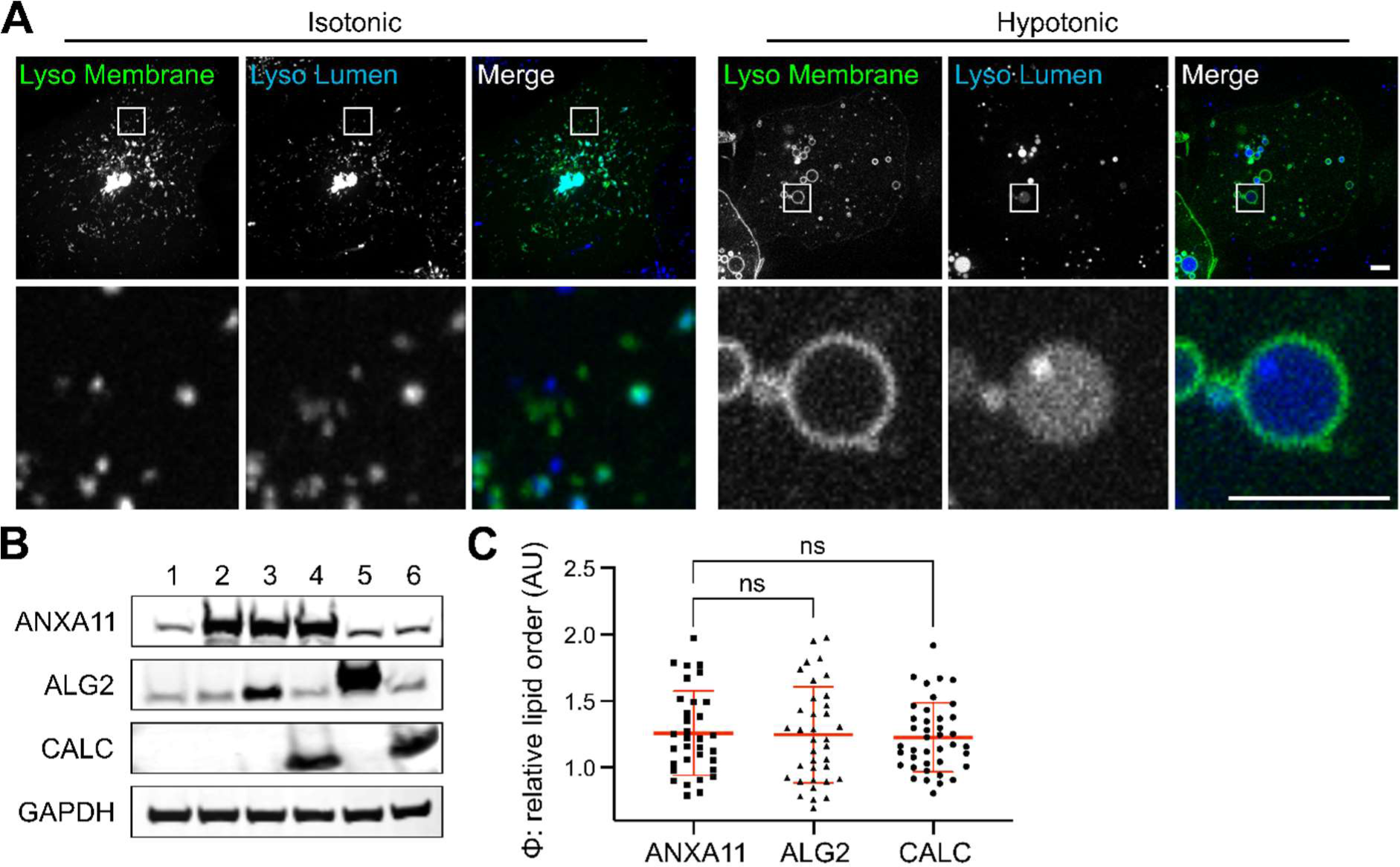
An *in cellula* system for studying the effects of ALG2 and CALC. **(A)** U2OS cells with fluorescently labelled lysosomal lumen (10-kDa dextran-AF594) and membrane (LAMP1-YFP) compartments, incubated in isotonic and hypotonic buffers (see STAR methods). The white ROI indicates zoomed regions displayed in the bottom row. Scale bars - 2 µm. **(B)** A Western blot of lysates from U2OS cells expressing the following constructs used to modulate ANXA11, ALG2 and CALC levels inside cells: (1) Untransfected; (2)ANXA11-T2A-TMEM192-Halo; (3)ANXA11-T2A-TMEM192-Halo/hPGK-ALG2; (4)ANXA11-T2A-TMEM192-Halo/hPGK-CALC; (5)ALG2-T2A-TMEM192-Halo; (6)CALC-T2A-TMEM192-Halo. Blots were probed with antibodies against ANXA11, ALG2, CALC and GAPDH. **(C)** Quantification of the relative lipid order (φ) of U2OS lysosomal membranes labelled with TMEM192-Halo-JF647 and incubated in 100 nM PK dye. Cells were expressing either ANXA11, or ALG2, or CALC. Mean ± SD. One-way ANOVA with Dunnett’s multiple comparison, ns - not significant (p > 0.05), n=5 (33-40 lysosomes).

**Figure S7.**
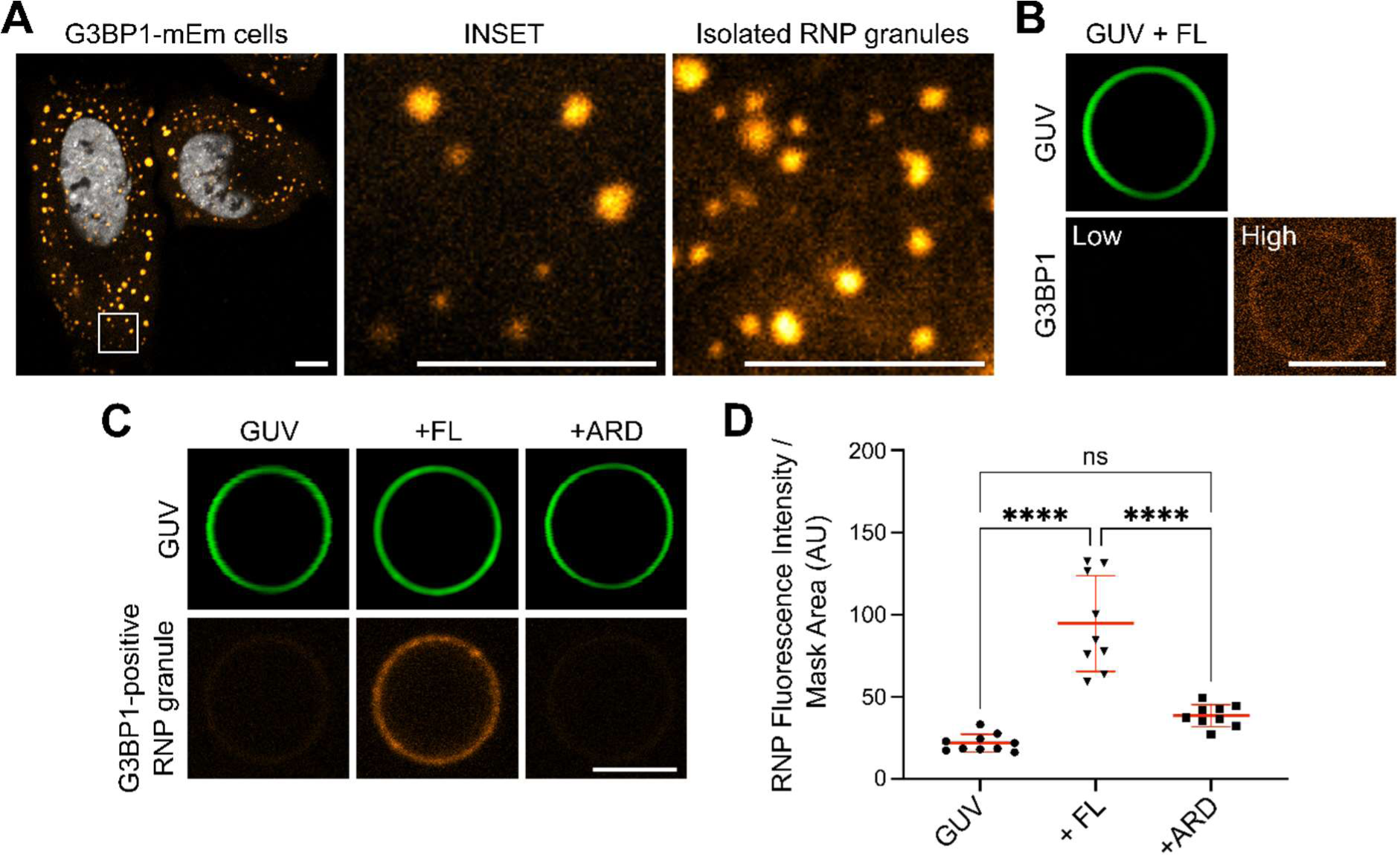
An *in vitro* RNP granule-ANXA11-GUV binding assay. **(A)** fluorescence images of a stable G3BP1-mEmerald (Orange) U2OS line labelled with after stress granule induction using 0.5 mM sodium arsenite for 30 min. SPY650-DNA was used to label the nucleus (white). On the right is an image of FAPS-isolated granules which have been pelleted and resuspended in a small volume of dilution buffer. Scale bar - 1.5 µm. **(B)** An ATTO594 GUV incubated with 100 µM Ca^2+^ and 0.5 µM ANXA11 FL and 0.2 mg/ml of recombinantly purified mEmerald-G3BP1 at a low and high imaging exposure. Scale bar - 5 µm. **(C)** Fluorescence images of ATTO594 GUVs incubated with 100 µM Ca^2+^ and either 0.5 µM ANXA11 FL or ARD. 0.2 mg/ml of G3BP1-mEmerald positive FAPS-isolated RNP granules were added to each condition. Scale bar - 5 µm. **(D)** Quantification of RNP granule recruitment to ANXA11-GUV assemblies as represented in (C). Mean ± SD. One-way ANOVA with Tukey’s multiple comparison, ****p < 0.0001, ns - not significant (p > 0.05), n=3 (40-106 GUVs).

**Movie 1** – A timelapse confocal movie of 50µM AF647-ANXA11 FL incubated at 37°C in order to visualise condensate foFrmation. A zoom panel is displayed inset. Scale bar – 10 µm.

**Table S1** – Proteins identified by mass spectrometry from FAPS-purified RNP granules (Figure 6) compared with a previously reported database from Youn et al., 2019.

## STAR Methods

### Plasmids and Cloning

Constructs encoding ANXA11 FL (aa 1-505), ANXA11 LCD (aa 1-185), ANXA11 ARD (aa 186-505), ALG2, CALC and G3BP1 were cloned by PCR amplification from HeLa cell cDNA into the bacterial expression vector pOPINS (Merck) containing a ULP protease cleavable N-terminal His-SUMO Tag. For mammalian cell line work, ANXA11 FL and TMEM192-Halo were inserted either side of a T2A self-cleaving peptide in a custom CMV-driven polycistronic vector (ANXA11-T2A-TMEM192-Halo). Where co-expression of ALG2 or CALC was required, either gene was inserted into an alternative ORF in the same vector with expression driven using a hPGK promoter. For experiments requiring ALG2 or CALC expression in the absence of ANXA11, the ANXA11 cassette in ANXA11-T2A-TMEM192-Halo was replaced by either ALG2 or CALC. G3BP1-mEmerald was created by subcloning the G3BP1 cassette from the pOPINS-G3BP1 construct into an mEm-N1 backbone derived from the Clontech N1 system. Cloning was performed throughout by PCR amplification using a Q5 polymerase system (NEB) and Gibson Assembly (NEB).

### Expression and Purification of Proteins

His-SUMO tagged pOPINS constructs (His-SUMO-ANXA11 FL, -LCD, -ARD, -ALG2,-CALC -G3BP1) were transformed into competent *E. coli* BL21(DE3) (NEB) and expressed overnight at 25 °C in TB autoinduction media. Cells were harvested by centrifugation and the cell lysate for protein isolation was produced by subjecting the harvested cells to a high-pressure cell disruption system (Constant systems). Prior to loading on a Ni-Sepharose Advance column (Bioserve), the cell lysate was clarified by high-speed ultracentrifugation at 100,000 g to remove the cell debris. A standard Ni-affinity protein purification protocol was followed and the column eluates containing the protein were pooled, mixed with ULP protease to remove the His-SUMO tags, and dialysed in dilution buffer (25 mM HEPES pH 7.4, 225 mM NaCl) supplemented with 5% (w/v) glycerol. After the cleavage, protein was applied on a second Ni-Sepharose Advance column to remove the His-SUMO tag, followed by size-exclusion chromatography using a Superdex S-75 column (Cytiva) in the same buffer. Purified protein fractions were pooled and concentrated in a spin concentrator (Vivapsin), aliquoted, snap frozen in liquid nitrogen and stored at –80 °C for subsequent use.

Halo protein purification was described previously (Chambers et al., 2018). Protein purity was routinely assessed by SDS-PAGE and all proteins were purified at or above 95% purity level (Figure S4A).

### Alexa Fluor Protein Labelling

Alexa Fluor AF647, AF555 and AF488 dyes (Thermo Scientific) were conjugated to proteins using NHS-ester chemistry following standard protocols. Briefly, 2.0 mg/ml of the protein was mixed with 100 µg of the dye and incubated stirring at 4 °C for 4 hours. Following the incubation, proteins were run on a Superdex-25 column (Cytiva) to separate the free dye from the conjugated protein-dye complex.

### NMR Spectroscopy

Isotopically enriched proteins were expressed and purified as described above. ANXA11 FL was expressed with ^15^N enrichment and AXNA11 LCD was expressed with either ^13^C/^15^N or ^13^C/^15^N/^2^H enrichment for residue specific assignments.

All NMR samples were collected in 25 mM HEPES pH 7.4, 225 mM NaCl with the addition of 2-5% D_2_O for sample locking. All spectra were collected on a 700MHz Bruker Avance II+ spectrometer equipped with a 5mm TCI cryoprobe using Topspin 3.6.0 acquisition software. ^1^H-^15^N BEST-TROSY spectra (Favier and Brutscher, 2011) were collected at 278K and 308K for both the 200 μM ^15^N-ANXA11 FL and 200 μM ^15^N-ANXA11 LCD.

A partial assignment of backbone H_N_, N_H_ and Cα, Cβ, C’ resonances of the 150 µM ^13^C/^15^N/^2^H ANXA11 LCD sample was based on the following six 3D BEST-TROSY datasets at 278K, acquired as pairs to provide own and preceding carbon connectivities. B_trHNCO, b_trHN(CA)CO, b_trHN(CO)CA, b_trHNCA, b_trHN(CO)CACB and b_trHNCACB were obtained as experimental pairs with 1024 complex points in the proton, and 80, 128 complex points in the ^15^N and ^13^C dimensions, respectively. Data sets were collected with 10-50 % Non Uniform Sampling (NUS) to reduce acquisition times.

The assignment of a subset of proline residues benefitted from C’-detect experiments collected with a 50 μM ^15^N/^13^C ANXA11 LCD sample. In these experiments correlations are made between the N_H_ and its preceding C’ carbon (N_i_ & C’_i-1_) and their own or preceding Cα_i,i-1_ or Cβ_i,i-1_ shifts. ^1^Hα-start, C’-detect sequences c_hcacon_ia, c_hcacon_ia3d, c_hcanco_ia3d and c_hcbcacon_ia3d were used for the N_i_-C’_i-1_, N_i_-C’_i-1_Cα_i-1_, N_i_-C’_i-1_Cα_i-1_ + N_i_-C’_i-1_-Cα_i_ and N_i_-C’_i-1_Cβ_i-1_ connectivities, respectively. Typical data sets consisted of 1024 complex points in the direct carbon, and 64, 128 complex points in the ^15^N and indirect ^13^C dimensions, respectively. C’-detect data was processed using Topspin (Bruker) and triple resonance NUS data was processed using NMRPipe (Delaglio et al., 1995) including NUS data reconstruction via compressed sensing (Kazimierczuk and Orekhov, 2011). Data analysis was completed using NMRFAM-Sparky, and the assignment tool MARS.

Secondary structure analysis was completed using TALOS-N where backbone torsion angles are extracted from secondary Cα, Cβ chemical shifts, i.e. shift differences between experimental data and those seen for the same residue under random coil conditions (Shen and Bax, 2013). Where assignments are missing, TALOS-N can predict secondary structure elements based on sequence alone.

### *In Vitro* Protein Condensation Assays

To visualise protein condensates *in vitro*, AF647-conjugated purified ANXA11 FL, LCD, ARD, and unlabelled purified ALG2 and CALC were made up in dilution buffer (25 mM HEPES pH 7.4, 225 mM NaCl) to the desired concentrations (listed in figures). Final volumes of 25 μL were transferred to 8-well glass bottom chambered coverslips (Ibidi, 80826) and incubated at 37 °C for 15-30 min before being imaged using a scanning confocal LSM 780 (Zeiss) fitted with a Plan-Apochromat 100× 1.4 NA oil immersion objective (Zeiss). An HeNe 5 mW 633 nm laser line was used for excitation (1 mW max at focal plane), with fluorescence collected on a GaAsP detector in the 620 nm – 750 nm λ range. Samples were imaged at 37 °C in a humidified imaging chamber with a Definite Focus module (Zeiss) employed for thermal drift correction and ZEN Black v2.3 (Zeiss) software used for acquisition.

### Generation of Phase Diagrams

Recombinant ANXA11 FL was mixed on ice in low-protein bind tubes (Eppendorf) with either LCD, ALG2, CALC, or HALO at various molar ratios (listed in figures). 20 µL of each sample was deposited into 8-well glass bottom chambered coverslips (Ibidi, 80826). Before imaging, samples were incubated at 25 °C for 15 min. Sample imaging was performed at 25 °C using Leica Stellaris 5 confocal microscope equipped with a Plan-Apochromat 63× 1.4 NA oil immersion objective (Leica). A Supercontinuum white light laser (440-790 nm) was used for excitation at 653 nm, with fluorescence collected on a Power HyD detector in the 660 nm – 750 nm λ range.

Confocal images were collected on LAS X (Leica) software and analysed using FIJI (NIH). The condensate detection threshold for the sample to be considered in the condensed state was defined by (i) the average area of condensates (>0.4 µm^2^), and (ii) the average number of condensates over a 90 µm^2^ region (>20 condensates). These parameters were established using the inbuilt particle analysis tool in FIJI.

The data was plotted using Python. The phase boundary was determined by fitting the data using a support-vector machine (SVM) algorithm with a linear or 2^nd^ degree polynomial kernel.

### ANXA11 Temperature Cycling

For temperature-cycling experiments (Figure S1C-D), AF647 labelled ANXA11 FL was imaged at 50 µM on a custom spinning disk microscope permitting precise and rapid control of sample temperature while allowing rapid imaging. The microscope comprised a Nikon Ti stand equipped with perfect focus, a fast piezo z-stage (ASI), a Plan Apochromat lambda 100X NA 1.45 objective, a Yokogawa CSU-X1 spinning disk head and a Photometrics 95B back-illuminated sCMOS camera. The camera operated in global shutter mode and was synchronized with the spinning disk rotation. Excitation was performed using 637 nm (140 mW OBIS LX) laser fibered within a Cairn laser launch, and a single band emission filter was used (Chroma 655LP). To enable fast acquisition, the entire setup was synchronized at the hardware level by a Zynq-7020 Field Programmable Gate Array (FPGA) stand-alone card (National Instrument sbRIO 9637) running custom code. In particular, fast z-stacks were obtained by synchronizing the motion of the piezo z-stage during the readout time of the cameras. Instrument was controlled by Metamorph software.

For control of temperature, a Cherry Temp stage was used (Cherry Biotech), which allows rapid temperature changes (∼5s) between two set temperatures (Velve-Casquillas et al., 2010). The sample was mounted within a chamber formed between a glass coverslip and the Cherry Temp microfluidics chip with a 0.5 mm PDMS insert in between. The sample was cycled through 6 cycles of increasing temperature (5°C increments) for one minute at a time, interspersed with one-minute periods at 5°C (to dissolve the condensates). Z-stacks were acquired (ΔZ = 200 nm), and maximum intensity z-projections were computed.

The extent of condensate formation (“granulosity index”), was then evaluated as follows: Raw images were processed for homogenous background subtraction, then a Fast Fourier Transform (FFT) was computed and a high-pass filter was applied via a circular mask prior to an inverse FFT. This mask was kept constant for all images (all source data were cropped to have the same size). We then computed the granulosity index as the ratio between the standard deviation and the mean of the signal in the high pass filtered image.

### GUV & SUV Preparation

All lipids were purchased from Avanti Polar Lipids (Alabama, USA). GUVs and SUVs were prepared as described previously (Liao et al., 2019).

Briefly, giant unilamellar vesicles (GUVs) were prepared (w/v) from 50% 1-palmitoyl-2-oleoyl-3-phosphocholine (POPC, Avanti 850457), 20% 1-palmitoyl-2-oleoyl-sn-glycero-3-phosphoethanolamine (POPE, Avanti 850757), 10% 1-stearoyl-2-arachidonoyl-sn-glycero-3-phosphoinositol (SAPI, Avanti 850144), 10% 1-palmitoyl-2-oleoyl-sn-glycero-3-phospho-L-serine (POPS, Avanti 840034), 5% 1,2-dioleoyl-sn-glycero-3-phospho-(1’-myo-inositol-3’-phosphate) (PI(3)P, Avanti 850150), 0.05% ATTO488, ATTO594 or ATTO647 labelled 1,2-dioleoyl-sn-glycero-3-phosphoethanolamine (DOPE, Sigma), and 4.95% cholesterol (Avanti 700000) dissolved in chloroform. In certain experiments, 10% 1,2-dioleoyl-sn-glycero-3-[(N-(5-amino-1-carboxypentyl)iminodiacetic acid)succinyl] (nickel salt) (DGS-NTA(Ni), Avanti 790404) was added at the expense of 10% POPC. GUVs were formed via electroformation using a Nanoion Vesicle Prep Pro system (Nanion Technologies, Germany) according to the manufacturer’s instructions. Briefly, lipids were dispensed using a Hamilton syringe (Sigma) onto a conductive slide under nitrogen vapour. The chloroform was evaporated from the slide overnight, again under nitrogen vapour, in a dark sealed chamber. The following day, a greased O ring was placed over the lipid film, with lipids then rehydrated in 250 µl of liposome buffer (50 mM HEPES pH 7.4, 450 mM Sucrose). The slide was loaded onto the Vesicle Prep Pro with the following program details run: Frequency - 10, Amplitude - 1.4, Temperature - 60 °C, Rise - 3 min. Formed GUVs were then removed from the slide and used for future applications.

Small unilamellar vesicles (SUVs) for microfluidic assays were prepared using the same lipid composition as GUVs above. A dry and thin lipid film was prepared by gently evaporating the chloroform using a nitrogen stream. The film was then placed under vacuum overnight to remove all traces of chloroform. Liposome buffer (50 mM HEPES pH 7.4, 450 mM Sucrose) was added to hydrate the lipid film to the concentration of 800 µg/mL and the solution was stirred at room temperature for at least 2 hr. The lipid solution was then frozen and thawed 8 times using liquid nitrogen and a water bath set to 37 °C. The solution was then sonicated on ice with a probe sonicator (3 x 5 minutes, 50% cycles, 20% maximum power) and centrifuged for 30 min at 15,000 rpm to remove probe residues. The size of SUVs was checked with Dynamic Light Scattering (Zetasizer, Malvern Panalytical, UK). The radius of lipid vesicles was measured to be 18.65 +/- 2.5 nm (9 batches of SUVs produced independently). SUV solution was stored at 4 °C and diluted to 400 µg/mL for the diffusional sizing assay.

### Microfluidic Diffusional Sizing Assays

Microfluidic devices were fabricated using standard soft-photolitography techniques by using a silicone rubber compound (Momentive RTV615, Techsil) mixed with black carbon powder (PL-CB13, PlasmaChem GmbH) to minimise the background signal in fluorescence images. The channel height used was between 25 and 50 µm, with the specific height of each device measured using a Dektak profilometer (Bruker). Upon binding to a glass slide, the devices were filled with dilution buffer (25 mM HEPES pH 7.4, 225 mM NaCl) containing 0.01% v/v Tween 20 (Sigma) at least 1 hr prior to the experiment, to prevent protein adhesion to the channel surfaces.

Microfluidic diffusional sizing experiments were performed as described previously (Arosio et al., 2016; Schneider et al., 2021) using the chip design as displayed in Figure S1F. Prior to the sizing experiment, all samples were incubated for 1 hr at 37 °C. Proteins were used at 0.5 µM (ANXA11 FL, LCD, ARD), and SUVs at 400 µg/mL in dilution buffer (25 mM HEPES pH 7.4, 225 mM NaCl). Calcium chloride (Sigma; henceforth Ca^2+^) was used at 50, 100 and 500 µM. The sample and buffer were loaded at the inlets, with liquid withdrawn from the outlet using a glass syringe (Hamilton) mounted to a neMESYS syringe pump (Cetoni GmbH). The sample was co-flown with the dilution buffer (also containing appropriate Ca^2+^ concentrations) along the channel at 3-4 different flow rates ranging from 20 to 150 µL/hr. For each flow rate, once stabilisation of flow was achieved, fluorescent images were taken of 4 fluidic channels of differing path length corresponding to diffusion times of SUV-protein complexes. The images were obtained with either an inverted Axio Observer A1 microscope (Zeiss) equipped with ET-EGFP (49002, Chroma), ET-CY3/R (49004, Chroma), excitation 628/40 nm, emission 692/40 nm, dichroic mirror 660 nm (Cy5-4040C-000, Semrock) a 5× 0.12 NA A-Plan objective (Zeiss) and an Evolve 512 CCD camera (Photometrics).

The images were analysed with custom written Python scripts. First, the fluorescence profiles were extracted from the images to determine the spatial distribution of sample molecules as a function of time. Diffusion profiles corresponding to the diffusion coefficients for R_H_ = 0.1-50 nm, and the best fit to the observed sample distribution at the four time points was used to determine the average R_H_ for each measurement (Müller et al., 2016; Zhang et al., 2020).

### Imaging and Quantification of Protein Recruitment to GUVs

To determine the relative recruitment levels of ANXA11 FL, LCD and ARD to GUVs at different Ca^2+^ concentrations, purified AF647-labelled protein was added at a 0.5 μM final concentration to ATTO488-GUVs made up in dilution buffer (25 mM HEPES pH 7.4, 225 mM NaCl). After addition of the desired concentration of Ca^2+^ (listed in figures), 200 μL of the protein-GUV mixture was plated into 8-well glass bottom chambered coverslips (Ibidi, 80826) and incubated at 37 °C for 15-30 mins prior to imaging.

To examine the direct recruitment of AF488-conjugated ALG2 and CALC to GUVs, 2 μM ALG2 or 20 μM CALC were incubated in the presence of 100 μM Ca^2+^ with ATTO647 GUVs alone, or co-incubated with 0.5 μM ANXA11 FL (unlabelled) (Figure S5A). To determine the influence of ALG2 and CALC on ANXA11 FL recruitment to GUVs the same experiment was performed using ATTO488-GUVs, AF647-labelled ANXA11 FL and unlabelled ALG2 and CALC. To quantify the relative abundance of protein recruited to the GUV surface, confocal microscopy was carried out using an LSM 780 (Zeiss). Excitation was performed sequentially using an Argon multi-line 35 mW 488 nm laser (3 mW max at focal plane) and HeNe 5 mW 633 nm laser (1 mW max at focal plane), with imaging conditions experimentally selected to minimise crosstalk. The resulting fluorescence was collected using a 63× Plan-Apochromat 1.4 NA oil immersion objective (Zeiss) and detected on a 34-channel spectral array detector in the 460 – 600 nm and 620 – 750 nm λ range. Samples were imaged at 37 °C in a humidified imaging chamber with a Definite Focus module (Zeiss) employed for thermal drift correction and ZEN Black v2.3 (Zeiss) software used for acquisition. To measure protein recruitment to GUVs, data was analysed in FIJI (NIH) using a custom macro. The GUV channel was initially gaussian blurred (sigma radius 1.5) to reduce detector noise, thresholded by intensity, and masked. The integrated fluorescence intensity of the protein channel within the GUV-mask was calculated and normalized to the mask area.

### Fluorescence recovery after photobleaching (FRAP)

To observe the mobility of ANXA11 FL and ARD on the surface of GUVs, in the presence or absence of 25 μM LCD, or 2 μM ALG2, or 20 μM CALC, FRAP of 0.5 μM AF647-conjugated ANXA11 FL and ARD was performed. Labelled FL or ARD were added to 20 μM ATTO488 GUVs in dilution buffer (25 mM HEPES pH 7.4, 225 mM NaCl) in 100 μM Ca^2+^. 200 μL of the protein-GUV mixture was then plated into 8-well glass bottom chambered coverslips (Ibidi, 80826) and incubated at 37 °C for 15-30 minutes prior to imaging.

To test the effect of ANXA11 on lipid mobility, 20 μM GUVs in which the PI(3)P was replaced with fluorescent BODIPY-PI(3)P (Echelon) were incubated with or without 0.5 μM AF647-labelled ANXA11 FL for 15-30 min at 37 °C in 100 μM Ca^2+^ as described above.

FRAP experiments were performed on a scanning confocal LSM 780 (Zeiss) fitted with a Plan-Apochromat 100× 1.4 NA oil immersion objective (Zeiss) at 37 °C in a humidified imaging chamber. An Argon multi-line 35 mW 488 nm laser (3 mW max at focal plane) and HeNe 5 mW 633 nm laser (1 mW max at focal plane) were used for excitation and bleaching of GUVs and AF647-labelled protein respectively. Fluorescence was detected on a 34-channel spectral array detector in the 460 – 600 nm (GUV) and 620 – 750 nm (protein) λ range. Samples were stabilised for thermal drift using a Definite Focus module (Zeiss) and ZEN Black v2.3 (Zeiss) software was used for acquisition. Images were collected at 11 Hz. FRAP data were plotted using the FRAP Profiler plugin on FIJI (NIH).

For FRAP studies on condensates, AF647-ANXA11 FL was co-incubated with either recombinant LCD, or ALG2, or CALC, or Halo (see figures for concentrations). Imaging conditions were identical to protein FRAP on the surface of GUVs but with the FRAP region drawn internally within a condensate, and imaging performed at 1Hz.

### *In Vitro* Membrane Lipid Order Assays

To assess lipid membrane order using a solvatochromic pyrene probe (PK dye), GUV and protein coincubations were performed as described above (Imaging and Quantification of Protein Recruitment to GUVs), with the exception that DOPE within the GUVs was labelled with ATTO647 and the protein components were unlabelled. Specific protein, GUV and PK dye concentrations are detailed in the figure legends.

Imaging was performed on a scanning confocal LSM 780 (Zeiss) fitted with a Plan-Apochromat 100× 1.4 NA oil immersion objective (Zeiss) at 37 °C in a humidified imaging chamber. A 30 mW continuous wave 405 nm diode pumped solid state (DPSS) laser (<3 mW max at focal plane) and HeNe 5 mW 633 nm laser (1 mW max at focal plane) were used for excitation of the PK dye and GUVs respectively. Fluorescence was detected on a 34-channel spectral array detector across three spectral windows: 449 – 550 nm (BFP/GFP), 550 – 650 nm (RFP), and 650 – 750 nm (Cy5). Samples were stabilised for thermal drift using a Definite Focus module (Zeiss) and ZEN Black v2.3 (Zeiss) software was used for acquisition.

The Cy5 emission window was used to identify ATTO647 GUVs, with the ratio of the PK dye fluorescence from the BFP/GFP and RFP signals used to calculate the relative lipid order (φ) in FIJI (NIH), as reported in Valanciunaite et al., 2020. To minimise the contribution of low signal background pixels, a mask was generated from the GUV channel, with regions outside of the mask in the BFP/GFP/RFP channels set to a grey value of 0.

### Elastic Modulus

The relative elastic modulus of GUVs was measured in a two-layered microfluidic platform with the channel design shown in Figure 6A. A rectangular-shaped main channel with height 25 μm was connected by 14 V-shaped tapered small bridges with height 4 μm. The microfluidic system was fabricated on the silicon wafer by exposing once with a film mask and another time with a chrome mask to obtain channels with different heights. Once the GUVs were loaded and trapped in the small V-shaped bridges, different pressure drops (100 - 1000 Pa) were applied across the bridges by tuning the flow rate of the buffer as controlled by a neMESYS syringe pump (Cetoni). The relationship between the pressure drop across the V-shaped channels and the elongation of the vesicle reveals the elastic property of the GUV. Relative elastic moduli were calculated as described previously (Xu et al., 2022). Measurements for the deformation of the GUV were obtained using an AxioObserver inverted microscope (Zeiss) coupled to an Evolve 512 CCD camera (Photometrics). The images were analysed on FIJI (NIH).

### Infrared Nanospectroscopy (AFM-IR)

To perform the AFM-IR spectroscopy, we deposited ANXA11-GUV assemblies (prepared as described above) onto hydrophobic ZnSe surfaces. Our initial AFM survey revealed that these assemblies are heterogeneous, forming micron-sized fragments upon deposition. Some contained only GUV lipid fragments, with no protein present. Other assemblies contained GUV lipid fragments bound to ANXA11. We thus exploited this heterogeneity to compare, in the same sample, the morphological, and chemical properties of GUV lipids without protein versus GUV lipids coated with protein.

We used a nanoIR2 platform (Anasys), which combines high resolution and low-noise AFM with a tunable quantum cascade laser (QCL) with top illumination configuration (Ruggeri et al., 2016; Simone Ruggeri et al., 2016). The sample morphology was scanned by the nanoIR system, with a line rate within 0.1-0.4 Hz and in contact mode. A silicon gold coated PR-EX-nIR2 (Anasys) cantilever with a nominal radius of 30 nm and an elastic constant of about 0.2 N m^-1^ was used.

Both infrared (IR) spectra and images were acquired by using phase loop (PLL) tracking of contact resonance, the phase was zeroed to the desired off-resonant frequency on the left of the IR amplitude maximum and tracked with an integral gain I=0.1 and proportional gain P=5 (Ramer et al., 2018; Ruggeri et al., 2018). All images were acquired with a resolution between 1000×500 and 1000×100 pixels per line.

The AFM morphology maps were treated and analysed using SPIP software. The height images were first order flattened, while IR maps were flattened by a zero-order algorithm (offset). Nanoscale-localised spectra were collected by placing the AFM tip on the GUV fragments. A laser wavelength of 2 cm^-1^ and a spectral speed of 100 cm^-1^/s within the range 1400-1794 cm^1^ were used for sampling. Within a single GUV fragment, the spectra were acquired at multiple nanoscale localised positions. Each spectrum collected was averaged 5 times. Successively, the spectra were smoothed by an adjacent averaging filter (3pts) and a Savitzky-Golay filter (second order, 13 points) and normalised. Spectra second derivatives were calculated, smoothed by a Savitzky-Golay filter (second order, 13 points).

GUV fragments containing lipids only possessed typical C=O peaks for phospholipids and cholesterol esters at 1730-32 cm^-1^ (with a spectral width of ∼20 cm^-1^ by second derivative analysis) and CH_2_ group peaks at 1450-1475 cm^-1^ (Mantsch and McElhaney, 1991; Tamm and Tatulian, 1997).

The secondary structural organisation of protein components was evaluated by integrating the area of the different secondary structural contributions in the amide band I, as previously shown (Ruggeri et al., 2015b, 2020). The error here was calculated over the average of at least 5 independent spectra. Spectra were analysed using the microscope’s built-in Analysis Studio (Bruker) and OriginPRO (OriginLab). All measurements were performed at room temperature, with laser powers between 0.1-0.5 mW and under controlled Nitrogen atmosphere with residual real humidity below 5%.

### Fourier Transform Infrared Spectroscopy (FTIR)

Attenuated total reflection Fourier transform infrared spectroscopy (FTIR) was performed using a Bruker Vertex 70 spectrometer equipped with a diamond ATR element. The resolution was 4 cm^-1^ and all spectra were processed using OriginPRO software (OriginLab). The spectra were averaged (3-19 spectra with 256 co-averages), smoothed applying a Savitzky-Golay filter (2^nd^ order, 9 points) and then the second derivative was calculated applying a Savitzky-Golay filter (2^nd^ order, 11 points).

### Cell Culture and Treatments

U2OS cells were cultured in Dulbecco’s Modified Eagle Medium (DMEM, Gibco) with 10% fetal bovine serum (FBS, Gibco) and seeded in triplicate into Matrigel-coated 8-well glass bottom chambered coverslips (Ibidi, 80826). At ∼70% cell confluency, cells were transiently transfected using Lipofectamine 3000 (ThermoFisher) and 1 μg of the indicated plasmid according to the manufacturer’s instructions. Cells were subsequently imaged ∼24 hr after transient transfection.

For the PK dye experiments, U2OS cells were transiently transfected with the plasmids indicated as described above. 24 hours after transfection, cells were washed two times with PBS. To label Halo-TMEM192 on lysosomal membranes, the JF646 Halo ligand (a kind gift from the Lavis Lab) was diluted to 1:2000 in optiMEM (Gibco) and added to the cells for 1 min at 37 °C and 5% CO_2_. Cells were rinsed three times with PBS and 100 nM PK dye diluted in DMEM with 10% FBS was applied to the cells for 10 min. Cells were then rinsed three times with PBS and hypotonic buffer (95% ddH_2_O, 5% DMEM, pH 7.0) was applied to cells. Cells were then incubated for 10 min at 37 °C and 5% CO_2_ to allow for lysosomal swelling. Cells were subsequently imaged for up to one hour post hypotonic treatment.

For fluorescent labelling of the lysosomal lumen, cells were loaded with fluorescent dextran utilising a three hour pulse of 0.5 mM 10kDa AF594-dextran (ThermoFisher). After 5x washes with PBS, a chase was performed for three hours in full DMEM with 10% FBS. Cells were then treated with hypotonic media as above and imaged.

### Live cell microscopy PK dye analysis

Live cell microscopy was performed with a customized Nikon TiE inverted microscope utilizing a Yokogawa spinning disk scan head (CSU-X1, Yokogawa) and a Prime BSI sCMOS camera (Photometrics). Cell fluorescence was collected with 525/36m (GFP), 605/70m (RFP) and 700/75m (Cy5) filter sets (Chroma) through a 100× Plan-Apochromat 1.4 NA oil immersion objective (Nikon). Cells were maintained at 37 °C and 5% CO_2_ with a Tokai Hit stage-top incubator. For experiments interrogating lysosomal lipid order, PK dye signal was imaged with 405 nm laser excitation. Data were acquired in two sequential scans through the GFP and RFP filter sets, with a 200 ms exposure time.

Images from experimental replicates were pooled and regions of intracellular lysosome membranes with TMEM192 signal were selected from cells. The ratio of the PK dye fluorescence from the GFP and RFP signals for the regions was calculated after subtraction of background signal. Background signal was calculated for each image by selecting a region of the image plane with no membrane fluorescence in the PK dye channels and averaging the counts. Region selections were performed with FIJI (NIH) and ratios were calculated in Microsoft Excel. It is important to note that exposure to hypotonic conditions likely alters the lipid ordering of membranes and so results should be interpreted comparatively across conditions, and not as absolute measurements.

### Cell free RNP granule-lipid binding assay

RNP granules were isolated from U2OS cells stably expressing G3BP1-mEmerald using fluorescence-activated particle sorting (FAPS). FAPS-based RNP granule isolation has been described in detail previously (Hubstenberger et al., 2017). Briefly, stress granules, a class of RNP granule, were induced in cells with 0.5mM sodium arsenite (sigma) for 30 min prior to lysis. Cell pellets were then suspended in lysis buffer (50 mM Tris-HCl, 1 mM EDTA, 150 mM NaCl, 1% NP-40, pH 7.4) with freshly added cOmplete EDTA-free protease inhibitor cocktail (Roche, Merck) and 80 U/mL RNaseOut ribonuclease inhibitor (Promega). Nuclei and cell debris were cleared from the lysate by centrifugation at 200xg for 5 min. Supernatants were further spun at 10,000 g for 7min, and then pellets were resuspended into lysis buffer in the presence of 80 Units of RNaseOut (Promega). Particles in the pellets were sorted on a High-Speed Influx Cell Sorter (BD Biosciences). Stress granules were isolated based on their size and fluorescence, as detected with 488 nm excitation. To analyse protein composition of the isolated stress granules, the sample was firstly digested by trypsin, and then further processed and analysed using LC-MS/MS by the CIMR proteomics facility (Cambridge Institute for Medical Research, University of Cambridge, UK). To verify that the isolated particles were indeed stress granules, we compared our proteomic data with an RNP granule proteomic database (Table S1) (Youn et al., 2019).

To measure the association of purified RNP granules with ANXA11-GUV assemblies in the presence and absence of ALG2 and CALC, proteins and ATTO594 GUVs were prepared as above, with specific concentrations listed within the figure legends. 0.2mg/ml of FAPS-isolated RNP granules were then added to each sample. For quantification, the total fluorescence intensity of mEmerald-G3BP1 labelled RNP granules was integrated across and normalised to GUV mask area as described above (Imaging and Quantification of Protein Recruitment to GUVs).

### Overexpression construct validation

Overexpression constructs were designed to increase the levels of ANXA11, ALG2 and CALC in living cells (see Plasmids and Cloning section). To validate these plasmids, they were transfected in U2OS cells as described above (Cell Culture and Treatments). After ∼24 hr, cells were scraped and incubated at 4 °C in ice cold lysis buffer (25 mM TRIS pH 7.0, 150 mM NaCl, 1% NP40, 1mM EDTA, 5% glycerol) containing 1x protease inhibitor (Roche). After 30 min, lysates were spun at 10,000 *g* for 10 min and supernatants diluted into SDS-loading buffer. Samples were run on an SDS-PAGE 4-12% NuPAGE gel (ThermoFisher), transferred to a PVDF membrane (Millipore), blocked at RT for 1 hr in 5% BSA (Sigma) in 1x TBS-T, and probed overnight with antibodies against ANXA11 (10479-2-AP Proteintech), ALG2 (ab133326 Abcam), CALC (10245-1-AP Proteintech) and GAPDH (D16H11 Cell Signaling). Blots were thoroughly washed in TBS-T, labelled with fluorescent IRDye 680 (LiCor) secondary antibodies, washed again in TBS-T and imaged on an Odyssey CLx (LiCor).

### Image processing and display

Display images of GUVs throughout the manuscript are shown with an LUT that is baseline corrected to reduce the contribution of scattered light in the scope. A gaussian blur (sigma radius 1.5) has been applied after LUT application to remove detector noise and additionally supress background scattered light. Display PK dye ratios have been subjected to additional processing steps (detailed in *In Vitro* Membrane Lipid Order Assays). A gaussian blur (sigma radius 1.5) and LUT baseline correction was also applied to display images of cells in Figure 5 for the same reasons as above. In some display images linear compression of the LUT results in saturation of certain regions within a field of view. This is necessary to highlight regions of lower signal intensity to the reader.

### Statistics

Where possible, a series of normality tests (Anderson-Darling, D’Agostino & Pearson, Shapiro-Wilk, Kolmogorov-Smirnov) were performed to determine if parametric or non-parametric approaches were appropriate. Certain datasets were determined to be non-normally distributed. This is not surprising given that the fluorescence detection range of photomultiplier tubes (PMTs) on confocal microscopes is limited. As such, a dynamic range was selected to prevent pixel saturation at the expense of detecting low fluorescence signal events, thereby skewing the detection window (sampling) towards the upper tail of the distribution. In these cases, a nonparametric Kruskal-Wallis test was performed with Dunn’s multiple comparison to compare datasets. Further details are summarised below, with P-values provided within the figure legends.

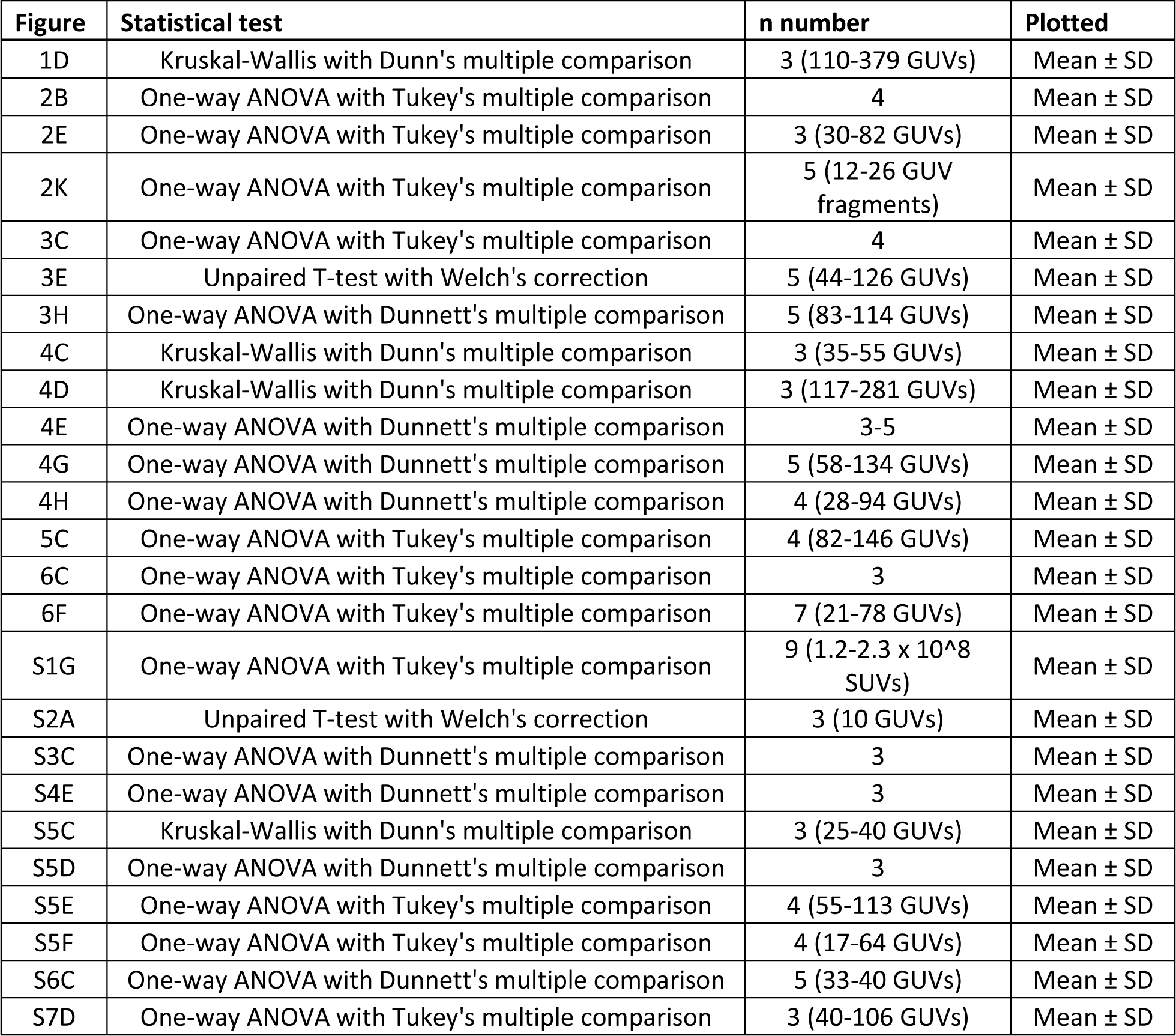

## REFERENCES

Aman, M.J., and Ravichandran, K.S. (2000). A requirement for lipid rafts in B cell receptor induced Ca2+ flux. Curr. Biol. 10, 393–396.

Andes-Koback, M., and Keating, C.D. (2011). Complete budding and asymmetric division of primitive model cells to produce daughter vesicles with different interior and membrane compositions. J. Am. Chem. Soc. 133, 9545–9555.

Arosio, P., Müller, T., Rajah, L., Yates, E. V., Aprile, F.A., Zhang, Y., Cohen, S.I.A., White, D.A., Herling, T.W., De Genst, E.J., et al. (2016). Microfluidic diffusion analysis of the sizes and interactions of proteins under native solution conditions. ACS Nano 10, 333–341.

Banani, S.F., Lee, H.O., Hyman, A.A., and Rosen, M.K. (2017). Biomolecular condensates: Organizers of cellular biochemistry. Nat. Rev. Mol. Cell Biol.

Chambers, J.E., Kubánková, M., Huber, R.G., López-Duarte, I., Avezov, E., Bond, P.J., Marciniak, S.J., and Kuimova, M.K. (2018). An Optical Technique for Mapping Microviscosity Dynamics in Cellular Organelles. ACS Nano 12, 4398–4407.

Chen, W., Duša, F., Witos, J., Ruokonen, S.K., and Wiedmer, S.K. (2018). Determination of the Main Phase Transition Temperature of Phospholipids by Nanoplasmonic Sensing. Sci. Rep. 8, 1–11.

Chung, J.K., Huang, W.Y.C., Carbone, C.B., Nocka, L.M., Parikh, A.N., Vale, R.D., and Groves, J.T. (2021). Coupled membrane lipid miscibility and phosphotyrosine-driven protein condensation phase transitions. Biophys. J. 120, 1257–1265.

Delaglio, F., Grzesiek, S., Vuister, G.W., Zhu, G., Pfeifer, J., and Bax, A. (1995). NMRPipe: A multidimensional spectral processing system based on UNIX pipes. J. Biomol. NMR 6, 277– 293.

Díaz, P., Sandoval-Bórquez, A., Bravo-Sagua, R., Quest, A.F.G., and Lavandero, S. (2021). Perspectives on Organelle Interaction, Protein Dysregulation, and Cancer Disease. Front. Cell Dev. Biol. 9, 291.

Domon, M., Nasir, M.N., Matar, G., Pikula, S., Besson, F., and Bandorowicz-Pikula, J. (2012). Annexins as organizers of cholesterol- and sphingomyelin-enriched membrane microdomains in Niemann-Pick type C disease. Cell. Mol. Life Sci. 69, 1773–1785.

Eggeling, C., Ringemann, C., Medda, R., Schwarzmann, G., Sandhoff, K., Polyakova, S., Belov, V.N., Hein, B., Von Middendorff, C., Schönle, A., et al. (2009). Direct observation of the nanoscale dynamics of membrane lipids in a living cell. Nature 457, 1159–1162.

Favier, A., and Brutscher, B. (2011). Recovering lost magnetization: Polarization enhancement in biomolecular NMR. J. Biomol. NMR 49, 9–15.

Gokhale, N.A., Abraham, A., Digman, M.A., Gratton, E., and Cho, W. (2005). Phosphoinositide specificity of and mechanism of lipid domain formation by annexin A2-p11 heterotetramer. J. Biol. Chem. 280, 42831–42840.

Guillén-Samander, A., Leonzino, M., Hanna, M.G., Tang, N., Shen, H., and De Camilli, P. (2021). VPS13D bridges the ER to mitochondria and peroxisomes via Miro. J. Cell Biol. 220.

Heberle, F.A., and Feigenson, G.W. (2011). Phase separation in lipid membranes. Cold Spring Harb. Perspect. Biol. 3, 1–13.

Huang, L., Mollet, S., Souquere, S., Le Roy, F., Ernoult-Lange, M., Pierron, G., Dautry, F., and Weil, D. (2011). Mitochondria associate with P-bodies and modulate microRNA-mediated RNA interference. J. Biol. Chem. 286, 24219–24220.

Hubstenberger, A., Courel, M., Bénard, M., Souquere, S., Ernoult-Lange, M., Chouaib, R., Yi, Z., Morlot, J.B., Munier, A., Fradet, M., et al. (2017). P-Body Purification Reveals the Condensation of Repressed mRNA Regulons. Mol. Cell 68, 144–157.e5.

Itzhak, D.N., Tyanova, S., Cox, J., and Borner, G.H.H. (2016). Global, quantitative and dynamic mapping of protein subcellular localization. Elife 5.

Kazimierczuk, K., and Orekhov, V.Y. (2011). Accelerated NMR spectroscopy by using compressed sensing. Angew. Chemie - Int. Ed. 50, 5556–5559.

King, C., Sengupta, P., Seo, A.Y., and Lippincott-Schwartz, J. (2020). ER membranes exhibit phase behavior at sites of organelle contact. Proc. Natl. Acad. Sci. 201910854.

Lee, I.H., Imanaka, M.Y., Modahl, E.H., and Torres-Ocampo, A.P. (2019). Lipid Raft Phase Modulation by Membrane-Anchored Proteins with Inherent Phase Separation Properties. ACS Omega 4, 6551–6559.

Lee, J.E., Cathey, P.I., Wu, H., Parker, R., and Voeltz, G.K. (2020). Endoplasmic reticulum contact sites regulate the dynamics of membraneless organelles. Science (80-.). 367.

Levental, I., Levental, K.R., and Heberle, F.A. (2020). Lipid Rafts: Controversies Resolved, Mysteries Remain. Trends Cell Biol. 30, 341–353.

Liao, Y.C., Fernandopulle, M.S., Wang, G., Choi, H., Hao, L., Drerup, C.M., Patel, R., Qamar, S., Nixon-Abell, J., Shen, Y., et al. (2019). RNA Granules Hitchhike on Lysosomes for Long-Distance Transport, Using Annexin A 11 as a Molecular Tether. Cell 179, 147-164.e20.

Lin, Y.C., Chipot, C., and Scheuring, S. (2020). Annexin-V stabilizes membrane defects by inducing lipid phase transition. Nat. Commun. 11.

Ma, W., and Mayr, C. (2018). A Membraneless Organelle Associated with the Endoplasmic Reticulum Enables 3′UTR-Mediated Protein-Protein Interactions. Cell 175, 1492–1506.e19.

Mantsch, H.H., and McElhaney, R.N. (1991). Phospholipid phase transitions in model and biological membranes as studied by infrared spectroscopy. Chem. Phys. Lipids 57, 213–226.

Menke, M., Gerke, V., and Steinem, C. (2005). Phosphatidylserine membrane domain clustering induced by annexin A2/S100A10 heterotetramer. Biochemistry 44, 15296–15303.

Milovanovic, D., Wu, Y., Bian, X., and De Camilli, P. (2018). A liquid phase of synapsin and lipid vesicles. Science (80-.). 361, 604–607.

Müller, T., Arosio, P., Rajah, L., Cohen, S.I.A., Yates, E. V., Vendruscolo, M., Dobson, C.M., and Knowles, T.P.J. (2016). Particle-Based Monte-Carlo Simulations of Steady-State Mass Transport at Intermediate Péclet Numbers. Int. J. Nonlinear Sci. Numer. Simul. 17, 175–183.

Nesterov, S. V., Ilyinsky, N.S., and Uversky, V.N. (2021). Liquid-liquid phase separation as a common organizing principle of intracellular space and biomembranes providing dynamic adaptive responses. Biochim. Biophys. Acta - Mol. Cell Res. 1868, 119102.

Nicolini, C., Kraineva, J., Khurana, M., Periasamy, N., Funari, S.S., and Winter, R. (2006). Temperature and pressure effects on structural and conformational properties of POPC/SM/cholesterol model raft mixtures-a FT-IR, SAXS, DSC, PPC and Laurdan fluorescence spectroscopy study. Biochim. Biophys. Acta - Biomembr. 1758, 248–258.

Qamar, S., Wang, G.Z., Randle, S.J., Ruggeri, F.S., Varela, J.A., Lin, J.Q., Phillips, E.C., Miyashita, A., Williams, D., Ströhl, F., et al. (2018). FUS Phase Separation Is Modulated by a Molecular Chaperone and Methylation of Arginine Cation-π Interactions. Cell 173, 720–734.e15.

Ramer, G., Ruggeri, F.S., Levin, A., Knowles, T.P.J., and Centrone, A. (2018). Determination of Polypeptide Conformation with Nanoscale Resolution in Water. ACS Nano 12, 6612–6619.

Rouches, M., Veatch, S.L., and Machta, B.B. (2021). Surface densities prewet a near-critical membrane. Proc. Natl. Acad. Sci. U. S. A. 118.

Ruggeri, F.S., Adamcik, J., Jeong, J.S., Lashuel, H.A., Mezzenga, R., and Dietler, G. (2015a). Influence of the β-sheet content on the mechanical properties of aggregates during amyloid fibrillization. Angew. Chemie - Int. Ed. 54, 2462–2466.

Ruggeri, F.S., Longo, G., Faggiano, S., Lipiec, E., Pastore, A., and Dietler, G. (2015b). Infrared nanospectroscopy characterization of oligomeric and fibrillar aggregates during amyloid formation. Nat. Commun. 6, 1–9.

Ruggeri, F.S., Vieweg, S., Cendrowska, U., Longo, G., Chiki, A., Lashuel, H.A., and Dietler, G. (2016). Nanoscale studies link amyloid maturity with polyglutamine diseases onset. Sci. Rep. 6, 1–11.

Ruggeri, F.S., Marcott, C., Dinarelli, S., Longo, G., Girasole, M., Dietler, G., and Knowles, T.P.J. (2018). Identification of oxidative stress in red blood cells with nanoscale chemical resolution by infrared nanospectroscopy. Int. J. Mol. Sci. 19.

Ruggeri, F.S., Mannini, B., Schmid, R., Vendruscolo, M., and Knowles, T.P.J. (2020). Single molecule secondary structure determination of proteins through infrared absorption nanospectroscopy. Nat. Commun. 11, 1–9.

Ruggeri, F.S., Habchi, J., Chia, S., Horne, R.I., Vendruscolo, M., and Knowles, T.P.J. (2021). Infrared nanospectroscopy reveals the molecular interaction fingerprint of an aggregation inhibitor with single Aβ42 oligomers. Nat. Commun. 12, 1–9.

Satoh, H., Shibata, H., Nakano, Y., Kitaura, Y., and Maki, M. (2002). ALG-2 interacts with the amino-terminal domain of annexin XI in a Ca2+-dependent manner. Biochem. Biophys. Res. Commun. 291, 1166–1172.

Scheiffele, P., Rietveld, A., Wilk, T., and Simons, K. (1999). Influenza viruses select ordered lipid domains during budding from the plasma membrane. J. Biol. Chem. 274, 2038–2044.

Schneider, M.M., Gautam, S., Herling, T.W., Andrzejewska, E., Krainer, G., Miller, A.M., Trinkaus, V.A., Peter, Q.A.E., Ruggeri, F.S., Vendruscolo, M., et al. (2021). The Hsc70 disaggregation machinery removes monomer units directly from α-synuclein fibril ends. Nat. Commun. 12, 1–11.

Seo, A.Y., Sarkleti, F., Budin, I., Chang, C., King, C., Kohlwein, S.-D., Sengupta, P., and Lippincott-Schwartz, J. (2021). Vacuole phase-partitioning boosts mitochondria activity and cell lifespan through an inter-organelle lipid pipeline. BioRxiv 2021.04.11.439383.

Shaikh, S.R., Dumaual, A.C., Jenski, L.J., and Stillwell, W. (2001). Lipid phase separation in phospholipid bilayers and monolayers modeling the plasma membrane. Biochim. Biophys. Acta - Biomembr. 1512, 317–328.

Shen, Y., and Bax, A. (2013). Protein backbone and sidechain torsion angles predicted from NMR chemical shifts using artificial neural networks. J. Biomol. NMR 56, 227–241.

Shen, Y., Ruggeri, F.S., Vigolo, D., Kamada, A., Qamar, S., Levin, A., Iserman, C., Alberti, S., George-Hyslop, P.S., and Knowles, T.P.J. (2020). Biomolecular condensates undergo a generic shear-mediated liquid-to-solid transition. Nat. Nanotechnol. 15, 841–847.

Simone Ruggeri, F., Habchi, J., Cerreta, A., and Dietler, G. (2016). AFM-Based Single Molecule Techniques: Unraveling the Amyloid Pathogenic Species. Curr. Pharm. Des. 22, 3950–3970.

Simons, K., and Ikonen, E. (1997). Functional rafts in cell membranes. Nature 387, 569–572.

Speckner, K., Stadler, L., and Weiss, M. (2021). Unscrambling exit site patterns on the endoplasmic reticulum as a quenched demixing process. Biophys. J. 120, 2532–2542.

Su, X., Ditlev, J.A., Hui, E., Xing, W., Banjade, S., Okrut, J., King, D.S., Taunton, J., Rosen, M.K., and Vale, R.D. (2016). Phase separation of signaling molecules promotes T cell receptor signal transduction. Science (80-.). 352, 595–599.

Tamm, L.K., and Tatulian, S.A. (1997). Infrared spectroscopy of proteins and peptides in lipid bilayers. Q. Rev. Biophys. 30, 365–429.

Thelen, A.M., and Zoncu, R. (2017). Emerging Roles for the Lysosome in Lipid Metabolism. Trends Cell Biol. 27, 833–850.

Topp, S.D., Fallini, C., Shibata, H., Chen, H.J., Troakes, C., King, A., Ticozzi, N., Kenna, K.P., Soragia-Gkazi, A., Miller, J.W., et al. (2017). Mutations in the vesicular trafficking protein Annexin A11 are associated with amyotrophic lateral sclerosis Bradley N. Smith. Sci. Transl. Med. 9.

Valanciunaite, J., Kempf, E., Seki, H., Danylchuk, D.I., Peyriéras, N., Niko, Y., and Klymchenko, A.S. (2020). Polarity Mapping of Cells and Embryos by Improved Fluorescent Solvatochromic Pyrene Probe. Anal. Chem. 92, 6512–6520.

Velve-Casquillas, G., Costa, J., Carlier-Grynkorn, F., Mayeux, A., and Tran, P.T. (2010). A Fast Microfluidic Temperature Control Device for Studying Microtubule Dynamics in Fission Yeast (NIH Public Access).

Wang, H.-Y., Chan, S.H., Dey, S., Castello-Serrano, I., Ditlev, J.A., Rosen, M.K., Levental, K.R., and Levental, I. (2022). Coupling of protein condensates to ordered lipid domains determines functional membrane organization. BioRxiv 2022.08.02.502487.

Watanabe, M., Ando, Y., Tokumitsu, H., and Hidaka, H. (1993). Binding site of annexin XI on the calcyclin molecule. Biochem. Biophys. Res. Commun. 196, 1376–1382.

Weisz, J., and Uversky, V.N. (2020). Zooming into the dark side of human annexin-s100 complexes: Dynamic alliance of flexible partners. Int. J. Mol. Sci. 21, 1–30.

Xu, Y., Zhu, H., Shen, Y., Guttenplan, A.P.M., Saar, K.L., Lu, Y., Vigolo, D., Itzhaki, L.S., and Knowles, T.P.J. (2022). Micromechanics of soft materials using microfluidics. MRS Bull. 47, 119–126.

Yang, L., Zhang, G.J., Zhong, Y.G., and Zheng, Y.Z. (2000). Influence of membrane fluidity modifiers on lysosomal osmotic sensitivity. Cell Biol. Int. 24, 699–704.

Youn, J.Y., Dyakov, B.J.A., Zhang, J., Knight, J.D.R., Vernon, R.M., Forman-Kay, J.D., and Gingras, A.C. (2019). Properties of Stress Granule and P-Body Proteomes. Mol. Cell 76, 286– 294.

Zacharogianni, M., Gomez, A.A., Veenendaal, T., Smout, J., and Rabouille, C. (2014). A stress assembly that confers cell viability by preserving ERES components during amino-acid starvation. Elife 3, 1–25.

Zeng, M., Chen, X., Guan, D., Xu, J., Wu, H., Tong, P., and Zhang, M. (2018). Reconstituted Postsynaptic Density as a Molecular Platform for Understanding Synapse Formation and Plasticity. Cell 174, 1172–1187.e16.

Zhang, C., and Rabouille, C. (2019). Membrane-Bound Meet Membraneless in Health and Disease. Cells 8, 1000.

Zhang, Y., Herling, T.W., Kreida, S., Peter, Q.A.E., Kartanas, T., Törnroth-Horsefield, S., Linse, S., and Knowles, T.P.J. (2020). A microfluidic strategy for the detection of membrane protein interactions. Lab Chip 20, 3230–3238.

