## Supplementary material for "ANXA11 biomolecular condensates facilitate protein-lipid phase coupling on lysosomal membranes": Table S1

| Protein Group | Protein ID | Accession | -10lgP | Coverage (%) | #Peptides | #Unique | Avg. Mass | gene | Present on RNA granule database (Youn et al., 2019) |
|---|---|---|---|---|---|---|---|---|---|
| 2 | 2 | P68104\|EF1A1_HUMAN | 282.82 | 52 | 39 | 30 | 50141 | EEF1A1 | yes |
| 7 | 1 | Q14204\|DYHC1_HUMAN | 274.91 | 14 | 43 | 43 | 532412 | DYNC1H1 | yes |
| 3 | 11 | P07437\|TBB5_HUMAN | 265.17 | 51 | 30 | 4 | 49671 | TUBB | yes |
| 4 | 15 | Q9BVA1\|TBB2B_HUMAN | 258.77 | 44 | 28 | 0 | 49953 | TUBB2B | no |
| 5 | 20 | P68371\|TBB4B_HUMAN | 256.26 | 44 | 27 | 1 | 49831 | TUBB4B | no |
| 6 | 21 | Q13885\|TBB2A_HUMAN | 246.35 | 44 | 26 | 2 | 49907 | TUBB2A | yes |
| 9 | 7 | Q71U36\|TBA1A_HUMAN | 245.72 | 53 | 21 | 1 | 50136 | TUBA1A | yes |
| 10 | 9 | P68363\|TBA1B_HUMAN | 239.85 | 53 | 19 | 0 | 50152 | TUBA1B | yes |
| 14 | 4 | P42704\|LPPRC_HUMAN | 234.53 | 20 | 29 | 29 | 157904 | LRPPRC | yes |
| 11 | 14 | P68366\|TBA4A_HUMAN | 232.35 | 48 | 17 | 1 | 49924 | TUBA4A | no |
| 17 | 10 | Q9BQE3\|TBA1C_HUMAN | 225.62 | 45 | 17 | 0 | 49895 | TUBA1C | no |
| 12 | 17 | Q05639\|EF1A2_HUMAN | 224.43 | 42 | 15 | 6 | 50470 | EEF1A2 | yes |
| 13 | 6 | P08238\|HS90B_HUMAN | 223.26 | 31 | 17 | 11 | 83264 | HSP90AB1 | yes |
| 16 | 18 | Q13509\|TBB3_HUMAN | 219.83 | 42 | 18 | 7 | 50433 | TUBB3 | no |
| 16 | 19 | tr\|A0A0B4J269\|A0A0B4J269_HUMAN | 219.83 | 23 | 18 | 7 | 88382 | NA | no |
| 25 | 8 | P10809\|CH60_HUMAN | 219.46 | 37 | 17 | 17 | 61055 | HSPD1 | yes |
| 23 | 13 | P25705\|ATPA_HUMAN | 217.62 | 30 | 17 | 17 | 59751 | ATP5F1A | yes |
| 22 | 23 | P04406\|G3P_HUMAN | 213.14 | 42 | 16 | 16 | 36053 | GAPDH | yes |
| 26 | 22 | Q9BUF5\|TBB6_HUMAN | 211.48 | 33 | 20 | 5 | 49857 | TUBB6 | yes |
| 18 | 43 | O15240\|VGF_HUMAN | 208.31 | 36 | 19 | 19 | 67258 | VGF | no |
| 19 | 31 | P63261\|ACTG_HUMAN | 199.12 | 46 | 15 | 12 | 41793 | ACTG1 | no |
| 24 | 36 | P06576\|ATPB_HUMAN | 196.05 | 32 | 20 | 20 | 56560 | ATP5F1B | yes |
| 27 | 63 | Q9NY65\|TBA8_HUMAN | 194.64 | 23 | 9 | 0 | 50094 | TUBA8 | no |
| 20 | 24 | P07900\|HS90A_HUMAN | 193.13 | 23 | 12 | 6 | 84660 | HSP90AA1 | yes |
| 33 | 38 | Q9P2J5\|SYLC_HUMAN | 188.79 | 11 | 11 | 11 | 134466 | LARS | yes |
| 43 | 12 | P35580\|MYH10_HUMAN | 187.53 | 10 | 10 | 10 | 228997 | MYH10 | yes |
| 35 | 16 | P36776\|LONM_HUMAN | 186.12 | 22 | 12 | 12 | 106489 | LONP1 | yes |
| 60 | 265 | P30101\|PDIA3_HUMAN | 181.52 | 17 | 7 | 7 | 56782 | PDIA3 | yes |

| | | | | | | | | | |
|---|---|---|---|---|---|---|---|---|---|
| 50 | 66 | P14618\|KPYM_HUMAN | 179.82 | 17 | 7 | 7 | 57937 | PKM | yes |
| 31 | 40 | Q16658\|FSCN1_HUMAN | 177.51 | 28 | 12 | 12 | 54530 | FSCN1 | yes |
| 42 | 37 | P78527\|PRKDC_HUMAN | 174.45 | 4 | 10 | 10 | 469093 | PRKDC | yes |
| 29 | 45 | P12277\|KCRB_HUMAN | 172.25 | 27 | 11 | 11 | 42644 | CKB | yes |
| 37 | 33 | Q7KZF4\|SND1_HUMAN | 168.84 | 15 | 9 | 9 | 101997 | SND1 | yes |
| 44 | 116 | P17980\|PRS6A_HUMAN | 167.84 | 26 | 8 | 8 | 49204 | PSMC3 | yes |
| 40 | 58 | P27824\|CALX_HUMAN | 167.09 | 23 | 8 | 8 | 67568 | CANX | yes |
| 32 | 52 | P13639\|EF2_HUMAN | 166.99 | 14 | 10 | 10 | 95338 | EEF2 | yes |
| 36 | 32 | P60842\|IF4A1_HUMAN | 166.67 | 26 | 9 | 4 | 46154 | EIF4A1 | yes |
| 30 | 29 | Q92598\|HS105_HUMAN | 165.8 | 16 | 9 | 9 | 96865 | HSPH1 | yes |
| 72 | 170 | P06733\|ENOA_HUMAN | 163.59 | 25 | 6 | 5 | 47169 | ENO1 | yes |
| 41 | 65 | Q14195\|DPYL3_HUMAN | 163.22 | 26 | 8 | 5 | 61963 | DPYSL3 | yes |
| 34 | 47 | P08670\|VIME_HUMAN | 163.14 | 20 | 10 | 10 | 53652 | VIM | yes |
| 28 | 83 | P39023\|RL3_HUMAN | 162.36 | 18 | 7 | 7 | 46109 | RPL3 | yes |
| 78 | 97 | P78371\|TCPB_HUMAN | 162.18 | 17 | 7 | 7 | 57488 | CCT2 | yes |
| 80 | 35 | P41252\|SYIC_HUMAN | 161.83 | 9 | 8 | 8 | 144498 | IARS | yes |
| 52 | 386 | P63241\|IF5A1_HUMAN | 160.27 | 37 | 6 | 6 | 16832 | EIF5A | yes |
| 89 | 123 | P48643\|TCPE_HUMAN | 159.83 | 22 | 6 | 6 | 59671 | CCT5 | yes |
| 46 | 28 | P55072\|TERA_HUMAN | 158.79 | 14 | 8 | 8 | 89322 | VCP | yes |
| 51 | 121 | P24534\|EF1B_HUMAN | 157.14 | 37 | 9 | 9 | 24764 | EEF1B2 | no |
| 64 | 79 | Q99832\|TCPH_HUMAN | 156.48 | 17 | 8 | 8 | 59367 | CCT7 | yes |
| 39 | 44 | P63244\|RACK1_HUMAN | 155.77 | 28 | 6 | 6 | 35077 | RACK1 | yes |
| 75 | 51 | P52597\|HNRPF_HUMAN | 155.5 | 21 | 7 | 5 | 45672 | HNRNPF | yes |
| 66 | 73 | P13645\|K1C10_HUMAN | 152.2 | 19 | 7 | 6 | 58827 | KRT10 | no |
| 81 | 61 | Q13813\|SPTN1_HUMAN | 153.68 | 5 | 7 | 7 | 284538 | SPTAN1 | yes |
| 82 | 39 | Q6P2E9\|EDC4_HUMAN | 153.47 | 10 | 7 | 7 | 151661 | EDC4 | yes |
| 68 | 72 | Q14240\|IF4A2_HUMAN | 153.22 | 21 | 6 | 1 | 46402 | EIF4A2 | no |
| 74 | 46 | P45974\|UBP5_HUMAN | 153.12 | 11 | 7 | 7 | 95786 | USP5 | yes |
| 62 | 77 | Q00610\|CLH1_HUMAN | 152.71 | 7 | 7 | 7 | 191613 | CLTC | yes |
| 90 | 98 | P16615\|AT2A2_HUMAN | 151.39 | 7 | 6 | 5 | 114757 | ATP2A2 | yes |
| 56 | 74 | P31939\|PUR9_HUMAN | 150.57 | 24 | 8 | 8 | 64616 | ATIC | yes |
| 55 | 59 | Q14152\|EIF3A_HUMAN | 150 | 8 | 7 | 7 | 166569 | EIF3A | yes |
| 86 | 53 | P08865\|RSSA_HUMAN | 149.12 | 20 | 5 | 5 | 32854 | RPSA | yes |
| 65 | 153 | P46821\|MAP1B_HUMAN | 148.11 | 6 | 8 | 8 | 270632 | MAP1B | yes |

| | | | | | | | | | |
|---|---|---|---|---|---|---|---|---|---|
| 45 | 69 | P0DMV9\|HS71B_HUMAN | 146.87 | 24 | 8 | 7 | 70052 | HSPA1B | no |
| 48 | 114 | P35579\|MYH9_HUMAN | 146.5 | 7 | 8 | 8 | 226530 | MYH9 | yes |
| 85 | 85 | P55084\|ECHB_HUMAN | 146.11 | 21 | 5 | 5 | 51294 | HADHB | yes |
| 103 | 62 | P49368\|TCPG_HUMAN | 145.73 | 18 | 5 | 5 | 60534 | CCT3 | yes |
| 79 | 142 | P22314\|UBA1_HUMAN | 145.65 | 13 | 7 | 7 | 117849 | UBA1 | yes |
| 111 | 102 | P05388\|RLA0_HUMAN | 144.22 | 18 | 5 | 5 | 34274 | RPLP0 | yes |
| 67 | 50 | P19338\|NUCL_HUMAN | 143.73 | 11 | 5 | 5 | 76615 | NCL | yes |
| 122 | 272 | O60506\|HNRPQ_HUMAN | 143.69 | 13 | 5 | 5 | 69603 | SYNCRIP | yes |
| 47 | 42 | P04264\|K2C1_HUMAN | 143.14 | 15 | 10 | 10 | 66039 | KRT1 | no |
| 70 | 67 | P31943\|HNRH1_HUMAN | 142.42 | 20 | 6 | 4 | 49229 | HNRNPH1 | yes |
| 189 | 154 | P51148\|RAB5C_HUMAN | 141.06 | 28 | 4 | 4 | 23483 | RAB5C | yes |
| 53 | 279 | P04792\|HSPB1_HUMAN | 140.43 | 34 | 7 | 7 | 22783 | HSPB1 | yes |
| 125 | 55 | P07814\|SYEP_HUMAN | 139.6 | 4 | 4 | 4 | 170590 | EPRS | yes |
| 107 | 138 | P17987\|TCPA_HUMAN | 138.9 | 11 | 6 | 6 | 60344 | TCP1 | yes |
| 76 | 56 | Q14194\|DPYL1_HUMAN | 138.56 | 24 | 6 | 5 | 62184 | CRMP1 | no |
| 104 | 103 | P04844\|RPN2_HUMAN | 138.31 | 16 | 5 | 5 | 69284 | RPN2 | yes |
| 83 | 49 | P40939\|ECHA_HUMAN | 138.24 | 12 | 6 | 6 | 83000 | HADHA | yes |
| 54 | 48 | P49327\|FAS_HUMAN | 138.16 | 5 | 7 | 7 | 273424 | FASN | yes |
| 73 | 151 | P62937\|PPIA_HUMAN | 138.04 | 19 | 6 | 6 | 18012 | PPIA | yes |
| 15 | 234 | P07477\|TRY1_HUMAN | 137.85 | 19 | 13 | 9 | 26558 | PRSS1 | no |
| 108 | 57 | P08758\|ANXA5_HUMAN | 135.82 | 30 | 5 | 5 | 35937 | ANXA5 | yes |
| 119 | 300 | P50454\|SERPH_HUMAN | 135.43 | 19 | 4 | 4 | 46441 | SERPINH1 | yes |
| 87 | 92 | P40227\|TCPZ_HUMAN | 135.35 | 15 | 5 | 5 | 58024 | CCT6A | yes |
| 96 | 124 | P27797\|CALR_HUMAN | 135.25 | 16 | 6 | 6 | 48142 | CALR | yes |
| 77 | 89 | P04843\|RPN1_HUMAN | 134.69 | 15 | 6 | 6 | 68569 | RPN1 | yes |
| 112 | 115 | P35232\|PHB_HUMAN | 134.61 | 25 | 5 | 5 | 29804 | PHB | yes |
| 88 | 86 | P49411\|EFTU_HUMAN | 134.51 | 19 | 6 | 6 | 49542 | TUFM | yes |
| 173 | 246 | Q9NQC3\|RTN4_HUMAN | 134.07 | 3 | 3 | 3 | 129931 | RTN4 | yes |
| 49 | 277 | Q15019\|SEPT2_HUMAN | 132.46 | 19 | 5 | 5 | 41487 | Sep-02 | no |
| 63 | 135 | P35527\|K1C9_HUMAN | 132.25 | 19 | 10 | 10 | 62064 | KRT9 | no |
| 146 | 118 | P62191\|PRS4_HUMAN | 131.56 | 15 | 5 | 5 | 49185 | PSMC1 | yes |
| 129 | 220 | P12956\|XRCC6_HUMAN | 130.96 | 9 | 4 | 4 | 69843 | XRCC6 | yes |
| 21 | 709 | P07478\|TRY2_HUMAN | 130.91 | 21 | 10 | 6 | 26488 | PRSS2 | no |
| 57 | 137 | Q16555\|DPYL2_HUMAN | 130.68 | 17 | 6 | 4 | 62294 | DPYSL2 | yes |

| | | | | | | | | | |
|---|---|---|---|---|---|---|---|---|---|
| 61 | 449 | P63220\|RS21_HUMAN | 130.42 | 40 | 3 | 3 | 9111 | RPS21 | yes |
| 59 | 90 | P62701\|RS4X_HUMAN | 130.31 | 20 | 6 | 6 | 29598 | RPS4X | yes |
| 71 | 54 | P61978\|HNRPK_HUMAN | 129.94 | 27 | 6 | 6 | 50976 | HNRNPK | yes |
| 139 | 240 | Q9Y4L1\|HYOU1_HUMAN | 129.42 | 7 | 4 | 4 | 111335 | HYOU1 | yes |
| 58 | 229 | P62081\|RS7_HUMAN | 129.39 | 34 | 5 | 5 | 22127 | RPS7 | yes |
| 113 | 148 | Q9UHD8\|SEPT9_HUMAN | 128.34 | 10 | 3 | 3 | 65402 | Sep-09 | no |
| 136 | 119 | P30049\|ATPD_HUMAN | 128.32 | 48 | 5 | 5 | 17490 | ATP5F1D | no |
| 143 | 112 | P62495\|ERF1_HUMAN | 127.05 | 16 | 4 | 4 | 49031 | ETF1 | yes |
| 91 | 107 | P32969\|RL9_HUMAN | 127.03 | 35 | 5 | 5 | 21863 | RPL9 | yes |
| 102 | 78 | O95202\|LETM1_HUMAN | 126.72 | 11 | 4 | 4 | 83354 | LETM1 | no |
| 84 | 82 | P54289\|CA2D1_HUMAN | 126.53 | 8 | 4 | 4 | 124568 | CACNA2D1 | no |
| 93 | 139 | Q16181\|SEPT7_HUMAN | 125.54 | 16 | 5 | 5 | 50680 | Sep-07 | no |
| 123 | 310 | Q07960\|RHG01_HUMAN | 124.81 | 18 | 5 | 5 | 50436 | ARHGAP1 | no |
| 126 | 93 | Q13423\|NNTM_HUMAN | 124.69 | 7 | 4 | 4 | 113895 | NNT | yes |
| 141 | 104 | Q15084\|PDIA6_HUMAN | 124.01 | 17 | 4 | 4 | 48121 | PDIA6 | yes |
| 121 | 308 | Q92499\|DDX1_HUMAN | 123.82 | 8 | 5 | 5 | 82432 | DDX1 | yes |
| 115 | 195 | P12532\|KCRU_HUMAN | 123.44 | 17 | 3 | 3 | 47037 | CKMT1A | no |
| 147 | 120 | O43390\|HNRPR_HUMAN | 123.14 | 9 | 4 | 4 | 70943 | HNRNPR | yes |
| 99 | 80 | A6NKG5\|RTL1_HUMAN | 122.81 | 6 | 5 | 5 | 155047 | RTL1 | no |
| 144 | 113 | O60568\|PLOD3_HUMAN | 122.75 | 9 | 4 | 4 | 84785 | PLOD3 | yes |
| 170 | 146 | O75534\|CSDE1_HUMAN | 120.97 | 10 | 5 | 5 | 88885 | CSDE1 | yes |
| 133 | 127 | P38646\|GRP75_HUMAN | 120.56 | 11 | 4 | 4 | 73681 | HSPA9 | yes |
| 186 | 117 | Q9BSJ8\|ESYT1_HUMAN | 119.22 | 6 | 3 | 3 | 122856 | ESYT1 | yes |
| 155 | 268 | P06748\|NPM_HUMAN | 119.09 | 22 | 3 | 3 | 32575 | NPM1 | yes |
| 154 | 132 | Q9BPW8\|NIPS1_HUMAN | 118.63 | 21 | 4 | 4 | 33310 | NIPSNAP1 | no |
| 224 | 450 | P27635\|RL10_HUMAN | 118.51 | 13 | 3 | 3 | 24604 | RPL10 | no |
| 130 | 203 | Q9H857\|NT5D2_HUMAN | 117.23 | 8 | 4 | 4 | 60719 | NT5DC2 | no |
| 150 | 419 | Q10471\|GALT2_HUMAN | 117.18 | 11 | 4 | 4 | 64733 | GALNT2 | yes |
| 208 | 270 | P00387\|NB5R3_HUMAN | 116.83 | 16 | 3 | 3 | 34235 | CYB5R3 | yes |
| 295 | 343 | O00410\|IPO5_HUMAN | 116.79 | 5 | 3 | 3 | 123630 | IPO5 | yes |
| 187 | 129 | O60716\|CTND1_HUMAN | 116.44 | 6 | 3 | 3 | 108170 | CTNND1 | yes |
| 239 | 145 | P56730\|NETR_HUMAN | 116.2 | 7 | 3 | 3 | 97067 | PRSS12 | no |
| 120 | 109 | P13637\|AT1A3_HUMAN | 116.09 | 8 | 4 | 2 | 111748 | ATP1A3 | no |
| 120 | 108 | tr\|A0A2R8YEY8\|A0A2R8YEY8_HUMAN | 116.09 | 9 | 4 | 2 | 111196 | NA | no |

| | | | | | | | | | |
|---|---|---|---|---|---|---|---|---|---|
| 195 | 202 | Q8IXB1\|DJC10_HUMAN | 116.05 | 5 | 3 | 3 | 91080 | DNAJC10 | no |
| 193 | 185 | P22102\|PUR2_HUMAN | 115.56 | 6 | 3 | 3 | 107767 | GART | yes |
| 98 | 149 | P21333\|FLNA_HUMAN | 114.7 | 5 | 6 | 6 | 280737 | FLNA | yes |
| 106 | 100 | O43795\|MYO1B_HUMAN | 114.45 | 6 | 4 | 4 | 131985 | MYO1B | yes |
| 142 | 237 | Q14157\|UBP2L_HUMAN | 114.25 | 7 | 3 | 3 | 114534 | UBAP2L | yes |
| 321 | 264 | P09211\|GSTP1_HUMAN | 113.7 | 21 | 2 | 2 | 23356 | GSTP1 | yes |
| 151 | 352 | P28331\|NDUS1_HUMAN | 113.6 | 8 | 3 | 3 | 79468 | NDUFS1 | yes |
| 210 | 276 | P61764\|STXB1_HUMAN | 112.93 | 10 | 3 | 3 | 67569 | STXBP1 | yes |
| 110 | 91 | P53621\|COPA_HUMAN | 112.9 | 5 | 3 | 3 | 138345 | COPA | yes |
| 201 | 537 | Q16891\|MIC60_HUMAN | 112.57 | 6 | 4 | 4 | 83678 | IMMT | yes |
| 347 | 451 | Q9Y512\|SAM50_HUMAN | 112.4 | 6 | 2 | 2 | 51976 | SAMM50 | no |
| 135 | 171 | Q12931\|TRAP1_HUMAN | 112.2 | 10 | 4 | 4 | 80110 | TRAP1 | yes |
| 101 | 179 | P50991\|TCPD_HUMAN | 111.6 | 11 | 3 | 3 | 57924 | CCT4 | yes |
| 138 | 164 | O60282\|KIF5C_HUMAN | 111.2 | 5 | 3 | 2 | 109495 | KIF5C | yes |
| 128 | 99 | Q86UP2\|KTN1_HUMAN | 111.1 | 4 | 4 | 4 | 156275 | KTN1 | yes |
| 131 | 309 | Q99733\|NP1L4_HUMAN | 110.29 | 9 | 3 | 3 | 42823 | NAP1L4 | yes |
| 198 | 284 | P17844\|DDX5_HUMAN | 109.91 | 8 | 3 | 3 | 69148 | DDX5 | yes |
| 206 | 163 | P33991\|MCM4_HUMAN | 108.71 | 5 | 3 | 3 | 96558 | MCM4 | yes |
| 149 | 177 | O43143\|DHX15_HUMAN | 108.7 | 7 | 3 | 3 | 90933 | DHX15 | yes |
| 127 | 60 | P27708\|PYR1_HUMAN | 108.56 | 5 | 4 | 4 | 242981 | CAD | yes |
| 172 | 187 | O00429\|DNM1L_HUMAN | 107.61 | 9 | 3 | 3 | 81877 | DNM1L | yes |
| 118 | 87 | P55795\|HNRH2_HUMAN | 106.8 | 16 | 4 | 3 | 49264 | HNRNPH2 | yes |
| 134 | 75 | P00367\|DHE3_HUMAN | 106.59 | 9 | 4 | 4 | 61398 | GLUD1 | yes |
| 134 | 81 | P49448\|DHE4_HUMAN | 106.59 | 9 | 4 | 4 | 61434 | GLUD2 | no |
| 175 | 207 | O60763\|USO1_HUMAN | 106.47 | 6 | 3 | 3 | 107895 | USO1 | no |
| 181 | 430 | Q92616\|GCN1_HUMAN | 105.94 | 2 | 4 | 4 | 292756 | GCN1 | yes |
| 162 | 208 | P05023\|AT1A1_HUMAN | 105.54 | 6 | 3 | 1 | 112896 | ATP1A1 | yes |
| 153 | 618 | P00403\|COX2_HUMAN | 105.07 | 16 | 4 | 4 | 25565 | MT | no |
| 207 | 248 | Q14444\|CAPR1_HUMAN | 104.94 | 12 | 3 | 3 | 78366 | CAPRIN1 | yes |
| 191 | 181 | Q9Y285\|SYFA_HUMAN | 104.84 | 9 | 2 | 2 | 57564 | FARSA | yes |
| 109 | 76 | P13010\|XRCC5_HUMAN | 104.74 | 12 | 4 | 4 | 82705 | XRCC5 | yes |
| 95 | 125 | P68032\|ACTC_HUMAN | 104.74 | 15 | 4 | 1 | 42019 | ACTC1 | yes |
| 95 | 126 | P68133\|ACTS_HUMAN | 104.74 | 15 | 4 | 1 | 42051 | ACTA1 | no |
| 148 | 288 | Q9Y230\|RUVB2_HUMAN | 104.66 | 14 | 4 | 4 | 51157 | RUVBL2 | yes |

| | | | | | | | | | |
|---|---|---|---|---|---|---|---|---|---|
| 168 | 150 | P08133\|ANXA6_HUMAN | 104.56 | 7 | 3 | 3 | 75873 | ANXA6 | yes |
| 299 | 363 | P13667\|PDIA4_HUMAN | 104.32 | 4 | 2 | 2 | 72933 | PDIA4 | yes |
| 100 | 84 | Q04637\|IF4G1_HUMAN | 104.26 | 5 | 5 | 5 | 175490 | EIF4G1 | yes |
| 236 | 622 | Q00839\|HNRPU_HUMAN | 104.05 | 6 | 3 | 3 | 90585 | HNRNPU | yes |
| 158 | 131 | P61163\|ACTZ_HUMAN | 103.81 | 14 | 3 | 3 | 42614 | ACTR1A | yes |
| 156 | 128 | O75947\|ATP5H_HUMAN | 102.8 | 32 | 3 | 3 | 18491 | ATP5PD | no |
| 244 | 158 | Q7L2E3\|DHX30_HUMAN | 102.38 | 4 | 3 | 3 | 133938 | DHX30 | yes |
| 137 | 155 | P09651\|ROA1_HUMAN | 102.32 | 15 | 4 | 4 | 38747 | HNRNPA1 | no |
| 137 | 169 | Q32P51\|RA1L2_HUMAN | 102.32 | 17 | 4 | 4 | 34225 | HNRNPA1L2 | yes |
| 183 | 105 | P55209\|NP1L1_HUMAN | 102.01 | 15 | 3 | 3 | 45374 | NAP1L1 | yes |
| 213 | 324 | Q16623\|STX1A_HUMAN | 101.78 | 15 | 3 | 3 | 33023 | STX1A | no |
| 322 | 266 | Q92900\|RENT1_HUMAN | 101.71 | 3 | 2 | 2 | 124345 | UPF1 | yes |
| 277 | 387 | P23588\|IF4B_HUMAN | 101.7 | 8 | 2 | 2 | 69151 | EIF4B | yes |
| 180 | 252 | P08754\|GNAI3_HUMAN | 101.65 | 9 | 3 | 2 | 40532 | GNAI3 | yes |
| 190 | 180 | O43776\|SYNC_HUMAN | 100.92 | 6 | 3 | 3 | 62943 | NARS | yes |
| 246 | 217 | P47897\|SYQ_HUMAN | 100.34 | 5 | 2 | 2 | 87799 | QARS | yes |
| 351 | 472 | Q6PI48\|SYDM_HUMAN | 100.12 | 6 | 2 | 2 | 73563 | DARS2 | yes |
| 323 | 267 | P20290\|BTF3_HUMAN | 100.01 | 24 | 2 | 2 | 22168 | BTF3 | yes |
| 215 | 286 | P62258\|1433E_HUMAN | 99.85 | 13 | 3 | 3 | 29174 | YWHAE | yes |
| 94 | 269 | O00571\|DDX3X_HUMAN | 99.76 | 6 | 2 | 2 | 73244 | DDX3X | yes |
| 184 | 209 | Q92973\|TNPO1_HUMAN | 99.7 | 6 | 3 | 3 | 102355 | TNPO1 | yes |
| 69 | 111 | P15880\|RS2_HUMAN | 99.29 | 12 | 3 | 3 | 31324 | RPS2 | yes |
| 253 | 190 | Q9HCM4\|E41L5_HUMAN | 99.16 | 4 | 3 | 3 | 81856 | EPB41L5 | no |
| 273 | 165 | Q13435\|SF3B2_HUMAN | 99.02 | 6 | 3 | 3 | 100228 | SF3B2 | no |
| 232 | 420 | P30044\|PRDX5_HUMAN | 98.98 | 8 | 3 | 3 | 22086 | PRDX5 | yes |
| 182 | 94 | P14625\|ENPL_HUMAN | 98.51 | 5 | 3 | 3 | 92469 | HSP90B1 | yes |
| 352 | 475 | O43169\|CYB5B_HUMAN | 98.28 | 23 | 2 | 2 | 16332 | CYB5B | no |
| 314 | 1103 | Q53H12\|AGK_HUMAN | 98.25 | 13 | 3 | 3 | 47137 | AGK | no |
| 105 | 95 | P35908\|K22E_HUMAN | 98.11 | 10 | 5 | 5 | 65433 | KRT2 | yes |
| 216 | 236 | P04181\|OAT_HUMAN | 98.1 | 10 | 3 | 3 | 48535 | OAT | no |
| 165 | 676 | P62244\|RS15A_HUMAN | 97.95 | 26 | 3 | 3 | 14840 | RPS15A | yes |
| 185 | 110 | P30837\|AL1B1_HUMAN | 97.71 | 11 | 3 | 3 | 57206 | ALDH1B1 | yes |
| 331 | 456 | P46782\|RS5_HUMAN | 97.3 | 17 | 2 | 2 | 22876 | RPS5 | yes |
| 192 | 182 | P52815\|RM12_HUMAN | 97.17 | 24 | 3 | 3 | 21348 | MRPL12 | no |

| | | | | | | | | | |
|---|---|---|---|---|---|---|---|---|---|
| 192 | 183 | tr\|B4DLN1\|B4DLN1_HUMAN | 97.17 | 11 | 3 | 3 | 48099 | NA | no |
| 209 | 271 | Q99623\|PHB2_HUMAN | 97.09 | 11 | 2 | 2 | 33296 | PHB2 | yes |
| 218 | 617 | P61011\|SRP54_HUMAN | 96.88 | 12 | 3 | 3 | 55705 | SRP54 | yes |
| 290 | 250 | P67775\|PP2AA_HUMAN | 96.79 | 12 | 2 | 1 | 35594 | PPP2CA | no |
| 199 | 316 | P34897\|GLYM_HUMAN | 96.73 | 10 | 3 | 3 | 55993 | SHMT2 | yes |
| 285 | 221 | Q13310\|PABP4_HUMAN | 96.71 | 6 | 2 | 2 | 70783 | PABPC4 | yes |
| 227 | 466 | Q14103\|HNRPD_HUMAN | 96.64 | 6 | 2 | 2 | 38434 | HNRNPD | yes |
| 205 | 287 | Q7Z2W9\|RM21_HUMAN | 96.5 | 19 | 3 | 3 | 22815 | MRPL21 | no |
| 230 | 496 | O00487\|PSDE_HUMAN | 95.98 | 11 | 3 | 3 | 34577 | PSMD14 | yes |
| 203 | 101 | P48147\|PPCE_HUMAN | 95.88 | 6 | 3 | 3 | 80700 | PREP | no |
| 166 | 831 | O94826\|TOM70_HUMAN | 95.86 | 7 | 3 | 3 | 67455 | TOMM70 | yes |
| 262 | 173 | P50990\|TCPQ_HUMAN | 95.63 | 6 | 3 | 3 | 59621 | CCT8 | yes |
| 160 | 273 | P33176\|KINH_HUMAN | 95.34 | 3 | 2 | 1 | 109685 | KIF5B | yes |
| 179 | 175 | Q7Z4S6\|KI21A_HUMAN | 95.32 | 3 | 2 | 2 | 187178 | KIF21A | no |
| 202 | 122 | Q9Y3I0\|RTCB_HUMAN | 94.86 | 10 | 3 | 3 | 55210 | RTCB | yes |
| 298 | 360 | Q14257\|RCN2_HUMAN | 94.79 | 12 | 2 | 2 | 36876 | RCN2 | yes |
| 212 | 280 | Q14011\|CIRBP_HUMAN | 94.54 | 19 | 2 | 2 | 18648 | CIRBP | yes |
| 350 | 469 | Q13084\|RM28_HUMAN | 94.46 | 18 | 2 | 2 | 30157 | MRPL28 | yes |
| 226 | 459 | Q04917\|1433F_HUMAN | 93.8 | 14 | 2 | 2 | 28219 | YWHAH | yes |
| 197 | 251 | P04899\|GNAI2_HUMAN | 93.69 | 8 | 3 | 2 | 40451 | GNAI2 | yes |
| 145 | 106 | Q9NSE4\|SYIM_HUMAN | 93.03 | 5 | 3 | 3 | 113791 | IARS2 | yes |
| 358 | 495 | P62820\|RAB1A_HUMAN | 92.78 | 19 | 2 | 2 | 22678 | RAB1A | yes |
| 324 | 274 | P34932\|HSP74_HUMAN | 92.45 | 4 | 2 | 2 | 94331 | HSPA4 | yes |
| 161 | 554 | P05141\|ADT2_HUMAN | 92.45 | 21 | 3 | 2 | 32852 | SLC25A5 | no |
| 157 | 130 | Q9Y265\|RUVB1_HUMAN | 92.18 | 10 | 4 | 4 | 50228 | RUVBL1 | yes |
| 315 | 1277 | Q14974\|IMB1_HUMAN | 92.11 | 3 | 2 | 2 | 97170 | KPNB1 | yes |
| 301 | 398 | Q8NC51\|PAIRB_HUMAN | 92.03 | 4 | 2 | 2 | 44965 | SERBP1 | yes |
| 221 | 336 | Q9NR31\|SAR1A_HUMAN | 91.82 | 17 | 3 | 3 | 22367 | SAR1A | no |
| 353 | 482 | P61020\|RAB5B_HUMAN | 91.75 | 18 | 2 | 2 | 23707 | RAB5B | no |
| 97 | 303 | P23396\|RS3_HUMAN | 91.29 | 14 | 3 | 3 | 26688 | RPS3 | yes |
| 333 | 474 | P57737\|CORO7_HUMAN | 91.28 | 8 | 2 | 2 | 100605 | CORO7 | yes |
| 296 | 350 | tr\|H3BNC9\|H3BNC9_HUMAN | 91.28 | 8 | 2 | 2 | 64533 | NA | no |
| 296 | 546 | P08708\|RS17_HUMAN | 91.28 | 33 | 2 | 2 | 15550 | RPS17 | no |
| 328 | 282 | P63096\|GNAI1_HUMAN | 91.19 | 8 | 2 | 1 | 40361 | GNAI1 | no |

| | | | | | | | | | |
|---|---|---|---|---|---|---|---|---|---|
| 114 | 174 | Q7Z7H5\|TMED4_HUMAN | 90.89 | 16 | 2 | 2 | 25943 | TMED4 | no |
| 282 | 200 | P43686\|PRS6B_HUMAN | 90.81 | 9 | 2 | 2 | 47366 | PSMC4 | yes |
| 320 | 291 | P24752\|THIL_HUMAN | 90.66 | 7 | 2 | 2 | 45200 | ACAT1 | yes |
| 167 | 162 | Q08211\|DHX9_HUMAN | 90.48 | 3 | 2 | 2 | 140958 | DHX9 | yes |
| 169 | 156 | P11586\|C1TC_HUMAN | 90.37 | 8 | 3 | 3 | 101559 | MTHFD1 | yes |
| 272 | 806 | P62140\|PP1B_HUMAN | 90.27 | 16 | 3 | 3 | 37187 | PPP1CB | no |
| 272 | 1034 | P36873\|PP1G_HUMAN | 84.2 | 11 | 2 | 2 | 36984 | PPP1CC | no |
| 272 | 1035 | P62136\|PP1A_HUMAN | 84.2 | 11 | 2 | 2 | 37512 | PPP1CA | yes |
| 326 | 278 | O15069\|NACAD_HUMAN | 90.21 | 3 | 2 | 2 | 161100 | NACAD | no |
| 349 | 463 | Q9NVH0\|EXD2_HUMAN | 90.15 | 6 | 2 | 2 | 70353 | EXD2 | no |
| 217 | 294 | Q9Y262\|EIF3L_HUMAN | 90.09 | 7 | 3 | 3 | 66727 | EIF3L | yes |
| 204 | 133 | P28838\|AMPL_HUMAN | 90.03 | 7 | 3 | 3 | 56166 | LAP3 | yes |
| 325 | 275 | Q9H269\|VPS16_HUMAN | 89.88 | 4 | 2 | 2 | 94694 | VPS16 | yes |
| 453 | 446 | P78344\|IF4G2_HUMAN | 89.61 | 5 | 2 | 2 | 102362 | EIF4G2 | yes |
| 366 | 530 | Q12905\|ILF2_HUMAN | 89.35 | 14 | 2 | 2 | 43062 | ILF2 | no |
| 223 | 337 | P05386\|RLA1_HUMAN | 89.34 | 21 | 3 | 3 | 11514 | RPLP1 | yes |
| 276 | 184 | P27694\|RFA1_HUMAN | 89.31 | 5 | 2 | 2 | 68138 | RPA1 | yes |
| 163 | 332 | P20674\|COX5A_HUMAN | 89.21 | 27 | 3 | 3 | 16762 | COX5A | no |
| 222 | 452 | O14983\|AT2A1_HUMAN | 89 | 5 | 2 | 1 | 110252 | ATP2A1 | yes |
| 318 | 2698 | P33993\|MCM7_HUMAN | 88.7 | 5 | 2 | 2 | 81308 | MCM7 | yes |
| 265 | 326 | Q9Y4W6\|AFG32_HUMAN | 88.5 | 7 | 2 | 2 | 88584 | AFG3L2 | no |
| 329 | 507 | P37802\|TAGL2_HUMAN | 88.32 | 14 | 2 | 2 | 22391 | TAGLN2 | yes |
| 274 | 176 | P05091\|ALDH2_HUMAN | 88.31 | 10 | 2 | 2 | 56381 | ALDH2 | yes |
| 565 | 445 | Q9BPX5\|ARP5L_HUMAN | 88.22 | 25 | 1 | 1 | 16941 | ARPC5L | no |
| 292 | 289 | Q9UGP8\|SEC63_HUMAN | 88.19 | 6 | 2 | 2 | 87997 | SEC63 | no |
| 116 | 406 | Q562R1\|ACTBL_HUMAN | 88.04 | 12 | 4 | 2 | 42003 | ACTBL2 | no |
| 361 | 511 | P49755\|TMEDA_HUMAN | 87.78 | 8 | 2 | 2 | 24976 | TMED10 | yes |
| 279 | 194 | P26599\|PTBP1_HUMAN | 87.74 | 9 | 2 | 2 | 57221 | PTBP1 | yes |
| 319 | 224 | Q6DD88\|ATLA3_HUMAN | 87.55 | 9 | 2 | 2 | 60542 | ATL3 | yes |
| 177 | 147 | Q16643\|DREB_HUMAN | 87.51 | 6 | 3 | 3 | 71429 | DBN1 | yes |
| 178 | 160 | Q00341\|VIGLN_HUMAN | 87.29 | 4 | 4 | 4 | 141455 | HDLBP | yes |
| 356 | 489 | Q9NVI7\|ATD3A_HUMAN | 87.27 | 3 | 2 | 2 | 71369 | ATAD3A | yes |
| 248 | 318 | P08183\|MDR1_HUMAN | 87.22 | 5 | 3 | 3 | 141478 | ABCB1 | yes |
| 255 | 213 | P54136\|SYRC_HUMAN | 87.02 | 7 | 4 | 4 | 75379 | RARS | yes |

| | | | | | | | | | |
|---|---|---|---|---|---|---|---|---|---|
| 211 | 222 | Q8TCS8\|PNPT1_HUMAN | 86.62 | 4 | 2 | 2 | 85951 | PNPT1 | yes |
| 287 | 227 | O60841\|IF2P_HUMAN | 86.54 | 3 | 2 | 2 | 138827 | EIF5B | yes |
| 308 | 512 | Q9UNF1\|MAGD2_HUMAN | 86.45 | 3 | 2 | 2 | 64954 | MAGED2 | yes |
| 247 | 161 | P31942\|HNRH3_HUMAN | 86.31 | 13 | 2 | 2 | 36926 | HNRNPH3 | yes |
| 355 | 488 | Q7L0Y3\|TM10C_HUMAN | 86.26 | 4 | 2 | 2 | 47347 | TRMT10C | yes |
| 242 | 193 | Q8WVM8\|SCFD1_HUMAN | 86.15 | 8 | 4 | 4 | 72380 | SCFD1 | yes |
| 231 | 510 | P62277\|RS13_HUMAN | 85.95 | 11 | 2 | 2 | 17222 | RPS13 | yes |
| 303 | 409 | Q9BYD3\|RM04_HUMAN | 85.93 | 8 | 2 | 2 | 34919 | MRPL4 | yes |
| 283 | 201 | P30153\|2AAA_HUMAN | 85.8 | 6 | 2 | 1 | 65309 | PPP2R1A | yes |
| 174 | 172 | Q9Y490\|TLN1_HUMAN | 85.68 | 3 | 3 | 3 | 269765 | TLN1 | yes |
| 269 | 322 | P53396\|ACLY_HUMAN | 85.55 | 4 | 3 | 3 | 120839 | ACLY | yes |
| 194 | 369 | O75489\|NDUS3_HUMAN | 85.29 | 12 | 3 | 3 | 30242 | NDUFS3 | no |
| 176 | 157 | Q9UJS0\|CMC2_HUMAN | 84.8 | 10 | 4 | 3 | 74176 | SLC25A13 | yes |
| 341 | 312 | Q15185\|TEBP_HUMAN | 84.74 | 16 | 2 | 2 | 18697 | PTGES3 | yes |
| 267 | 204 | P50395\|GDIB_HUMAN | 84.69 | 10 | 2 | 2 | 50663 | GDI2 | no |
| 159 | 140 | P26641\|EF1G_HUMAN | 84.64 | 11 | 4 | 4 | 50119 | EEF1G | yes |
| 214 | 527 | P30154\|2AAB_HUMAN | 84.55 | 6 | 2 | 1 | 66214 | PPP2R1B | no |
| 327 | 281 | O00401\|WASL_HUMAN | 84.07 | 12 | 2 | 2 | 54827 | WASL | no |
| 437 | 232 | Q96TA2\|YMEL1_HUMAN | 83.73 | 7 | 3 | 3 | 86455 | YME1L1 | no |
| 196 | 230 | P40222\|TXLNA_HUMAN | 83.6 | 8 | 2 | 2 | 61891 | TXLNA | no |
| 406 | 216 | Q01082\|SPTB2_HUMAN | 83.51 | 3 | 3 | 3 | 274608 | SPTBN1 | yes |
| 291 | 431 | P11172\|UMPS_HUMAN | 83.38 | 11 | 2 | 2 | 52222 | UMPS | yes |
| 260 | 241 | P07954\|FUMH_HUMAN | 83.02 | 10 | 2 | 2 | 54637 | FH | yes |
| 254 | 205 | Q99829\|CPNE1_HUMAN | 82.91 | 5 | 2 | 2 | 59059 | CPNE1 | yes |
| 261 | 317 | Q96CW5\|GCP3_HUMAN | 82.88 | 6 | 3 | 3 | 103571 | TUBGCP3 | yes |
| 284 | 403 | Q96AC1\|FERM2_HUMAN | 82.74 | 6 | 2 | 2 | 77861 | FERMT2 | yes |
| 275 | 178 | Q9Y678\|COPG1_HUMAN | 82.35 | 4 | 2 | 2 | 97718 | COPG1 | yes |
| 220 | 306 | O00303\|EIF3F_HUMAN | 82.33 | 10 | 2 | 2 | 37564 | EIF3F | yes |
| 317 | 1859 | O14979\|HNRDL_HUMAN | 82.07 | 8 | 2 | 2 | 46438 | HNRNPDL | yes |
| 564 | 423 | Q9BXJ9\|NAA15_HUMAN | 82.05 | 3 | 1 | 1 | 101272 | NAA15 | yes |
| 365 | 526 | P09936\|UCHL1_HUMAN | 81.93 | 10 | 2 | 2 | 24824 | UCHL1 | no |
| 330 | 283 | Q14141\|SEPT6_HUMAN | 81.25 | 12 | 2 | 2 | 49717 | Sep-06 | no |
| 372 | 635 | P08559\|ODPA_HUMAN | 81.18 | 7 | 2 | 2 | 43296 | PDHA1 | no |
| 258 | 243 | P13929\|ENOB_HUMAN | 81.11 | 7 | 2 | 1 | 46987 | ENO3 | no |

| | | | | | | | | | |
|---|---|---|---|---|---|---|---|---|---|
| 251 | 226 | Q15021\|CND1_HUMAN | 81.07 | 4 | 4 | 4 | 157182 | NCAPD2 | no |
| 124 | 210 | Q13045\|FLII_HUMAN | 80.96 | 3 | 3 | 3 | 144751 | FLII | yes |
| 385 | 952 | P62879\|GBB2_HUMAN | 80.43 | 10 | 2 | 2 | 37331 | GNB2 | yes |
| 567 | 447 | P82930\|RT34_HUMAN | 80.41 | 12 | 1 | 1 | 25650 | MRPS34 | no |
| 438 | 340 | Q9NS69\|TOM22_HUMAN | 79.68 | 20 | 2 | 2 | 15522 | TOMM22 | no |
| 271 | 442 | P23246\|SFPQ_HUMAN | 79.07 | 5 | 2 | 2 | 76150 | SFPQ | yes |
| 243 | 214 | Q15149\|PLEC_HUMAN | 79.07 | 1 | 2 | 2 | 531796 | PLEC | yes |
| 566 | 448 | P09874\|PARP1_HUMAN | 79.04 | 2 | 1 | 1 | 113084 | PARP1 | yes |
| 249 | 244 | Q01813\|PFKAP_HUMAN | 78.87 | 3 | 1 | 1 | 85596 | PFKP | yes |
| 380 | 705 | Q96AG4\|LRC59_HUMAN | 78.61 | 17 | 2 | 2 | 34930 | LRRC59 | no |
| 471 | 338 | Q9ULH0\|KDIS_HUMAN | 78.49 | 2 | 2 | 2 | 196541 | KIDINS220 | yes |
| 200 | 378 | P48735\|IDHP_HUMAN | 78.48 | 6 | 2 | 2 | 50909 | IDH2 | yes |
| 440 | 341 | P51659\|DHB4_HUMAN | 78.4 | 6 | 2 | 2 | 79686 | HSD17B4 | yes |
| 237 | 665 | P12236\|ADT3_HUMAN | 78.23 | 20 | 3 | 2 | 32866 | SLC25A6 | yes |
| 238 | 410 | Q9UMS4\|PRP19_HUMAN | 78.06 | 9 | 3 | 3 | 55181 | PRPF19 | no |
| 241 | 228 | P53618\|COPB_HUMAN | 77.92 | 4 | 2 | 2 | 107142 | COPB1 | yes |
| 235 | 592 | Q15056\|IF4H_HUMAN | 77.68 | 12 | 2 | 2 | 27385 | EIF4H | yes |
| 334 | 295 | Q3ZCQ8\|TIM50_HUMAN | 77.31 | 8 | 2 | 2 | 39646 | TIMM50 | no |
| 343 | 315 | Q14789\|GOGB1_HUMAN | 77.24 | 1 | 2 | 2 | 376019 | GOLGB1 | yes |
| 383 | 766 | Q8NE71\|ABCF1_HUMAN | 77.23 | 6 | 2 | 2 | 95926 | ABCF1 | no |
| 368 | 598 | P18124\|RL7_HUMAN | 77.02 | 5 | 2 | 2 | 29226 | RPL7 | yes |
| 454 | 473 | Q96P70\|IPO9_HUMAN | 76.79 | 2 | 2 | 2 | 115963 | IPO9 | yes |
| 360 | 509 | P60953\|CDC42_HUMAN | 76.7 | 14 | 2 | 2 | 21259 | CDC42 | yes |
| 171 | 167 | Q7L2H7\|EIF3M_HUMAN | 76.69 | 9 | 2 | 2 | 42503 | EIF3M | yes |
| 250 | 247 | Q9Y2A7\|NCKP1_HUMAN | 76.67 | 4 | 2 | 2 | 128790 | NCKAP1 | yes |
| 428 | 245 | Q9Y5B9\|SP16H_HUMAN | 76.25 | 2 | 2 | 2 | 119914 | SUPT16H | yes |
| 293 | 597 | P50148\|GNAQ_HUMAN | 76.17 | 9 | 2 | 2 | 42142 | GNAQ | no |
| 332 | 290 | P38606\|VATA_HUMAN | 75.96 | 7 | 2 | 2 | 68304 | ATP6V1A | no |
| 490 | 389 | P31689\|DNJA1_HUMAN | 75.82 | 6 | 1 | 1 | 44868 | DNAJA1 | yes |
| 496 | 379 | tr\|C9J1V9\|C9J1V9_HUMAN | 75.56 | 17 | 1 | 1 | 17018 | EEF1E1 | no |
| 496 | 441 | O43324\|MCA3_HUMAN | 75.56 | 14 | 1 | 1 | 19811 | EEF1E1 | no |
| 252 | 159 | Q96N67\|DOCK7_HUMAN | 75.35 | 2 | 2 | 2 | 242558 | DOCK7 | yes |
| 416 | 370 | Q9UHQ9\|NB5R1_HUMAN | 75.18 | 9 | 1 | 1 | 34095 | CYB5R1 | no |
| 280 | 196 | E9PAV3\|NACAM_HUMAN | 74.83 | 2 | 2 | 2 | 205419 | NACA | no |

| | | | | | | | | | |
|---|---|---|---|---|---|---|---|---|---|
| 568 | 453 | P63104\|1433Z_HUMAN | 74.8 | 8 | 1 | 1 | 27745 | YWHAZ | yes |
| 408 | 238 | P10253\|LYAG_HUMAN | 74.74 | 2 | 1 | 1 | 105324 | GAA | yes |
| 264 | 197 | P55060\|XPO2_HUMAN | 74.21 | 3 | 2 | 2 | 110417 | CSE1L | yes |
| 307 | 454 | P62195\|PRS8_HUMAN | 73.96 | 5 | 1 | 1 | 45626 | PSMC5 | yes |
| 433 | 307 | Q99729\|ROAA_HUMAN | 73.63 | 6 | 1 | 1 | 36225 | HNRNPAB | yes |
| 448 | 365 | Q8WX77\|IBPL1_HUMAN | 73.42 | 8 | 1 | 1 | 29005 | IGFBPL1 | no |
| 335 | 572 | Q9Y5V3\|MAGD1_HUMAN | 73.4 | 4 | 2 | 2 | 86161 | MAGED1 | no |
| 188 | 191 | P02768\|ALBU_HUMAN | 73.32 | 6 | 3 | 1 | 69367 | ALB | yes |
| 569 | 455 | Q7Z2W4\|ZCCHV_HUMAN | 73.29 | 2 | 1 | 1 | 101431 | ZC3HAV1 | no |
| 240 | 212 | Q09666\|AHNK_HUMAN | 72.83 | 1 | 3 | 3 | 629114 | AHNAK | no |
| 225 | 457 | P31946\|1433B_HUMAN | 72.65 | 8 | 1 | 1 | 28082 | YWHAB | yes |
| 309 | 458 | P51665\|PSMD7_HUMAN | 72.63 | 7 | 1 | 1 | 37025 | PSMD7 | no |
| 570 | 460 | Q9NP58\|ABCB6_HUMAN | 72.5 | 3 | 1 | 1 | 93886 | ABCB6 | no |
| 348 | 461 | P27348\|1433T_HUMAN | 72.16 | 8 | 1 | 1 | 27764 | YWHAQ | yes |
| 514 | 462 | P37198\|NUP62_HUMAN | 71.92 | 4 | 1 | 1 | 53255 | NUP62 | no |
| 379 | 697 | P28288\|ABCD3_HUMAN | 71.76 | 6 | 2 | 2 | 75476 | ABCD3 | yes |
| 316 | 1390 | Q08945\|SSRP1_HUMAN | 71.7 | 3 | 2 | 2 | 81075 | SSRP1 | no |
| 270 | 335 | Q5JWF2\|GNAS1_HUMAN | 71.62 | 3 | 1 | 1 | 111024 | GNAS | no |
| 571 | 464 | O14967\|CLGN_HUMAN | 71.62 | 3 | 1 | 1 | 70039 | CLGN | no |
| 382 | 725 | P51114\|FXR1_HUMAN | 71.48 | 5 | 2 | 2 | 69721 | FXR1 | yes |
| 337 | 305 | tr\|E5RI56\|E5RI56_HUMAN | 71.46 | 45 | 2 | 1 | 10271 | NA | no |
| 572 | 465 | Q9BR76\|COR1B_HUMAN | 71.25 | 6 | 1 | 1 | 54235 | CORO1B | no |
| 574 | 467 | Q8IXM3\|RM41_HUMAN | 70.77 | 19 | 1 | 1 | 15383 | MRPL41 | yes |
| 573 | 468 | P63172\|DYLT1_HUMAN | 70.73 | 16 | 1 | 1 | 12452 | DYNLT1 | no |
| 117 | 1349 | Q8TAT6\|NPL4_HUMAN | 70.67 | 4 | 2 | 2 | 68120 | NPLOC4 | yes |
| 377 | 681 | Q9NTK5\|OLA1_HUMAN | 70.52 | 7 | 2 | 2 | 44744 | OLA1 | yes |
| 338 | 599 | Q8NBJ5\|GT251_HUMAN | 70.22 | 7 | 2 | 2 | 71636 | COLGALT1 | yes |
| 412 | 299 | Q5JSZ5\|PRC2B_HUMAN | 70.07 | 1 | 1 | 1 | 242964 | PRRC2B | no |
| 266 | 198 | Q8IY17\|PLPL6_HUMAN | 69.8 | 3 | 2 | 2 | 150953 | PNPLA6 | no |
| 482 | 372 | O00411\|RPOM_HUMAN | 69.73 | 2 | 1 | 1 | 138620 | POLRMT | yes |
| 289 | 239 | O95573\|ACSL3_HUMAN | 69.46 | 5 | 2 | 2 | 80420 | ACSL3 | yes |
| 575 | 470 | P31146\|COR1A_HUMAN | 69.38 | 6 | 1 | 1 | 51026 | CORO1A | no |
| 257 | 262 | tr\|A0A1W2PQ90\|A0A1W2PQ90_HUMAN | 68.93 | 2 | 2 | 2 | 142941 | NA | no |
| 257 | 263 | Q92878\|RAD50_HUMAN | 68.93 | 2 | 2 | 2 | 153892 | RAD50 | no |

| | | | | | | | | | |
|---|---|---|---|---|---|---|---|---|---|
| 268 | 260 | P08240\|SRPRA_HUMAN | 68.81 | 4 | 2 | 2 | 69811 | SRPRA | yes |
| 463 | 882 | Q96MX6\|WDR92_HUMAN | 68.51 | 12 | 2 | 2 | 39740 | WDR92 | yes |
| 375 | 650 | P47755\|CAZA2_HUMAN | 68.45 | 16 | 2 | 2 | 32949 | CAPZA2 | no |
| 576 | 401 | O60499\|STX10_HUMAN | 68.28 | 10 | 1 | 1 | 28114 | STX10 | no |
| 392 | 733 | Q96HP0\|DOCK6_HUMAN | 68.25 | 2 | 3 | 3 | 229556 | DOCK6 | no |
| 259 | 838 | P26640\|SYVC_HUMAN | 68.1 | 5 | 3 | 3 | 140476 | VARS | yes |
| 219 | 296 | P46778\|RL21_HUMAN | 68.08 | 21 | 2 | 2 | 18565 | RPL21 | yes |
| 409 | 249 | Q13459\|MYO9B_HUMAN | 68.04 | 1 | 2 | 2 | 243398 | MYO9B | no |
| 505 | 439 | Q9BZE1\|RM37_HUMAN | 67.99 | 5 | 1 | 1 | 48118 | MRPL37 | no |
| 278 | 152 | Q9HAV4\|XPO5_HUMAN | 67.94 | 3 | 2 | 2 | 136311 | XPO5 | yes |
| 577 | 476 | P12268\|IMDH2_HUMAN | 67.62 | 4 | 1 | 1 | 55805 | IMPDH2 | yes |
| 245 | 219 | P31948\|STIP1_HUMAN | 67.61 | 3 | 1 | 1 | 62639 | STIP1 | no |
| 376 | 653 | P23921\|RIR1_HUMAN | 67.59 | 4 | 2 | 2 | 90070 | RRM1 | yes |
| 340 | 594 | Q9Y613\|FHOD1_HUMAN | 67.54 | 5 | 2 | 2 | 126551 | FHOD1 | no |
| 374 | 645 | P48444\|COPD_HUMAN | 67.43 | 9 | 2 | 2 | 57210 | ARCN1 | yes |
| 509 | 477 | Q96GA3\|LTV1_HUMAN | 67.38 | 4 | 1 | 1 | 54855 | LTV1 | no |
| 304 | 425 | P15311\|EZRI_HUMAN | 67.34 | 7 | 2 | 2 | 69413 | EZR | yes |
| 427 | 231 | Q9Y2Z4\|SYYM_HUMAN | 67.29 | 6 | 2 | 2 | 53199 | YARS2 | yes |
| 578 | 478 | Q14165\|MLEC_HUMAN | 67.26 | 8 | 1 | 1 | 32234 | MLEC | no |
| 579 | 479 | P52306\|GDS1_HUMAN | 67.07 | 3 | 1 | 1 | 66317 | RAP1GDS1 | yes |
| 391 | 323 | Q8WY21\|SORC1_HUMAN | 66.91 | 2 | 1 | 1 | 129635 | SORCS1 | no |
| 472 | 342 | O95757\|HS74L_HUMAN | 66.91 | 5 | 2 | 2 | 94512 | HSPA4L | yes |
| 580 | 480 | P53992\|SC24C_HUMAN | 66.86 | 2 | 1 | 1 | 118325 | SEC24C | yes |
| 132 | 256 | P0CG38\|POTEI_HUMAN | 66.51 | 3 | 2 | 1 | 121282 | POTEI | no |
| 370 | 607 | Q96A57\|TM230_HUMAN | 66.42 | 17 | 2 | 2 | 13188 | TMEM230 | no |
| 442 | 368 | Q15154\|PCM1_HUMAN | 66.36 | 1 | 1 | 1 | 228542 | PCM1 | yes |
| 373 | 638 | Q5T1M5\|FKB15_HUMAN | 66.23 | 4 | 2 | 2 | 133630 | FKBP15 | yes |
| 480 | 366 | O95299\|NDUAA_HUMAN | 66.13 | 6 | 1 | 1 | 40751 | NDUFA10 | no |
| 535 | 1109 | Q15393\|SF3B3_HUMAN | 66.07 | 2 | 2 | 2 | 135577 | SF3B3 | yes |
| 395 | 218 | O43747\|AP1G1_HUMAN | 65.95 | 4 | 2 | 2 | 91351 | AP1G1 | no |
| 581 | 483 | P53999\|TCP4_HUMAN | 65.87 | 19 | 1 | 1 | 14395 | SUB1 | no |
| 342 | 188 | Q15366\|PCBP2_HUMAN | 65.6 | 13 | 2 | 2 | 38580 | PCBP2 | yes |
| 389 | 744 | Q9UQE7\|SMC3_HUMAN | 65.4 | 2 | 2 | 2 | 141541 | SMC3 | yes |
| 310 | 514 | P60228\|EIF3E_HUMAN | 65.33 | 4 | 1 | 1 | 52221 | EIF3E | yes |

| | | | | | | | | | |
|---|---|---|---|---|---|---|---|---|---|
| 354 | 484 | Q9NUP9\|LIN7C_HUMAN | 65.32 | 10 | 1 | 1 | 21834 | LIN7C | yes |
| 582 | 413 | Q9UNM6\|PSD13_HUMAN | 65.31 | 6 | 1 | 1 | 42946 | PSMD13 | yes |
| 483 | 374 | Q8NEV1\|CSK23_HUMAN | 65.25 | 6 | 1 | 1 | 45220 | CSNK2A3 | no |
| 483 | 375 | P68400\|CSK21_HUMAN | 65.25 | 6 | 1 | 1 | 45144 | CSNK2A1 | yes |
| 583 | 485 | P42166\|LAP2A_HUMAN | 65.2 | 3 | 1 | 1 | 75492 | TMPO | yes |
| 473 | 346 | P62424\|RL7A_HUMAN | 64.88 | 6 | 1 | 1 | 29996 | RPL7A | yes |
| 584 | 486 | Q9NP72\|RAB18_HUMAN | 64.84 | 9 | 1 | 1 | 22977 | RAB18 | no |
| 228 | 487 | P62841\|RS15_HUMAN | 64.78 | 13 | 1 | 1 | 17040 | RPS15 | yes |
| 507 | 548 | Q12765\|SCRN1_HUMAN | 64.78 | 8 | 2 | 2 | 46382 | SCRN1 | yes |
| 536 | 1113 | Q8N6R0\|MET13_HUMAN | 64.67 | 4 | 2 | 2 | 78768 | METTL13 | no |
| 288 | 233 | Q13085\|ACACA_HUMAN | 64.49 | 1 | 2 | 2 | 265551 | ACACA | yes |
| 336 | 298 | P26885\|FKBP2_HUMAN | 64.44 | 23 | 2 | 2 | 15649 | FKBP2 | no |
| 443 | 345 | P07205\|PGK2_HUMAN | 64.36 | 6 | 2 | 2 | 44796 | PGK2 | no |
| 443 | 540 | P00558\|PGK1_HUMAN | 56.62 | 4 | 1 | 1 | 44615 | PGK1 | yes |
| 585 | 491 | Q13438\|OS9_HUMAN | 64.16 | 3 | 1 | 1 | 75562 | OS9 | no |
| 300 | 373 | P20042\|IF2B_HUMAN | 64.15 | 6 | 1 | 1 | 38388 | EIF2S2 | no |
| 586 | 490 | P10645\|CMGA_HUMAN | 64.07 | 4 | 1 | 1 | 50688 | CHGA | yes |
| 419 | 804 | Q9BTW9\|TBCD_HUMAN | 63.83 | 2 | 1 | 1 | 132600 | TBCD | yes |
| 357 | 492 | P11177\|ODPB_HUMAN | 63.79 | 4 | 1 | 1 | 39233 | PDHB | yes |
| 587 | 493 | Q5VSL9\|STRP1_HUMAN | 63.56 | 2 | 1 | 1 | 95576 | STRIP1 | yes |
| 256 | 215 | P54886\|P5CS_HUMAN | 63.51 | 4 | 2 | 2 | 87302 | ALDH18A1 | yes |
| 152 | 371 | Q96A33\|CCD47_HUMAN | 63.46 | 5 | 1 | 1 | 55874 | CCDC47 | no |
| 552 | 1849 | O75663\|TIPRL_HUMAN | 63.42 | 8 | 1 | 1 | 31444 | TIPRL | yes |
| 229 | 494 | P62750\|RL23A_HUMAN | 63.33 | 10 | 1 | 1 | 17695 | RPL23A | yes |
| 263 | 186 | O00232\|PSD12_HUMAN | 63.12 | 6 | 2 | 2 | 52904 | PSMD12 | yes |
| 434 | 319 | Q9NVH1\|DJC11_HUMAN | 62.87 | 3 | 1 | 1 | 63278 | DNAJC11 | no |
| 411 | 297 | Q8WUM4\|PDC6I_HUMAN | 62.42 | 4 | 1 | 1 | 96023 | PDCD6IP | yes |
| 588 | 497 | Q96JC1\|VPS39_HUMAN | 62.41 | 2 | 1 | 1 | 101809 | VPS39 | no |
| 297 | 359 | P27695\|APEX1_HUMAN | 62.4 | 5 | 1 | 1 | 35555 | APEX1 | yes |
| 359 | 498 | Q06787\|FMR1_HUMAN | 62.26 | 2 | 1 | 1 | 71175 | FMR1 | yes |
| 589 | 499 | O76021\|RL1D1_HUMAN | 62.09 | 4 | 1 | 1 | 54973 | RSL1D1 | yes |
| 590 | 500 | O94903\|PLPHP_HUMAN | 61.89 | 5 | 1 | 1 | 30344 | PLPBP | yes |
| 441 | 334 | Q15067\|ACOX1_HUMAN | 61.86 | 2 | 1 | 1 | 74424 | ACOX1 | no |
| 513 | 501 | P00338\|LDHA_HUMAN | 61.61 | 5 | 1 | 1 | 36689 | LDHA | yes |

| | | | | | | | | | |
|---|---|---|---|---|---|---|---|---|---|
| 591 | 502 | P53582\|MAP11_HUMAN | 61.54 | 5 | 1 | 1 | 43215 | METAP1 | no |
| 592 | 503 | Q9ULC4\|MCTS1_HUMAN | 61.46 | 9 | 1 | 1 | 20555 | MCTS1 | yes |
| 593 | 504 | O96005\|CLPT1_HUMAN | 61.29 | 3 | 1 | 1 | 76097 | CLPTM1 | no |
| 294 | 325 | Q9UPQ0\|LIMC1_HUMAN | 60.94 | 2 | 2 | 2 | 121867 | LIMCH1 | yes |
| 286 | 225 | P11142\|HSP7C_HUMAN | 60.9 | 5 | 2 | 1 | 70898 | HSPA8 | yes |
| 378 | 683 | Q9Y3B3\|TMED7_HUMAN | 60.52 | 9 | 1 | 1 | 25172 | TMED7 | no |
| 378 | 682 | tr\|A0A0A6YYA0\|A0A0A6YYA0_HUMAN | 60.52 | 11 | 1 | 1 | 21233 | TMED7 | no |
| 595 | 400 | Q9NP92\|RT30_HUMAN | 60.35 | 5 | 1 | 1 | 50365 | MRPS30 | yes |
| 484 | 376 | Q14697\|GANAB_HUMAN | 60.35 | 2 | 1 | 1 | 106874 | GANAB | yes |
| 381 | 721 | Q9Y3F4\|STRAP_HUMAN | 60.15 | 13 | 2 | 2 | 38438 | STRAP | no |
| 596 | 513 | P09543\|CN37_HUMAN | 60.04 | 5 | 1 | 1 | 47579 | CNP | no |
| 537 | 1122 | P15313\|VATB1_HUMAN | 59.98 | 5 | 2 | 2 | 56833 | ATP6V1B1 | no |
| 597 | 515 | Q16543\|CDC37_HUMAN | 59.84 | 6 | 1 | 1 | 44468 | CDC37 | yes |
| 362 | 516 | A1L0T0\|ILVBL_HUMAN | 59.78 | 3 | 1 | 1 | 67868 | ILVBL | no |
| 501 | 418 | Q15031\|SYLM_HUMAN | 59.72 | 3 | 1 | 1 | 101976 | LARS2 | yes |
| 598 | 517 | P10515\|ODP2_HUMAN | 59.65 | 2 | 1 | 1 | 68997 | DLAT | yes |
| 599 | 518 | Q16566\|KCC4_HUMAN | 59.38 | 3 | 1 | 1 | 51926 | CAMK4 | no |
| 600 | 519 | P28289\|TMOD1_HUMAN | 59.36 | 5 | 1 | 1 | 40569 | TMOD1 | no |
| 233 | 520 | Q9P0J0\|NDUAD_HUMAN | 59.34 | 9 | 1 | 1 | 16698 | NDUFA13 | no |
| 601 | 521 | Q96EY5\|MB12A_HUMAN | 59.22 | 6 | 1 | 1 | 28783 | MVB12A | no |
| 363 | 522 | P0DN76\|U2AF5_HUMAN | 59.19 | 8 | 1 | 1 | 27872 | U2AF1L5 | no |
| 363 | 523 | Q01081\|U2AF1_HUMAN | 59.19 | 8 | 1 | 1 | 27872 | U2AF1 | yes |
| 444 | 347 | P18621\|RL17_HUMAN | 59.12 | 9 | 1 | 1 | 21397 | RPL17 | no |
| 444 | 348 | tr\|A0A0A6YYL6\|A0A0A6YYL6_HUMAN | 59.12 | 7 | 1 | 1 | 26373 | RPL17 | no |
| 553 | 2006 | O75131\|CPNE3_HUMAN | 59.02 | 4 | 1 | 1 | 60131 | CPNE3 | no |
| 410 | 261 | A4D1E9\|GTPBA_HUMAN | 58.88 | 6 | 1 | 1 | 42933 | GTPBP10 | no |
| 602 | 524 | P40938\|RFC3_HUMAN | 58.86 | 4 | 1 | 1 | 40556 | RFC3 | yes |
| 542 | 1230 | Q14694\|UBP10_HUMAN | 58.77 | 3 | 1 | 1 | 87134 | USP10 | no |
| 558 | 2380 | Q13347\|EIF3I_HUMAN | 58.67 | 9 | 1 | 1 | 36502 | EIF3I | no |
| 495 | 397 | Q5XKP0\|MIC13_HUMAN | 58.48 | 19 | 1 | 1 | 13087 | MIC13 | no |
| 402 | 847 | Q9NRY4\|RHG35_HUMAN | 58.44 | 2 | 1 | 1 | 170513 | ARHGAP35 | no |
| 603 | 402 | P52272\|HNRPM_HUMAN | 58.36 | 2 | 1 | 1 | 77516 | HNRNPM | yes |
| 604 | 528 | O75340\|PDCD6_HUMAN | 58.2 | 16 | 1 | 1 | 21868 | PDCD6 | no |
| 605 | 529 | P07195\|LDHB_HUMAN | 58.1 | 4 | 1 | 1 | 36639 | LDHB | yes |

| | | | | | | | | | |
|---|---|---|---|---|---|---|---|---|---|
| 606 | 531 | Q8WXF1\|PSPC1_HUMAN | 58.05 | 4 | 1 | 1 | 58744 | PSPC1 | yes |
| 399 | 776 | A3KMH1\|VWA8_HUMAN | 58.05 | 1 | 1 | 1 | 214823 | VWA8 | no |
| 281 | 199 | Q99714\|HCD2_HUMAN | 58.02 | 11 | 2 | 2 | 26923 | HSD17B10 | yes |
| 549 | 1809 | Q92804\|RBP56_HUMAN | 57.87 | 6 | 1 | 1 | 61830 | TAF15 | no |
| 607 | 532 | P52788\|SPSY_HUMAN | 57.75 | 6 | 1 | 1 | 41268 | SMS | no |
| 608 | 533 | Q16836\|HCDH_HUMAN | 57.43 | 7 | 1 | 1 | 34294 | HADH | no |
| 609 | 534 | P54578\|UBP14_HUMAN | 57.37 | 5 | 1 | 1 | 56069 | USP14 | no |
| 610 | 535 | Q15005\|SPCS2_HUMAN | 57.26 | 8 | 1 | 1 | 25003 | SPCS2 | no |
| 611 | 536 | Q13151\|ROA0_HUMAN | 57.19 | 7 | 1 | 1 | 30841 | HNRNPA0 | no |
| 512 | 538 | P22570\|ADRO_HUMAN | 56.98 | 5 | 1 | 1 | 53837 | FDXR | no |
| 467 | 782 | Q6UB35\|C1TM_HUMAN | 56.95 | 1 | 1 | 1 | 105790 | MTHFD1L | yes |
| 557 | 2349 | Q9H2M9\|RBGPR_HUMAN | 56.85 | 2 | 1 | 1 | 155984 | RAB3GAP2 | yes |
| 425 | 1752 | Q7L8L6\|FAKD5_HUMAN | 56.85 | 2 | 1 | 1 | 86574 | FASTKD5 | yes |
| 612 | 539 | P51398\|RT29_HUMAN | 56.79 | 5 | 1 | 1 | 45566 | DAP3 | no |
| 538 | 1132 | Q96QK1\|VPS35_HUMAN | 56.65 | 3 | 2 | 2 | 91707 | VPS35 | yes |
| 510 | 550 | Q15029\|U5S1_HUMAN | 56.58 | 1 | 1 | 1 | 109436 | EFTUD2 | yes |
| 613 | 542 | O75718\|CRTAP_HUMAN | 56.54 | 4 | 1 | 1 | 46562 | CRTAP | yes |
| 614 | 541 | O43681\|ASNA_HUMAN | 56.52 | 5 | 1 | 1 | 38793 | ASNA1 | yes |
| 550 | 1812 | Q9NZ01\|TECR_HUMAN | 56.43 | 7 | 1 | 1 | 36034 | TECR | yes |
| 615 | 543 | Q9BYD2\|RM09_HUMAN | 56.42 | 6 | 1 | 1 | 30243 | MRPL9 | yes |
| 616 | 544 | P50579\|MAP2_HUMAN | 56.3 | 3 | 1 | 1 | 52892 | METAP2 | yes |
| 344 | 320 | Q9BSH4\|TACO1_HUMAN | 56.26 | 11 | 2 | 2 | 32477 | TACO1 | no |
| 617 | 545 | Q9H8Y8\|GORS2_HUMAN | 56.16 | 12 | 1 | 1 | 47145 | GORASP2 | yes |
| 432 | 304 | Q14643\|ITPR1_HUMAN | 56.09 | 1 | 1 | 1 | 313928 | ITPR1 | yes |
| 618 | 547 | O15372\|EIF3H_HUMAN | 56.07 | 5 | 1 | 1 | 39930 | EIF3H | no |
| 413 | 321 | Q6ZRI8\|RHG36_HUMAN | 55.91 | 4 | 1 | 1 | 61664 | ARHGAP36 | no |
| 431 | 292 | Q16537\|2A5E_HUMAN | 55.87 | 3 | 1 | 1 | 54699 | PPP2R5E | yes |
| 619 | 549 | Q96EY1\|DNJA3_HUMAN | 55.85 | 3 | 1 | 1 | 52489 | DNAJA3 | no |
| 445 | 349 | Q9Y383\|LC7L2_HUMAN | 55.73 | 5 | 2 | 2 | 46514 | LUC7L2 | yes |
| 439 | 333 | P61019\|RAB2A_HUMAN | 55.54 | 7 | 1 | 1 | 23546 | RAB2A | yes |
| 620 | 551 | P07948\|LYN_HUMAN | 55.41 | 4 | 1 | 1 | 58574 | LYN | no |
| 621 | 424 | Q9NYL9\|TMOD3_HUMAN | 55.33 | 6 | 1 | 1 | 39595 | TMOD3 | yes |
| 622 | 552 | Q9UBT2\|SAE2_HUMAN | 55.33 | 3 | 1 | 1 | 71224 | UBA2 | yes |
| 623 | 553 | P51648\|AL3A2_HUMAN | 55.26 | 4 | 1 | 1 | 54848 | ALDH3A2 | no |

| | | | | | | | | | |
|---|---|---|---|---|---|---|---|---|---|
| 624 | 555 | Q9BSF4\|TIM29_HUMAN | 55.05 | 9 | 1 | 1 | 29233 | TIMM29 | no |
| 625 | 556 | Q9NPQ8\|RIC8A_HUMAN | 54.92 | 4 | 1 | 1 | 59710 | RIC8A | yes |
| 464 | 980 | Q2M389\|WASC4_HUMAN | 54.65 | 2 | 2 | 2 | 136403 | WASHC4 | yes |
| 478 | 361 | P21912\|SDHB_HUMAN | 54.64 | 5 | 1 | 1 | 31630 | SDHB | no |
| 449 | 421 | O95793\|STAU1_HUMAN | 54.6 | 2 | 1 | 1 | 63182 | STAU1 | yes |
| 474 | 351 | O15357\|SHIP2_HUMAN | 54.55 | 2 | 1 | 1 | 138599 | INPPL1 | yes |
| 560 | 2566 | B2RXH8\|HNRC2_HUMAN | 54.45 | 6 | 1 | 1 | 32072 | HNRNPCL2 | no |
| 560 | 2567 | O60812\|HNRC1_HUMAN | 54.45 | 6 | 1 | 1 | 32142 | HNRNPCL1 | no |
| 511 | 426 | O14531\|DPYL4_HUMAN | 54.45 | 3 | 1 | 1 | 61878 | DPYSL4 | no |
| 626 | 561 | tr\|F5H423\|F5H423_HUMAN | 54.34 | 9 | 1 | 1 | 23346 | NA | no |
| 626 | 559 | P61204\|ARF3_HUMAN | 54.34 | 10 | 1 | 1 | 20601 | ARF3 | no |
| 626 | 560 | P84077\|ARF1_HUMAN | 54.34 | 10 | 1 | 1 | 20697 | ARF1 | yes |
| 626 | 557 | P84085\|ARF5_HUMAN | 54.34 | 10 | 1 | 1 | 20530 | ARF5 | no |
| 626 | 558 | P18085\|ARF4_HUMAN | 54.34 | 10 | 1 | 1 | 20511 | ARF4 | no |
| 627 | 562 | Q96HY6\|DDRGK_HUMAN | 54.28 | 4 | 1 | 1 | 35611 | DDRGK1 | no |
| 234 | 563 | Q9C0H2\|TTYH3_HUMAN | 54.27 | 3 | 1 | 1 | 57545 | TTYH3 | no |
| 541 | 1225 | P41250\|GARS_HUMAN | 54.16 | 3 | 1 | 1 | 83166 | GARS | yes |
| 407 | 223 | Q9UBF2\|COPG2_HUMAN | 54.02 | 2 | 1 | 1 | 97622 | COPG2 | yes |
| 628 | 564 | Q9Y676\|RT18B_HUMAN | 53.97 | 13 | 1 | 1 | 29396 | MRPS18B | no |
| 305 | 433 | Q14008\|CKAP5_HUMAN | 53.91 | 1 | 2 | 2 | 225493 | CKAP5 | yes |
| 629 | 565 | Q04760\|LGUL_HUMAN | 53.89 | 9 | 1 | 1 | 20778 | GLO1 | yes |
| 508 | 567 | Q9H9E3\|COG4_HUMAN | 53.86 | 2 | 1 | 1 | 89083 | COG4 | no |
| 506 | 440 | Q9UBQ6\|EXTL2_HUMAN | 53.79 | 5 | 1 | 1 | 37466 | EXTL2 | no |
| 414 | 344 | Q96RP9\|EFGM_HUMAN | 53.49 | 5 | 2 | 2 | 83472 | GFM1 | no |
| 630 | 568 | P61586\|RHOA_HUMAN | 53.44 | 13 | 1 | 1 | 21768 | RHOA | no |
| 415 | 356 | P36542\|ATPG_HUMAN | 53.22 | 4 | 1 | 1 | 32996 | ATP5F1C | yes |
| 631 | 569 | Q9NVA2\|SEP11_HUMAN | 53.21 | 5 | 1 | 1 | 49398 | Sep-11 | no |
| 632 | 570 | Q9BRJ2\|RM45_HUMAN | 53.19 | 5 | 1 | 1 | 35351 | MRPL45 | no |
| 405 | 1099 | O60313\|OPA1_HUMAN | 53.17 | 2 | 1 | 1 | 111630 | OPA1 | yes |
| 491 | 391 | P07602\|SAP_HUMAN | 53.11 | 3 | 1 | 1 | 58113 | PSAP | no |
| 504 | 438 | Q9BWS9\|CHID1_HUMAN | 53.09 | 3 | 1 | 1 | 44941 | CHID1 | no |
| 367 | 573 | P06899\|H2B1J_HUMAN | 52.95 | 12 | 1 | 1 | 13904 | HIST1H2BJ | no |
| 367 | 574 | Q99880\|H2B1L_HUMAN | 52.95 | 12 | 1 | 1 | 13952 | HIST1H2BL | no |
| 367 | 575 | O60814\|H2B1K_HUMAN | 52.95 | 12 | 1 | 1 | 13890 | HIST1H2BK | no |

| | | | | | | | | | |
|---|---|---|---|---|---|---|---|---|---|
| 367 | 576 | Q8N257\|H2B3B_HUMAN | 52.95 | 12 | 1 | 1 | 13908 | HIST3H2BB | no |
| 367 | 577 | Q16778\|H2B2E_HUMAN | 52.95 | 12 | 1 | 1 | 13920 | HIST2H2BE | no |
| 367 | 578 | Q99879\|H2B1M_HUMAN | 52.95 | 12 | 1 | 1 | 13989 | HIST1H2BM | no |
| 367 | 579 | P58876\|H2B1D_HUMAN | 52.95 | 12 | 1 | 1 | 13936 | HIST1H2BD | no |
| 367 | 580 | Q93079\|H2B1H_HUMAN | 52.95 | 12 | 1 | 1 | 13892 | HIST1H2BH | no |
| 367 | 581 | P23527\|H2B1O_HUMAN | 52.95 | 12 | 1 | 1 | 13906 | HIST1H2BO | no |
| 367 | 582 | P33778\|H2B1B_HUMAN | 52.95 | 12 | 1 | 1 | 13950 | HIST1H2BB | no |
| 367 | 583 | Q5QNW6\|H2B2F_HUMAN | 52.95 | 12 | 1 | 1 | 13920 | HIST2H2BF | no |
| 367 | 584 | Q99877\|H2B1N_HUMAN | 52.95 | 12 | 1 | 1 | 13922 | HIST1H2BN | no |
| 367 | 585 | P57053\|H2BFS_HUMAN | 52.95 | 12 | 1 | 1 | 13944 | H2BFS | no |
| 367 | 586 | P62807\|H2B1C_HUMAN | 52.95 | 12 | 1 | 1 | 13906 | HIST1H2BC | no |
| 633 | 587 | Q7Z460\|CLAP1_HUMAN | 52.9 | 1 | 1 | 1 | 169450 | CLASP1 | yes |
| 346 | 339 | Q02790\|FKBP4_HUMAN | 52.81 | 9 | 2 | 2 | 51805 | FKBP4 | yes |
| 635 | 590 | P05455\|LA_HUMAN | 52.78 | 3 | 1 | 1 | 46837 | SSB | yes |
| 634 | 589 | P49591\|SYSC_HUMAN | 52.77 | 4 | 1 | 1 | 58777 | SARS | no |
| 636 | 591 | Q9H3U1\|UN45A_HUMAN | 52.73 | 2 | 1 | 1 | 103077 | UNC45A | no |
| 489 | 388 | P03886\|NU1M_HUMAN | 52.4 | 6 | 1 | 1 | 35661 | MT | no |
| 563 | 417 | O76031\|CLPX_HUMAN | 52.32 | 2 | 1 | 1 | 69224 | CLPX | yes |
| 637 | 593 | O00165\|HAX1_HUMAN | 52.31 | 11 | 1 | 1 | 31621 | HAX1 | no |
| 638 | 595 | Q9Y6C2\|EMIL1_HUMAN | 52.29 | 2 | 1 | 1 | 106695 | EMILIN1 | no |
| 639 | 596 | P09622\|DLDH_HUMAN | 52.27 | 3 | 1 | 1 | 54177 | DLD | yes |
| 466 | 1010 | Q9UMS6\|SYNP2_HUMAN | 52.13 | 1 | 1 | 1 | 117514 | SYNPO2 | no |
| 369 | 604 | Q9Y224\|RTRAF_HUMAN | 52.03 | 8 | 1 | 1 | 28068 | RTRAF | no |
| 436 | 329 | A6NJ78\|MET15_HUMAN | 51.73 | 4 | 1 | 1 | 46121 | METTL15 | yes |
| 640 | 600 | P46940\|IQGA1_HUMAN | 51.67 | 1 | 1 | 1 | 189251 | IQGAP1 | no |
| 476 | 355 | Q16795\|NDUA9_HUMAN | 51.5 | 5 | 1 | 1 | 42510 | NDUFA9 | no |
| 485 | 377 | P49754\|VPS41_HUMAN | 51.46 | 3 | 1 | 1 | 98566 | VPS41 | no |
| 756 | 2519 | P16435\|NCPR_HUMAN | 51.45 | 4 | 1 | 1 | 76690 | POR | yes |
| 641 | 601 | O14980\|XPO1_HUMAN | 51.41 | 2 | 1 | 1 | 123386 | XPO1 | no |
| 554 | 2270 | Q14254\|FLOT2_HUMAN | 51.36 | 5 | 1 | 1 | 47064 | FLOT2 | no |
| 475 | 353 | O15145\|ARPC3_HUMAN | 51.35 | 7 | 1 | 1 | 20547 | ARPC3 | yes |
| 426 | 1805 | P30041\|PRDX6_HUMAN | 51.31 | 8 | 1 | 1 | 25035 | PRDX6 | yes |
| 642 | 602 | Q9UKK9\|NUDT5_HUMAN | 51.22 | 7 | 1 | 1 | 24328 | NUDT5 | no |
| 643 | 603 | Q9BW62\|KATL1_HUMAN | 51.17 | 3 | 1 | 1 | 55392 | KATNAL1 | no |

| | | | | | | | | | |
|---|---|---|---|---|---|---|---|---|---|
| 644 | 605 | Q9H0A8\|COMD4_HUMAN | 51.08 | 11 | 1 | 1 | 21764 | COMMD4 | no |
| 339 | 311 | Q8N163\|CCAR2_HUMAN | 51.05 | 4 | 2 | 2 | 102902 | CCAR2 | no |
| 645 | 606 | Q8WU90\|ZC3HF_HUMAN | 51.03 | 6 | 1 | 1 | 48603 | ZC3H15 | no |
| 646 | 608 | Q08AM6\|VAC14_HUMAN | 51 | 2 | 1 | 1 | 87973 | VAC14 | yes |
| 430 | 258 | Q6P158\|DHX57_HUMAN | 50.95 | 1 | 1 | 1 | 155604 | DHX57 | yes |
| 497 | 331 | Q02809\|PLOD1_HUMAN | 50.94 | 2 | 1 | 1 | 83550 | PLOD1 | yes |
| 450 | 422 | Q9BXI3\|5NT1A_HUMAN | 50.82 | 5 | 1 | 1 | 41021 | NT5C1A | no |
| 647 | 610 | O43347\|MSI1H_HUMAN | 50.81 | 5 | 1 | 1 | 39125 | MSI1 | no |
| 648 | 611 | O75390\|CISY_HUMAN | 50.8 | 3 | 1 | 1 | 51712 | CS | yes |
| 446 | 354 | Q12906\|ILF3_HUMAN | 50.55 | 1 | 1 | 1 | 95339 | ILF3 | yes |
| 446 | 616 | Q96SI9\|STRBP_HUMAN | 50.55 | 2 | 1 | 1 | 73653 | STRBP | yes |
| 371 | 613 | P62263\|RS14_HUMAN | 50.52 | 13 | 1 | 1 | 16273 | RPS14 | no |
| 649 | 614 | Q99470\|SDF2_HUMAN | 50.46 | 11 | 1 | 1 | 23026 | SDF2 | no |
| 479 | 615 | P04004\|VTNC_HUMAN | 50.4 | 3 | 1 | 1 | 54306 | VTN | no |
| 396 | 259 | Q13740\|CD166_HUMAN | 49.9 | 2 | 1 | 1 | 65102 | ALCAM | no |
| 651 | 620 | P07858\|CATB_HUMAN | 49.67 | 4 | 1 | 1 | 37822 | CTSB | yes |
| 650 | 619 | Q15036\|SNX17_HUMAN | 49.66 | 3 | 1 | 1 | 52901 | SNX17 | no |
| 435 | 328 | O00116\|ADAS_HUMAN | 49.1 | 3 | 1 | 1 | 72912 | AGPS | yes |
| 652 | 621 | Q9Y3B4\|SF3B6_HUMAN | 49.03 | 10 | 1 | 1 | 14585 | SF3B6 | yes |
| 653 | 623 | Q96JB2\|COG3_HUMAN | 48.94 | 1 | 1 | 1 | 94096 | COG3 | yes |
| 498 | 404 | Q02218\|ODO1_HUMAN | 48.77 | 1 | 1 | 1 | 115935 | OGDH | yes |
| 345 | 327 | O75306\|NDUS2_HUMAN | 48.73 | 6 | 2 | 2 | 52546 | NDUFS2 | no |
| 555 | 2275 | P06400\|RB_HUMAN | 48.64 | 2 | 1 | 1 | 106159 | RB1 | no |
| 654 | 624 | P25325\|THTM_HUMAN | 48.61 | 8 | 1 | 1 | 33178 | MPST | no |
| 655 | 625 | P24539\|AT5F1_HUMAN | 48.45 | 5 | 1 | 1 | 28909 | ATP5PB | no |
| 656 | 627 | tr\|H3BVE0\|H3BVE0_HUMAN | 48.29 | 6 | 1 | 1 | 27673 | BOLA2 | no |
| 656 | 626 | Q9H3K6\|BOLA2_HUMAN | 48.29 | 19 | 1 | 1 | 10117 | BOLA2 | no |
| 657 | 628 | Q96H79\|ZCCHL_HUMAN | 48.19 | 11 | 1 | 1 | 32962 | ZC3HAV1L | no |
| 658 | 629 | Q9H9J2\|RM44_HUMAN | 48.11 | 4 | 1 | 1 | 37535 | MRPL44 | yes |
| 659 | 630 | Q96A35\|RM24_HUMAN | 47.9 | 6 | 1 | 1 | 24915 | MRPL24 | yes |
| 503 | 434 | P62834\|RAP1A_HUMAN | 47.78 | 7 | 1 | 1 | 20987 | RAP1A | no |
| 503 | 435 | A6NIZ1\|RP1BL_HUMAN | 47.78 | 7 | 1 | 1 | 20925 | NA | no |
| 503 | 436 | P61224\|RAP1B_HUMAN | 47.78 | 7 | 1 | 1 | 20825 | RAP1B | no |
| 660 | 633 | tr\|E7EQ34\|E7EQ34_HUMAN | 47.62 | 7 | 1 | 1 | 26864 | NA | no |

| | | | | | | | | | |
|---|---|---|---|---|---|---|---|---|---|
| 660 | 632 | O14653\|GOSR2_HUMAN | 47.62 | 8 | 1 | 1 | 24775 | GOSR2 | no |
| 477 | 357 | Q01105\|SET_HUMAN | 47.58 | 5 | 1 | 1 | 33489 | SET | no |
| 477 | 358 | P0DME0\|SETLP_HUMAN | 47.58 | 5 | 1 | 1 | 34882 | SETSIP | no |
| 661 | 634 | P23528\|COF1_HUMAN | 47.4 | 8 | 1 | 1 | 18502 | CFL1 | yes |
| 500 | 411 | P52732\|KIF11_HUMAN | 47.01 | 1 | 1 | 1 | 119159 | KIF11 | yes |
| 551 | 1837 | O94905\|ERLN2_HUMAN | 46.43 | 7 | 1 | 1 | 37840 | ERLIN2 | no |
| 562 | 2842 | Q9NTJ3\|SMC4_HUMAN | 46.37 | 1 | 1 | 1 | 147182 | SMC4 | no |
| 663 | 637 | O75396\|SC22B_HUMAN | 46.27 | 5 | 1 | 1 | 24593 | SEC22B | no |
| 662 | 636 | O15031\|PLXB2_HUMAN | 46.2 | 1 | 1 | 1 | 205126 | PLXNB2 | no |
| 469 | 2327 | Q00534\|CDK6_HUMAN | 46.01 | 5 | 1 | 1 | 36938 | CDK6 | yes |
| 666 | 642 | Q9Y2R5\|RT17_HUMAN | 45.93 | 7 | 1 | 1 | 14502 | MRPS17 | no |
| 666 | 644 | tr\|I3L0E3\|I3L0E3_HUMAN | 45.93 | 4 | 1 | 1 | 25720 | hCG_1984214 | no |
| 664 | 640 | O14950\|ML12B_HUMAN | 45.41 | 12 | 1 | 1 | 19779 | MYL12B | no |
| 664 | 639 | P19105\|ML12A_HUMAN | 45.41 | 12 | 1 | 1 | 19794 | MYL12A | no |
| 665 | 641 | P46781\|RS9_HUMAN | 45.3 | 5 | 1 | 1 | 22591 | RPS9 | no |
| 470 | 255 | Q9UFG5\|CS025_HUMAN | 45.28 | 19 | 1 | 1 | 12878 | C19orf25 | no |
| 533 | 1024 | O75746\|CMC1_HUMAN | 45.02 | 4 | 2 | 1 | 74762 | SLC25A12 | no |
| 561 | 2665 | Q12849\|GRSF1_HUMAN | 44.94 | 6 | 1 | 1 | 53126 | GRSF1 | no |
| 548 | 1710 | P15170\|ERF3A_HUMAN | 44.76 | 2 | 1 | 1 | 55756 | GSPT1 | no |
| 667 | 646 | O43920\|NDUS5_HUMAN | 44.49 | 11 | 1 | 1 | 12517 | NDUFS5 | no |
| 543 | 1311 | Q6PML9\|ZNT9_HUMAN | 44.3 | 2 | 1 | 1 | 63515 | SLC30A9 | no |
| 492 | 392 | Q9Y4G6\|TLN2_HUMAN | 44.07 | 1 | 1 | 1 | 271611 | TLN2 | no |
| 668 | 647 | P56556\|NDUA6_HUMAN | 44.07 | 12 | 1 | 1 | 15137 | NDUFA6 | no |
| 669 | 649 | tr\|F5H5P2\|F5H5P2_HUMAN | 44.03 | 3 | 1 | 1 | 54203 | NA | no |
| 669 | 648 | P12694\|ODBA_HUMAN | 44.03 | 3 | 1 | 1 | 50471 | BCKDHA | no |
| 452 | 416 | P11498\|PYC_HUMAN | 43.97 | 1 | 1 | 1 | 129634 | PC | yes |
| 487 | 383 | P02749\|APOH_HUMAN | 43.54 | 3 | 1 | 1 | 38298 | APOH | no |
| 670 | 652 | tr\|U3KQV3\|U3KQV3_HUMAN | 43.49 | 9 | 1 | 1 | 30342 | NA | no |
| 670 | 651 | P08134\|RHOC_HUMAN | 43.49 | 12 | 1 | 1 | 22006 | RHOC | no |
| 671 | 654 | O15020\|SPTN2_HUMAN | 43.19 | 1 | 1 | 1 | 271323 | SPTBN2 | no |
| 546 | 1658 | Q14738\|2A5D_HUMAN | 43.11 | 2 | 1 | 1 | 69992 | PPP2R5D | no |
| 672 | 661 | Q9BVK6\|TMED9_HUMAN | 42.91 | 9 | 1 | 1 | 27277 | TMED9 | no |
| 673 | 662 | P61604\|CH10_HUMAN | 42.86 | 14 | 1 | 1 | 10932 | HSPE1 | yes |
| 673 | 663 | tr\|S4R3N1\|S4R3N1_HUMAN | 42.86 | 5 | 1 | 1 | 29737 | HSPE1 | yes |

| | | | | | | | | | |
|---|---|---|---|---|---|---|---|---|---|
| 674 | 664 | P07196\|NFL_HUMAN | 42.73 | 3 | 1 | 1 | 61517 | NEFL | no |
| 544 | 1426 | Q9H361\|PABP3_HUMAN | 42.67 | 5 | 1 | 1 | 70031 | PABPC3 | no |
| 675 | 666 | P51553\|IDH3G_HUMAN | 42.41 | 6 | 1 | 1 | 42794 | IDH3G | no |
| 494 | 396 | P07686\|HEXB_HUMAN | 42 | 5 | 2 | 2 | 63111 | HEXB | no |
| 559 | 2500 | P0CG08\|GPHRB_HUMAN | 41.96 | 6 | 1 | 1 | 52917 | GPR89B | no |
| 559 | 2501 | B7ZAQ6\|GPHRA_HUMAN | 41.96 | 6 | 1 | 1 | 52917 | GPR89A | no |
| 676 | 667 | O95373\|IPO7_HUMAN | 41.94 | 1 | 1 | 1 | 119516 | IPO7 | no |
| 677 | 668 | P17858\|PFKAL_HUMAN | 41.91 | 2 | 1 | 1 | 85018 | PFKL | yes |
| 306 | 437 | P62314\|SMD1_HUMAN | 41.83 | 17 | 1 | 1 | 13282 | SNRPD1 | yes |
| 481 | 367 | Q9H078\|CLPB_HUMAN | 41.74 | 3 | 1 | 1 | 78729 | CLPB | yes |
| 447 | 364 | P31040\|SDHA_HUMAN | 41.63 | 2 | 1 | 1 | 72692 | SDHA | yes |
| 678 | 672 | P0CG48\|UBC_HUMAN | 41.54 | 2 | 1 | 1 | 77039 | UBC | no |
| 678 | 671 | P0CG47\|UBB_HUMAN | 41.54 | 7 | 1 | 1 | 25762 | UBB | no |
| 678 | 670 | P62979\|RS27A_HUMAN | 41.54 | 10 | 1 | 1 | 17965 | RPS27A | no |
| 678 | 669 | P62987\|RL40_HUMAN | 41.54 | 12 | 1 | 1 | 14728 | UBA52 | yes |
| 515 | 674 | Q8TEX9\|IPO4_HUMAN | 41.43 | 1 | 1 | 1 | 118715 | IPO4 | yes |
| 679 | 673 | Q9BZF1\|OSBL8_HUMAN | 41.39 | 2 | 1 | 1 | 101196 | OSBPL8 | no |
| 680 | 675 | P35606\|COPB2_HUMAN | 41.26 | 1 | 1 | 1 | 102487 | COPB2 | yes |
| 681 | 677 | Q14746\|COG2_HUMAN | 41.18 | 2 | 1 | 1 | 83208 | COG2 | no |
| 682 | 678 | O75323\|NIPS2_HUMAN | 41.17 | 3 | 1 | 1 | 33743 | NIPSNAP2 | no |
| 683 | 679 | C9J798\|RAS4B_HUMAN | 40.97 | 2 | 1 | 1 | 90406 | RASA4B | no |
| 683 | 680 | O43374\|RASL2_HUMAN | 40.97 | 2 | 1 | 1 | 90458 | RASA4 | no |
| 684 | 684 | P62491\|RB11A_HUMAN | 40.65 | 5 | 1 | 1 | 24394 | RAB11A | no |
| 684 | 685 | Q15907\|RB11B_HUMAN | 40.65 | 5 | 1 | 1 | 24488 | RAB11B | no |
| 685 | 686 | O60566\|BUB1B_HUMAN | 40.63 | 1 | 1 | 1 | 119545 | BUB1B | yes |
| 688 | 689 | Q96I24\|FUBP3_HUMAN | 40.34 | 2 | 1 | 1 | 61640 | FUBP3 | no |
| 688 | 690 | Q96AE4\|FUBP1_HUMAN | 40.34 | 1 | 1 | 1 | 67560 | FUBP1 | yes |
| 686 | 687 | P51149\|RAB7A_HUMAN | 40.33 | 6 | 1 | 1 | 23490 | RAB7A | yes |
| 687 | 688 | O94874\|UFL1_HUMAN | 40.27 | 1 | 1 | 1 | 89595 | UFL1 | no |
| 689 | 691 | P19404\|NDUV2_HUMAN | 40.2 | 4 | 1 | 1 | 27392 | NDUFV2 | no |
| 690 | 692 | Q14161\|GIT2_HUMAN | 39.75 | 1 | 1 | 1 | 84543 | GIT2 | yes |
| 691 | 693 | Q15813\|TBCE_HUMAN | 39.6 | 4 | 1 | 1 | 59346 | TBCE | yes |
| 692 | 414 | Q16204\|CCDC6_HUMAN | 39.51 | 4 | 1 | 1 | 53291 | CCDC6 | no |
| 693 | 694 | Q92552\|RT27_HUMAN | 39 | 5 | 1 | 1 | 47611 | MRPS27 | no |

| | | | | | | | | | |
|---|---|---|---|---|---|---|---|---|---|
| 493 | 395 | Q5T9A4\|ATD3B_HUMAN | 38.97 | 2 | 1 | 1 | 72573 | ATAD3B | yes |
| 694 | 696 | P46109\|CRKL_HUMAN | 38.63 | 4 | 1 | 1 | 33777 | CRKL | no |
| 695 | 698 | O15144\|ARPC2_HUMAN | 38.45 | 5 | 1 | 1 | 34333 | ARPC2 | no |
| 696 | 699 | Q04206\|TF65_HUMAN | 38.43 | 3 | 1 | 1 | 60219 | RELA | no |
| 697 | 700 | P57678\|GEMI4_HUMAN | 38.11 | 1 | 1 | 1 | 120037 | GEMIN4 | no |
| 417 | 380 | Q8N3C0\|ASCC3_HUMAN | 38.02 | 0 | 1 | 1 | 251458 | ASCC3 | no |
| 698 | 703 | P36957\|ODO2_HUMAN | 37.7 | 3 | 1 | 1 | 48755 | DLST | yes |
| 488 | 384 | Q13362\|2A5G_HUMAN | 37.66 | 3 | 1 | 1 | 61061 | PPP2R5C | no |
| 699 | 704 | Q14693\|LPIN1_HUMAN | 37.5 | 2 | 1 | 1 | 98664 | LPIN1 | no |
| 502 | 428 | Q99426\|TBCB_HUMAN | 37 | 9 | 1 | 1 | 27326 | TBCB | yes |
| 700 | 707 | P46459\|NSF_HUMAN | 36.91 | 2 | 1 | 1 | 82594 | NSF | no |
| 701 | 385 | Q99996\|AKAP9_HUMAN | 36.86 | 0 | 1 | 1 | 452990 | AKAP9 | yes |
| 486 | 381 | P49321\|NASP_HUMAN | 35.48 | 3 | 1 | 1 | 85238 | NASP | no |
| 703 | 712 | Q9C0C9\|UBE2O_HUMAN | 35.44 | 1 | 1 | 1 | 141293 | UBE2O | no |
| 704 | 714 | Q8N0Y7\|PGAM4_HUMAN | 35.37 | 6 | 1 | 1 | 28777 | PGAM4 | no |
| 704 | 715 | P18669\|PGAM1_HUMAN | 35.37 | 6 | 1 | 1 | 28804 | PGAM1 | no |
| 704 | 713 | P15259\|PGAM2_HUMAN | 35.37 | 6 | 1 | 1 | 28766 | PGAM2 | no |
| 516 | 702 | P21964\|COMT_HUMAN | 35.15 | 6 | 1 | 1 | 30037 | COMT | no |
| 706 | 717 | P09455\|RET1_HUMAN | 35.13 | 7 | 1 | 1 | 15850 | RBP1 | no |
| 705 | 716 | P55884\|EIF3B_HUMAN | 35.08 | 2 | 1 | 1 | 92482 | EIF3B | no |
| 302 | 408 | Q15436\|SC23A_HUMAN | 34.9 | 3 | 1 | 1 | 86161 | SEC23A | no |
| 386 | 722 | Q96Q15\|SMG1_HUMAN | 34.86 | 1 | 3 | 2 | 410501 | SMG1 | yes |
| 386 | 1613 | Q9NZJ4\|SACS_HUMAN | 20.05 | 0 | 1 | 0 | 521131 | SACS | no |
| 386 | 1902 | P33402\|GCYA2_HUMAN | 20.05 | 1 | 1 | 0 | 81750 | GUCY1A2 | no |
| 386 | 2032 | Q14241\|ELOA1_HUMAN | 20.05 | 1 | 1 | 0 | 89909 | ELOA | no |
| 386 | 2033 | Q6AI14\|SL9A4_HUMAN | 20.05 | 1 | 1 | 0 | 89814 | SLC9A4 | no |
| 386 | 2311 | Q8TDW7\|FAT3_HUMAN | 20.05 | 0 | 1 | 0 | 501983 | FAT3 | no |
| 386 | 2375 | A6NNT2\|CP096_HUMAN | 20.05 | 0 | 1 | 0 | 125041 | C16orf96 | no |
| 386 | 2432 | Q5VT97\|SYDE2_HUMAN | 20.05 | 0 | 1 | 0 | 133230 | SYDE2 | no |
| 386 | 2705 | Q6ZRR7\|LRRC9_HUMAN | 20.05 | 0 | 1 | 0 | 166910 | LRRC9 | no |
| 386 | 2751 | O14795\|UN13B_HUMAN | 20.05 | 0 | 1 | 0 | 180678 | UNC13B | no |
| 386 | 2757 | O00750\|P3C2B_HUMAN | 20.05 | 0 | 1 | 0 | 184767 | PIK3C2B | no |
| 386 | 2845 | Q9UPW8\|UN13A_HUMAN | 20.05 | 0 | 1 | 0 | 193013 | UNC13A | no |
| 386 | 1762 | Q8IVF4\|DYH10_HUMAN | 20.05 | 0 | 1 | 0 | 514845 | DNAH10 | no |

| | | | | | | | | | |
|---|---|---|---|---|---|---|---|---|---|
| 386 | 2864 | Q07283\|TRHY_HUMAN | 20.05 | 0 | 1 | 0 | 253922 | TCHH | no |
| 540 | 1214 | Q9Y5Q9\|TF3C3_HUMAN | 34.78 | 1 | 1 | 1 | 101272 | GTF3C3 | no |
| 384 | 818 | Q6T4R5\|NHS_HUMAN | 34.31 | 1 | 2 | 2 | 179134 | NHS | no |
| 707 | 719 | Q96EL3\|RM53_HUMAN | 33.26 | 12 | 1 | 1 | 12107 | MRPL53 | no |
| 708 | 720 | P07949\|RET_HUMAN | 33.25 | 1 | 1 | 1 | 124319 | RET | no |
| 709 | 723 | P60660\|MYL6_HUMAN | 32.64 | 11 | 1 | 1 | 16930 | MYL6 | no |
| 710 | 728 | Q9BPU6\|DPYL5_HUMAN | 32.15 | 3 | 1 | 1 | 61421 | DPYSL5 | no |
| 455 | 731 | Q9UKN7\|MYO15_HUMAN | 32.02 | 0 | 1 | 1 | 395294 | MYO15A | no |
| 711 | 730 | P56537\|IF6_HUMAN | 32.02 | 7 | 1 | 1 | 26599 | EIF6 | no |
| 711 | 729 | tr\|A0A0U1RQV5\|A0A0U1RQV5_HUMAN | 32.02 | 15 | 1 | 1 | 11832 | NA | no |
| 394 | 842 | Q13402\|MYO7A_HUMAN | 31.53 | 1 | 1 | 1 | 254388 | MYO7A | no |
| 311 | 708 | Q13283\|G3BP1_HUMAN | 31.52 | 4 | 1 | 1 | 52164 | G3BP1 | yes |
| 312 | 734 | P35637\|FUS_HUMAN | 30.28 | 3 | 1 | 1 | 53426 | FUS | yes |
| 518 | 737 | Q9H115\|SNAB_HUMAN | 30.26 | 4 | 1 | 1 | 33557 | NAPB | no |
| 313 | 738 | P81605\|DCD_HUMAN | 30.18 | 8 | 1 | 1 | 11284 | DCD | no |
| 421 | 822 | Q9P2D7\|DYH1_HUMAN | 29.38 | 0 | 1 | 1 | 487482 | DNAH1 | no |
| 755 | 1592 | P48681\|NEST_HUMAN | 29.07 | 1 | 1 | 1 | 177438 | NES | no |
| 713 | 746 | Q86UE4\|LYRIC_HUMAN | 28.95 | 5 | 1 | 1 | 63837 | MTDH | no |
| 716 | 757 | Q9NNW5\|WDR6_HUMAN | 28.77 | 1 | 1 | 1 | 121724 | WDR6 | no |
| 715 | 755 | P38919\|IF4A3_HUMAN | 28.51 | 2 | 1 | 1 | 46871 | EIF4A3 | no |
| 766 | 750 | Q8WZ42\|TITIN_HUMAN | 28.25 | 0 | 2 | 2 | 4E+06 | TTN | no |
| 717 | 758 | P82650\|RT22_HUMAN | 27.87 | 4 | 1 | 1 | 41280 | MRPS22 | no |
| 718 | 759 | O43237\|DC1L2_HUMAN | 27.86 | 3 | 1 | 1 | 54099 | DYNC1LI2 | no |
| 719 | 760 | P32322\|P5CR1_HUMAN | 27.83 | 5 | 1 | 1 | 33361 | PYCR1 | no |
| 519 | 739 | Q96GQ5\|RUS1_HUMAN | 27.48 | 2 | 1 | 1 | 51018 | C16orf58 | no |
| 721 | 770 | P52735\|VAV2_HUMAN | 26.86 | 2 | 1 | 1 | 101289 | VAV2 | yes |
| 722 | 773 | P67936\|TPM4_HUMAN | 26.83 | 8 | 1 | 1 | 28522 | TPM4 | no |
| 723 | 791 | Q5JPH6\|SYEM_HUMAN | 26.44 | 4 | 1 | 1 | 58689 | EARS2 | no |
| 724 | 793 | Q641Q2\|WAC2A_HUMAN | 26.37 | 3 | 1 | 1 | 147183 | WASHC2A | no |
| 724 | 792 | Q9Y4E1\|WAC2C_HUMAN | 26.37 | 3 | 1 | 1 | 144911 | WASHC2C | no |
| 451 | 415 | Q14318\|FKBP8_HUMAN | 25.85 | 4 | 1 | 1 | 44562 | FKBP8 | no |
| 725 | 800 | Q6NSJ5\|LRC8E_HUMAN | 25.83 | 2 | 1 | 1 | 90247 | LRRC8E | no |
| 727 | 865 | Q9H270\|VPS11_HUMAN | 24.9 | 1 | 1 | 1 | 107837 | VPS11 | no |
| 730 | 929 | Q8ND24\|RN214_HUMAN | 24.05 | 3 | 1 | 1 | 77667 | RNF214 | no |

| | | | | | | | | | |
|---|---|---|---|---|---|---|---|---|---|
| 423 | 932 | Q96RD1\|OR6C1_HUMAN | 23.63 | 3 | 1 | 1 | 35660 | OR6C1 | no |
| 731 | 934 | Q9BXX0\|EMIL2_HUMAN | 23.29 | 1 | 1 | 1 | 115687 | EMILIN2 | no |
| 733 | 936 | Q9NYB9\|ABI2_HUMAN | 23.09 | 4 | 1 | 1 | 55663 | ABI2 | no |
| 523 | 830 | Q12879\|NMDE1_HUMAN | 23 | 1 | 1 | 1 | 165282 | GRIN2A | no |
| 734 | 939 | P35573\|GDE_HUMAN | 22.91 | 2 | 1 | 1 | 174763 | AGL | no |
| 736 | 941 | A6NGY5\|O51F1_HUMAN | 22.9 | 3 | 1 | 1 | 35849 | OR51F1 | no |
| 398 | 724 | Q99460\|PSMD1_HUMAN | 22.87 | 3 | 1 | 1 | 105836 | PSMD1 | no |
| 400 | 826 | Q53RY4\|KCP3_HUMAN | 22.75 | 4 | 1 | 1 | 25627 | KRTCAP3 | no |
| 735 | 940 | Q14684\|RRP1B_HUMAN | 22.69 | 2 | 1 | 1 | 84428 | RRP1B | no |
| 547 | 1704 | Q5T7N3\|KANK4_HUMAN | 22.47 | 1 | 1 | 1 | 107341 | KANK4 | no |
| 738 | 944 | P61289\|PSME3_HUMAN | 22.41 | 6 | 1 | 1 | 29506 | PSME3 | no |
| 741 | 950 | Q8N0X4\|CLYBL_HUMAN | 22.18 | 6 | 1 | 1 | 37359 | CLYBL | no |
| 742 | 953 | Q9UEW8\|STK39_HUMAN | 21.62 | 4 | 1 | 1 | 59474 | STK39 | no |
| 422 | 908 | O75369\|FLNB_HUMAN | 21.62 | 0 | 1 | 1 | 278162 | FLNB | yes |
| 753 | 987 | Q5JSL3\|DOC11_HUMAN | 21.56 | 0 | 1 | 1 | 237669 | DOCK11 | no |
| 521 | 827 | Q2T9J0\|TYSD1_HUMAN | 21.49 | 2 | 1 | 1 | 59309 | TYSND1 | no |
| 743 | 956 | P49590\|SYHM_HUMAN | 21.46 | 4 | 1 | 1 | 56888 | HARS2 | no |
| 462 | 876 | Q9Y4A5\|TRRAP_HUMAN | 21.36 | 0 | 1 | 1 | 437603 | TRRAP | no |
| 462 | 880 | Q5T1H1\|EYS_HUMAN | 21.36 | 0 | 1 | 1 | 350797 | EYS | no |
| 744 | 958 | C9JH25\|PRRT4_HUMAN | 21.31 | 2 | 1 | 1 | 92712 | PRRT4 | no |
| 745 | 960 | Q13637\|RAB32_HUMAN | 21.24 | 5 | 1 | 1 | 24997 | RAB32 | no |
| 539 | 1139 | O94887\|FARP2_HUMAN | 20.99 | 1 | 1 | 1 | 119888 | FARP2 | no |
| 748 | 967 | Q8TDU6\|GPBAR_HUMAN | 20.98 | 5 | 1 | 1 | 35248 | GPBAR1 | no |
| 556 | 2309 | Q96NH3\|BROMI_HUMAN | 20.78 | 1 | 1 | 1 | 144755 | TBC1D32 | no |
| 531 | 973 | Q70YC5\|ZN365_HUMAN | 20.57 | 2 | 1 | 1 | 46542 | ZNF365 | no |
| 750 | 974 | P69849\|NOMO3_HUMAN | 20.47 | 1 | 1 | 1 | 134133 | NOMO3 | no |
| 750 | 975 | Q15155\|NOMO1_HUMAN | 20.47 | 1 | 1 | 1 | 134324 | NOMO1 | no |
| 750 | 976 | Q5JPE7\|NOMO2_HUMAN | 20.47 | 1 | 1 | 1 | 139439 | NOMO2 | no |
| 752 | 979 | Q5T6F2\|UBAP2_HUMAN | 20.34 | 1 | 1 | 1 | 117115 | UBAP2 | no |
| 525 | 802 | Q7Z7M0\|MEGF8_HUMAN | 20.31 | 0 | 1 | 1 | 303100 | MEGF8 | no |
